# A data-informed mean-field approach to mapping of cortical parameter landscapes

**DOI:** 10.1101/2021.10.23.465568

**Authors:** Zhuo-Cheng Xiao, Kevin K. Lin, Lai-Sang Young

## Abstract

Constraining the many biological parameters that govern cortical dynamics is computationally and conceptually difficult because of the curse of dimensionality. This paper addresses these challenges by proposing (1) a novel data-informed mean-field (MF) approach to efficiently map the parameter space of network models; and (2) an organizing principle for studying parameter space that enables the extraction biologically meaningful relations from this high-dimensional data. We illustrate these ideas using a large-scale network model of the *Macaque* primary visual cortex. Of the 10-20 model parameters, we identify 7 that are especially poorly constrained, and use the MF algorithm in (1) to discover the firing rate contours in this 7D parameter cube. Defining a “biologically plausible” region to consist of parameters that exhibit spontaneous Excitatory and Inhibitory firing rates compatible with experimental values, we find that this region is a slightly thickened codimension-1 submanifold. An implication of this finding is that while plausible regimes depend sensitively on parameters, they are also robust and flexible provided one compensates appropriately when parameters are varied. Our organizing principle for conceptualizing parameter dependence is to focus on certain 2D parameter planes that govern lateral inhibition: Intersecting these planes with the biologically plausible region leads to very simple geometric structures which, when suitably scaled, have a universal character independent of where the intersections are taken. In addition to elucidating the geometry of the plausible region, this invariance suggests useful approximate scaling relations. Our study offers, for the first time, a complete characterization of the set of all biologically plausible parameters for a detailed cortical model, which has been out of reach due to the high dimensionality of parameter space.

**Author Summary:** Cortical circuits are characterized by a high degree of structural and dynamical complexity, and this biological reality is reflected in the large number of parameters in even semi-realistic cortical models. A fundamental task of computational neuroscience is to understand how these parameters govern network dynamics. While some neuronal parameters can be measured *in vivo*, many remain poorly constrained due to limitations of available experimental techniques. Computational models can address this problem by relating difficult-to-measure parameters to observable quantities, but to do so one must overcome two challenges: (1) the computational expense of mapping a high dimensional parameter space, and (2) extracting biological insights from such a map. This study aims to address these challenges in the following ways: First, we propose a parsimonious data-informed algorithm that efficiently predicts spontaneous cortical activity, thereby speeding up the mapping of parameter landscapes. Second, we show that lateral inhibition provides a basis for conceptualizing cortical parameter space, enabling us to begin to make sense of its geometric structure and attendant scaling relations. We illustrate our approach on a biologically realistic model of the monkey primary visual cortex.

## Introduction

From spatially and temporally homogeneous but sensitive resting states to highly structured evoked responses, neuronal circuits in the cerebral cortex exhibit an extremely broad range of dynamics in support of information processing in the brain [1–8]. Accompanying this dynamical flexibility is a high degree of morphological and physiological complexity [9–15]. As a result, any effort to characterize cortical circuits necessarily involves a large number of biological parameters [16–21]. Understanding the range of parameters compatible with biologically plausible cortical dynamics and how individual parameters impact neural computation are, in our view, basic questions in computational neuroscience.

Due to limitations of available experimental techniques, many neuronal and network parameters are poorly constrained. Biologically realistic network models can bridge this gap by quantifying the dependence of observable quantities like firing rates on parameters, thereby constraining their values. However, two challenges stand in the way of efforts to map the parameter landscape of detailed cortical networks. First, a direct approach, i.e., parameter sweeps using network models, may be extremely costly or even infeasible. This is because even a single layer of a small piece of cortex consists of tens of thousands of neurons, and the computational cost grows rapidly with the size of the network. This cost is compounded by the need for repeated model runs during parameter sweeps, and by the “curse of dimensionality,” i.e., the exponential growth of parameter space volume with the number of parameters. Second, even after conducting parameter sweeps, one is still faced with the daunting task of making sense of the high dimensional data to identify *interpretable, biologically meaningful features.*

This paper addresses the twin challenges of computational cost and interpretable cortical parameter mapping. Starting from a biologically realistic network model, we define as “viable” those parameters that yield predictions compatible with empirically observed firing rates, and seek to identify the viable region. To mitigate the computational cost of parameter space scans, we propose a parsimonious, data-informed mean-field (MF) algorithm. MF methods replace rapidly-fluctuating quantities like membrane potentials with their mean values; they have been used a great deal in neuroscience [22–38]. MF models of neuronal networks are all based on the relevant biology to different degrees, but most rely on idealized voltage-rate relations (so-called “gain” or “activation” function) to make the system amenable to analysis (see, e.g., [39, 40] and the Discussion for more details). In contrast, our MF equations are derived from a biologically realistic network model: Instead of making assumptions on gain functions, our MF equations follow closely the anatomical and physiological information incorporated in the network, hence reflecting its key features. To stress this tight connection to a realistic network model, we have described our method as a “*data-informed* MF approach”. As we will show, the algorithm we propose is capable of accurately predicting network firing rates at a small fraction of the expense of direct network simulations.

We illustrate the power of this approach and how one might conceptualize the mapping it produces using a biologically realistic model of Macaque primary visual cortex (V1). Focusing on spontaneous activity, our main result is that the viable region is a thin neighborhood of a *codimension-1 manifold* in parameter space. (A codimension-1 manifold is an (*n* – 1)-dimensional surface in an *n*-dimensional parameter space.) Being approximately codimension-1 implies that the viable region is simultaneously sensitive and flexible to parameter changes: sensitive in that a small perturbation can easily move a point off the manifold; flexible in the sense that it allows for a great variety of parameter combinations, consistent with the wide variability observed in biology. Our analysis of parameter dependence is based on the following organizing principle: By restricting attention to certain 2D planes associated with lateral inhibition, we discover geometric structures that are remarkably similar across all such “inhibition planes”. Our findings suggest a number of simple *approximate scaling relations* among neuronal parameters.

The Macaque V1 network model we use for illustration involves ≳ 4 × 10^4^ neurons in Layer 4C*α* of V1, and has been carefully benchmarked to a large number of known features of V1 response [41]. In [41], the authors focused on a small parameter region which they had reason to believe to be viable. The present study produces a much more comprehensive characterization of the set of all viable parameters defined in terms of spontaneous activity. The reason we have focused on spontaneous activity is that it is a relatively simple, homogeneous equilibrium steady state, and understanding it is necessary before tackling more complex, evoked responses. However, as all cortical activity depends on a delicate balance between excitation and inhibition, even background dynamics can be rather nontrivial.

Parameter search and tuning are problems common to all areas of computational biology. By significantly reducing the cost of mapping parameter landscapes, we hope the computational strategy proposed in the present paper will enable computational neuroscientists to construct high-fidelity cortical models, and to use these models to shed light on spontaneous and evoked dynamics in neural circuitry. Moreover, reduced models of the type proposed here may be useful as a basis for parameter and state estimation on the basis of experimental data.

## Results

As explained in the Introduction, this work (1) proposes a novel data-informed mean-field approach to facilitate efficient and systematic parameter analysis of neuronal networks, which we validate using a previously constructed model of the monkey visual cortex; and (2) we develop ways to conceptualize and navigate the complexities of high-dimensional parameter spaces of neuronal models by organizing around certain relationships among parameters, notably those governing lateral inhibition.

Sect. 1 describes the network model of the visual cortex that will be used both to challenge the MF algorithm and to assess its efficacy, together with a brief introduction to the algorithm itself; details are given in **Methods**. Sect. 2 uses the algorithm to explore the parameter landscape of the model. Qualitative analysis is offered along the way leading to a conceptual understanding of parameter dependence.

### 1 Network model and parameter landscape

We use as starting point the large-scale network model in [41]. This is a mechanistic model of an input layer to the primary visual cortex (V1) of the Macaque monkey, which has vision very similar to that of humans [42–46] Among existing neuronal network models, [41] is at the very high end on the scale of details and biological realism: It incorporates a good amount of known neuroanatomy in its network architecture, capturing the dynamics of individual neurons as well as their dynamical interaction.

In [41], the authors located a small set of potentially viable parameters, which they refined by benchmarking the resulting regimes against multiple sets of experimental data. No claims were made on the uniqueness or optimality of the parameters considered. Indeed, because of the intensity of the work involved in locating viable parameters, little attempt was made to consider parameters farther from the ones used. This offers a natural testing ground for our novel approach to parameter determination: We borrow certain aspects of the model from [41], including network architecture, equations of neuronal dynamics and parameter structure, but instead of using information on the parameters found, we will search for viable parameter regions using the techniques developed here.

For a set of parameters to be viable, it must produce firing rates similar to those of the real cortex, including *background* firing, the spontaneous spiking produced when cortex is unstimulated. Background activity provides a natural way to constrain parameters: It is an especially simple state of equilibrium, one in which spiking is statistically homogeneous in space and time and involves fewer features of cortical dynamics. For example, synaptic depression and facilitation are not known to play essential roles in spontaneous activity. A goal in this paper will be to systematically identify all regions in parameter space with acceptable background firing rates.

#### 1.1 Network model of an input layer to primate visual cortex

The network model is that of a small patch of Macaque V1, layer 4C*α* (hereafter “L4”), the input layer of the Magnocellular pathway. This layer receives input from the lateral geniculate nucleus (LGN) and feedback from layer 6 (L6) of V1. Model details are as in [41], except for omission of features not involved in background activity. We provide below a model description that is sufficient for understanding – at least on a conceptual level – the material in **Results**. Precise numerical values of the various quantities are given in **SI**. For more detailed discussions of the neurobiology behind the material in this subsection, we refer interested readers to [41] and references therein.

##### Network architecture

Of primary interest to us is L4, which is modeled as a 2D network of point neurons. Locations within this layer are identified with locations in the retina via neuronal projections, and distances on the retina are measured in degrees. Cells^1^ are organized into hypercolumns of about 4000 neurons each, covering a 0.25° × 0.25° area. The neurons are assumed to be of two kinds: 75 – 80% are Excitatory (E), and the rest are Inhibitory (I). The E-population is evenly placed in a lattice on the cortical surface; the same is true for I-cells. Projections to E- and I-cells are assumed to be isotropic, with probabilities of connection described by truncated Gaussians. E-neurons (which are assumed to be spiny stellate cells) have longer axons, about twice that of I-neurons (which are assumed to be basket cells). E-to-E coupling is relatively sparse, at about 15% at the peak, while E-to-I, I-to-E and I-to-I coupling is denser, at about 60%. Connections are drawn randomly subject to the probabilities above. On average, each E-neuron receives synaptic input from slightly over 200 E-cells and about 100 I-cells, while each I-cell receives input from ~ 800 E-cells and 100 I-cells. Exact numbers are given in **SI**.

Cells in L4 also receive synaptic input from two external sources, “external” in the sense that they originate from outside of this layer. One source is LGN: Each L4 cell, E or I, is assumed to be connected to 4 LGN cells on average; each LGN cell is assumed to provide 20 spikes per sec, a typical firing rate in background. These spikes are assumed to be delivered to L4 cells in a Poissonian manner, independently from cell to cell. L6 provides another source of synaptic input: We assume each E-cell in L4 receives input from 50 E-cells from L6, consistent with known neurobiology, and that each L6 E-cell fires on average 5 spikes per sec in background. For I-cells, the number of presynaptic L6 cells is unknown; this is one of the free parameters we will explore. Spike times from L6 are also assumed to be Poissonian and independent from cell to cell, a slight simplification of [41]. All inputs from external sources are excitatory. Finally, we lump together all top-down modulatory influences on L4 not modeled into a quantity we call “ambient”. Again, see **SI** for all pertinent details.

##### Equations of neuronal dynamics

We model only the dynamics of neurons in L4. Each neuron is modeled as a conductance-based point neuron whose membrane potential *v* evolves according to the leaky integrate-and-fire (LIF) equation [47, 48]

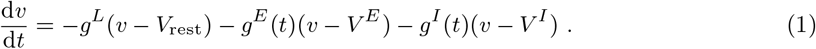

Following [41], we nondimensionalize *v*, with resting potential *V*_rest_ = 0 and spiking threshold *V*_th_ = 1. In Eq. (1), *v* is driven toward *V*th by the Excitatory current *g^E^*(*t*)(*v* – *V^E^*), and away from it by the leak term *g^L^v* and the Inhibitory current *g^I^*(*t*)(*v* – *V^I^*). When v reaches 1, a spike is fired, and *v* is immediately reset to 0, where it is held for a refractory period of 2 ms. The membrane leakage times 1/*g^L^* = 20 ms for E-neurons and 16.7 ms for I-neurons, as well as the reversal potentials *V^E^* = 14/3 and *V^I^* = −2/3, are standard [49].

The quantities *g^E^*(*t*) and *g^I^*(*t*), the Excitatory and Inhibitory conductances, are defined as follows. First,

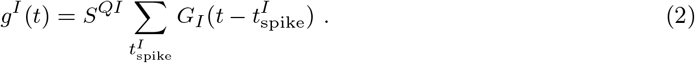

Here, the neuron whose dynamics are described in Eq. (1) is assumed to be of type *Q, Q* = E or I, and the constant *S^QI^* is the synaptic coupling weight from I-neurons to neurons of type *Q*. The summation is taken over 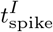, times at which a spike from a presynaptic I-cell from within layer 4C*α* is received by the neuron in question. Upon the arrival of each spike, *g_I_*(*t*) is elevated for 5-10 ms and *G_I_*(·) describes the waveform in the IPSC. Second, the Excitatory conductance *_gE_*(*t*) is the sum of 4 terms, the first three of which are analogs of the right side of Eq. (2), with an EPSC lasting 3-5 ms: they represent synaptic input from E-cells from within L4, from LGN and from L6. The 4th term is from ambient.

This completes the main features of the network model. Details are given in **SI**.

#### 1.2 Parameter space to be explored

Network dynamics can be very sensitive – or relatively robust – to parameter changes, and dynamic regimes can change differently depending on which parameter (or combination of parameters) is varied. To demonstrate the multiscale and anisotropic nature of the parameter landscape, we study the effects of parameter perturbations on L4 firing rates, using as reference point a set of biologically realistic parameters [41]. Specifically, we denote L4 E/I-firing rates at the reference point by 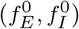, and fix a region 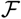 around 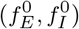 consisting of firing rates (*f_E_, f_I_*) we are willing to tolerate. We then vary network parameters one at a time, changing it in small steps and computing network firing rates 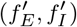 until they reach the boundary of 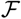, thereby determining the minimum perturbations needed to force L4 firing rates out of 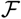.

Table 1 shows the results. We categorize the parameters according to the aspect of network dynamics they govern. As can be seen, L4 firing rates show varying degrees of sensitivity to perturbations in different parameter groups. They are most sensitive to perturbations to synaptic coupling weights within L4, where deviations as small as 1% can push the firing rates out of 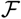. This likely reflects the delicate balance between excitation and inhibition, as well as the fact that the bulk of the synaptic input to a L4 neuron comes from lateral interaction, facts consistent with earlier findings [41]. With respect to parameters governing inputs from external sources, we find perturbing LGN parameters to have the most impact, followed by amb and L6, consistent with their net influence on *g_E,I_* in background. We also note that the parameters governing afferents to I cells are more tolerant of perturbations than those for E cells.

**Table 1:** Network parameters and response. Using the parameters from [41] as a reference point, we set 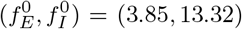 Hz and 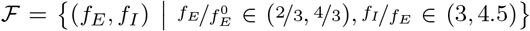 and vary one parameter at a time. We then compute the minimum perturbation needed to force the network firing rates out of 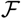. Values such as *F*^*E*P^, where *P* ∈ {lgn, L6, amb}, represent the total number of spikes per second received by an E-cell in L4 from source P. For example, in the reference set, each L4 cell has 4 afferent LGN cells on average, the mean firing rate of each is assumed to be 20 spikes/s, so *F*^*E*lgn^ = 80 Hz. Column 5 gives the lower and upper bounds of single-parameter variation (rounded to the nearest 1%) from the reference point that yield firing rates within 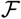.

| Group | Parameter | Meaning | Value | Bounds |
| --- | --- | --- | --- | --- |
| within L4 | $S^{EE}$ | E-to-E synaptic weight | 0.024 | (-3%, 1%) |
| | $S^{II}$ | I-to-I synaptic weight | 0.120 | (-4%, 1%) |
| | $S^{EI}$ | I-to-E synaptic weight | 0.0362 | (-1%, 3%) |
| | $S^{IE}$ | E-to-I synaptic weight | 0.0176 | (-1%, 3%) |
| LGN to L4 | $S^{Elgn}$ | lgn-to-E synaptic weight | 0.048 | (-5%, 3%) |
| | $S^{Ilgn}$ | lgn-to-I synaptic weight | 0.096 | (-6%, 9%) |
| | $F^{Elgn}$ | total # lgn spikes/s to E | 80 Hz | (-7%, 4%) |
| | $F^{Ilgn}$ | total # lgn spikes/s to I | 80 Hz | (-9%, 11%) |
| L6 to L4 | $S^{EL6}$ | L6-to-E synaptic weight | 0.008 | (-16%, 11%) |
| | $S^{IL6}$ | L6-to-I synaptic weight | 0.0058 | (-19%, 30%) |
| | $F^{EL6}$ | total # L6 spikes/s to E | 250 Hz | (-16%, 10%) |
| | $F^{IL6}$ | total # L6 spikes/s to I | 750 Hz | (-16%, 29%) |
| amb to L4 | $S^{amb}$ | ambient-to-E/I synaptic wt. | 0.01 | (-8%, 6%) |
| | $F^{Eamb}$ | rate of ambient to E | 500 Hz | (-7%, 5%) |
| | $F^{Iamb}$ | rate of ambient to I | 500 Hz | (-10%, 27%) |

These observations suggest that network dynamics depend in a complex and subtle way on parameters; they underscore the challenges one faces when attempting to tune parameters “by hand.” We now identify the parameters in the network model description in Sect. 1.1 to be treated as *free parameters* in the study to follow.

##### Free parameters

We consider a parameter “free” if it is hard to measure (or has not yet been measured) directly in the laboratory, or when data offer conflicting guidance. When available data are sufficient to confidently associate a value to a parameter, we consider it fixed. Following this principle, we designate the following 6 synaptic coupling weights governing recurrent interactions within L4 and its thalamic inputs as free parameters:

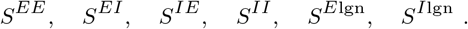

As shown in Table 1, these are also the parameters to which network response rates are the most sensitive. As for *S*^*E*L6^ and *S*^*I*L6^, which govern synaptic coupling from L6 to E- and I-neurons in L4, we assume

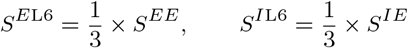

following [50] (see also [41]). This means that in our study, these quantities will vary, but they are indexed to *S^EE^* and *S^IE^* in a fixed manner and we will not regard them as free parameters.

A second category of parameters govern external sources. Here we regard *F*^*E*lgn^, *F*^*I*lgn^ and *F*^*E*L6^ as fixed to the values given in Table 1. L6 firing rates in background have been measured, but we know of no estimates on the number of presynaptic L6 cells to I-cells, so we treat *F*^*I*L6^ (which combines the effects of both) as a free parameter. The relation between *S*^*I*L6^ and *S^IE^* assumed above is in fact unknown from experiments. On the other hand, we are assuming that errors in the estimate of *S*^*I*L6^ can be absorbed into *F*^*I*L6^, which we vary. As for “ambient”, these inputs are thought to be significant, though not enough to drive spikes on their own. Since so little is known about this category of inputs, we fix the values of *S*^amb^, *F*^*E*amb^, and *F*^*I*amb^ to those given in Table 1, having checked that they meet the conditions above.

As discussed earlier, we are interested in L4 firing rates under background conditions. Denoting E- and I-firing rates by *f_E_* and *f_I_* respectively, the aim of our study can be summarized as follows:

###### Aim

*To produce maps of fE and fi as functions of the 7 parameters*

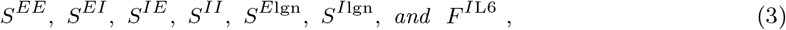

*to identify biologically relevant regions, and to provide a conceptual understanding of the results.*

#### 1.3 A brief introduction to our proposed MF approach

The approach we take is a MF computation of firing rates augmented by synthetic voltage data, a scheme we will refer to as “MF+v”. To motivate the “+v” part of the scheme, we first write down the MF equations obtained from Eq. (1) by balancing mean membrane currents. These MF equations will turn out to be incomplete. We discuss briefly how to secure the missing information; details are given in **Methods**.

##### MF equations

Eq. (1) reflects instantaneous current balance across the cell membrane of a L4 neuron. Assuming that this neuron’s firing rate coincides with that of the L4 E/I-population to which it belongs and neglecting (for now) refractory periods, we obtain a general relation between firing rates and mean currents by integrating Eq. (1). We will refer to the equations below as our “MF equations”. They have the general form

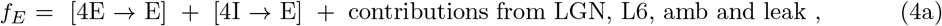

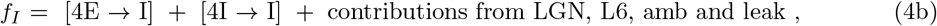

where [4E → E] represents the integral of the current contribution from E-cells in L4 to E-cells in L4, [4I → E] represents the corresponding quantity from I-cells in L4 to E-cells in L4, and so on. The contribution from lateral, intralaminar interactions can be further decomposed into, e.g.,

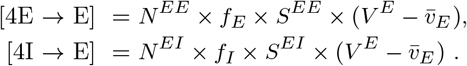

Here *N^EE^* and *N^EI^* are the mean numbers of presynaptic E- and I-cells from within L4 to an Eneuron, *f_E_, f_I_, S^EE^* and *S^EI^* are as defined earlier, and 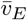 is the mean membrane potential *v* among E-neurons in L4. Other terms in Eq. (4a) and in Eq. (4b) are defined similarly; detailed derivation of the MF equations is given in **Methods**.

Network connectivity and parameters that are not considered “free parameters” are assumed to be fixed throughout. If additionally we fix a set of the 7 free parameters in (3), then Eq. (4) is linear in *f_E_* and *f_I_*, and are easily solved — except for two undetermined quantities, 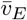 and 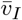. For network neurons, 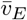 and 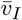 are emergent quantities that cannot be easily estimated from the equations of evolution or parameters chosen.

##### Estimating mean voltages

We explain here the ideas that lead to the algorithm we use for determining 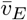 and 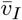, leaving technical details to **Methods**.

Our first observation is that the values of *f_E_* and *f_I_* computed from Eq. (4) depend delicately on 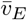 and 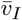; they can vary wildly with small changes in 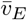 and 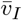. This ruled out the use of (guessed) approximate values, and even called into question the usefulness of the MF equations. But as we demonstrate in **Methods**, if one collects mean voltages 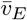 and 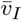 from network simulations and plug them into Eq. (4) to solve for *f_E_* and *f_I_*, then one obtains results that agree very well with actual network firing rates. This suggests Eq. (4) can be useful, provided we correctly estimate 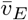 and 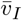.

As it defeats the purpose of an MF approach to use network simulations to determine 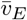 and 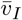, we sought to use a pair of LIF-neurons, one E and one I, to provide this information. To do that, we must create an environment for this pair of neurons that is similar to that within the network, incorporating the biological features with the LIF neurons. For example, one must use the same parameters and give them the same external drives, i.e., LGN, L6, and ambient. But a good fraction of the synaptic input to neurons in L4 are generated from lateral interactions; to simulate *that*, we would have to first learn what *f_E_* and *f_I_* are. The problem has now come full circle: what we need are *self-consistent* values of *f_E_* and *f_I_* for the LIF-neurons, so that their input and output firing rates coincide.

These and other ideas to be explained (e.g., efficiency and stability) go into the algorithm proposed. In a nutshell, we use the aid of a pair of LIF-neurons to help tie down 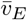 and 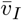, and use the MF equations to compute *f_E_* and *f_I_*. This mean-field algorithm aided by voltage closures (MF+v) is discussed in detail in **Methods**. We present next firing rate plots generated using this algorithm.

#### 2 Dependence of firing rates on system parameters

Even with a fast algorithm, so that many data points can be computed, discovery and representation of functions depending on more than 3 or 4 variables can be a challenge, not to mention conceptualization of the results. In Sects. 2.1–2.3, we propose to organize the 7D parameter space described in Sect. 1.2 in ways that take advantage of insights on how the parameters interact: Instead of attempting to compute 6D level surfaces for *f_E_* and *f_I_* embedded in the 7D parameter space, we identify a biologically plausible region of parameters called the “viable region”, and propose to study parameter structures by slicing the 7D space with certain 2D planes called “inhibition planes”. We will show that intersections of the viable region and inhibition planes – called “good areas” – possess certain canonical geometric structures, and that these structures offer a biologically interpretable landscape of parameter dependence. The three terms, *viable regions*, *inhibition planes* and *good areas*, the precise definitions of which are given in Sect. 2.1, are objects of interest throughout this section. In Sect. 2.4 we show comparison of firing rate computations from our algorithm and from actual network simulations.

##### 2.1 Canonical structures in inhibition planes

We have found it revealing to slice the parameter space using 2D planes defined by varying the parameters governing lateral inhibition, *S^IE^* and *S^EI^*, with all other parameters fixed. As we will show, these planes contain very simple and stable geometric structures around which we will organize our thinking about parameter space. Fig. 1A shows one such 2D slice. We computed raw contour curves for *f_E_* and *f_I_* on a 480 × 480 grid using the MF+v algorithm, with red curves for *f_E_* and blue for *f_I_*.

**Fig. 1:**
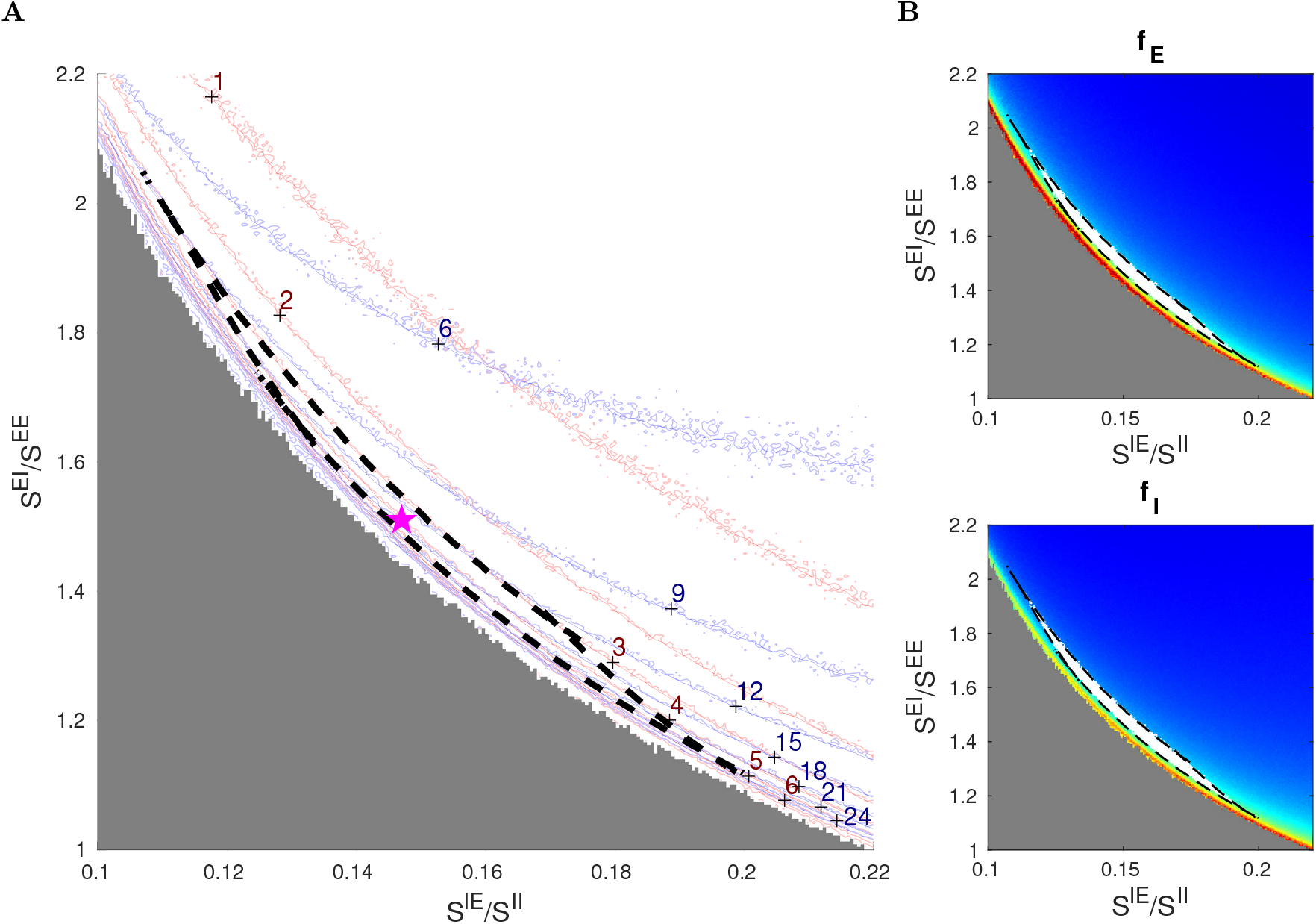
Canonical structures in an inhibition plane. **A.** Firing rates contours computed from firing rate maps on a 480×480 grid, showing *f_E_* = 1 – 6 Hz (red) and *f_I_* = 6 – 24 Hz (blue). The good area is indicated by the black dash lines (*f_E_* ∈ (3, 5) Hz and *f_I_*/*f_E_* ∈ (3,4.25)); the reference point is indicated by the purple star (⋆). The MF+v method becomes unstable and fails when the inhibitory index SI_*E*_ is too low (the gray region). **B.** Firing rate maps of *f_E_* (upper) and *f_I_* (lower), in which the good areas are indicated by white bands surrounded by black dash lines. The other five free parameters are as in Table 1: *S^EE^* = 0.024, *S^II^* = 0.12, *S*^*E*lgn^ = 2 × *S^EE^*, *S*^*I*lgn^ = 2 × *S*^*E*lgn^, and *F*^*I*L6^ = 3 × *F*^*E*L6^.

A striking feature of Fig. 1A is that the level curves are roughly hyperbolic in shape. We argue that this is necessarily so. First, note that in Fig. 1A, we used *S^IE^/S^II^* as *x*-axis and *S^EI^/S^EE^* as *y*-axis. The reason for this choice is that *S^IE^/S^II^* can be viewed as the degree of cortical excitation of I-cells, and *S^EI^/S^EE^* the suppressive power of I-cells from the perspective of E-cells. The product

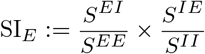

can therefore be seen as a *suppression index* for E-cells: the larger this quantity, the smaller *f_E_*. This suggests that the contours for *f_E_* should be of the form *xy* = constant, i.e., they should have the shape of hyperbolas. As E and I firing rates in local populations are known to covary, these approximately hyperbolic shapes are passed to contours of *f_I_*.

A second feature of Fig. 1A is that *f_I_* contours are less steep than those of *f_E_* at lower firing rates. That I-firing covaries with E-firing is due in part to the fact that I-cells receive a large portion of their excitatory input from E-cells through lateral interaction, at least when E-firing is robust. When *f_E_* is low, *f_I_* is lowered as well as I-cells lose their supply of excitation from E-cells, but the drop is less severe as I-cells also receive excitatory input from external sources. This causes *f_I_* contours to bend upwards relative to the *f_E_*-hyperbolas at lower firing rates, a fact quite evident from Fig. 1.

We now define the *viable region*, the biologically plausible region in our 7D parameter space, consisting of parameters that produce firing rates we deem close enough to experimentally observed values. For definiteness, we take these to be [51]

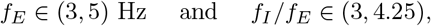

and refer to the intersection of the viable region with the 2D slice depicted in Fig. 1A as the “good area”. Here, the good area is the crescent-shaped set bordered by dashed black lines. For the parameters in Fig. 1, it is bordered by 4 curves, two corresponding to the *f_E_* = 3 and 5 Hz contours and the other two are where *f_I_/f_E_* = 3 and 4.25. That such an area should exist as a narrow strip of finite length, with unions of segments of hyperbolas as boundaries, is a consequence of the fact that *f_E_* and *f_I_*-contours are roughly but not exactly parallel. Fig. 1B shows the good area (white) on firing rate maps for *f_E_* and *f_I_*.

Hereafter, we will refer to 2D planes parametrized by *S^IE^/S^II^* and *S^EI^/S^EE^*, with all parameters other than *S^IE^* and *S^EI^* fixed, as *inhibition planes*, and will proceed to investigate the entire parameter space through these 2D slices and the good areas they contain. Though far from guaranteed, our aim is to show that the structures in Fig. 1A persist, and to describe how they vary with the other 5 parameters.

Fig. 1 is for a particular set of parameters. We presented it in high resolution to show our computational capacity and to familiarize the reader with the picture. As we vary parameters in the rest of this paper, we will present only heat maps for *f_E_* for each set of parameters studied. The good area, if there is one, will be marked in white, in analogy with the top panel of Fig. 1B.

Finally, we remark that the MF+v algorithm does not always return reasonable estimates of L4 firing rates. MF+v tends to fail especially for a low suppression index (gray area in Fig. 1A), where the network simulation also exhibits explosive, biologically unrealistic dynamics. This issue is discussed in **Methods** and **SI**.

##### 2.2 Dependence on external drives

There are two main sources of external input, LGN and L6 (while ambient input is assumed fixed). In both cases, it is their effect on E- *versus* I-cells in L4, and the variation thereof as LGN and L6 inputs are varied, that is of interest here.

###### LGN-to-E *versus* LGN-to-I

Results from [50] suggest that the sizes of EPSPs from LGN are ~ 2× those from L4. Based on this, we consider the range *S*^*E*lgn^/*S^EE^* ∈ (1.5, 3.0) in our study. Also, data show that LGN produces somewhat larger EPSCs in I-cells than in E-cells [52], though the relative coupling weights to E and I-cells are not known. Here, we index *S*^*I*lgn^ to *S*^*E*lgn^, and consider *S*^*I*lgn^/*S*^*E*lgn^ ∈ (1.5, 3.0).

Fig. 2 shows a 3 × 4 matrix of 2D panels, each one of which is an inhibition plane (see Fig. 1). This is the language we will use here and in subsequent figures: we will refer to the rows and columns of the matrix of panels, while *x* and *y* are reserved for the axes for each smaller panel.

**Fig. 2:**
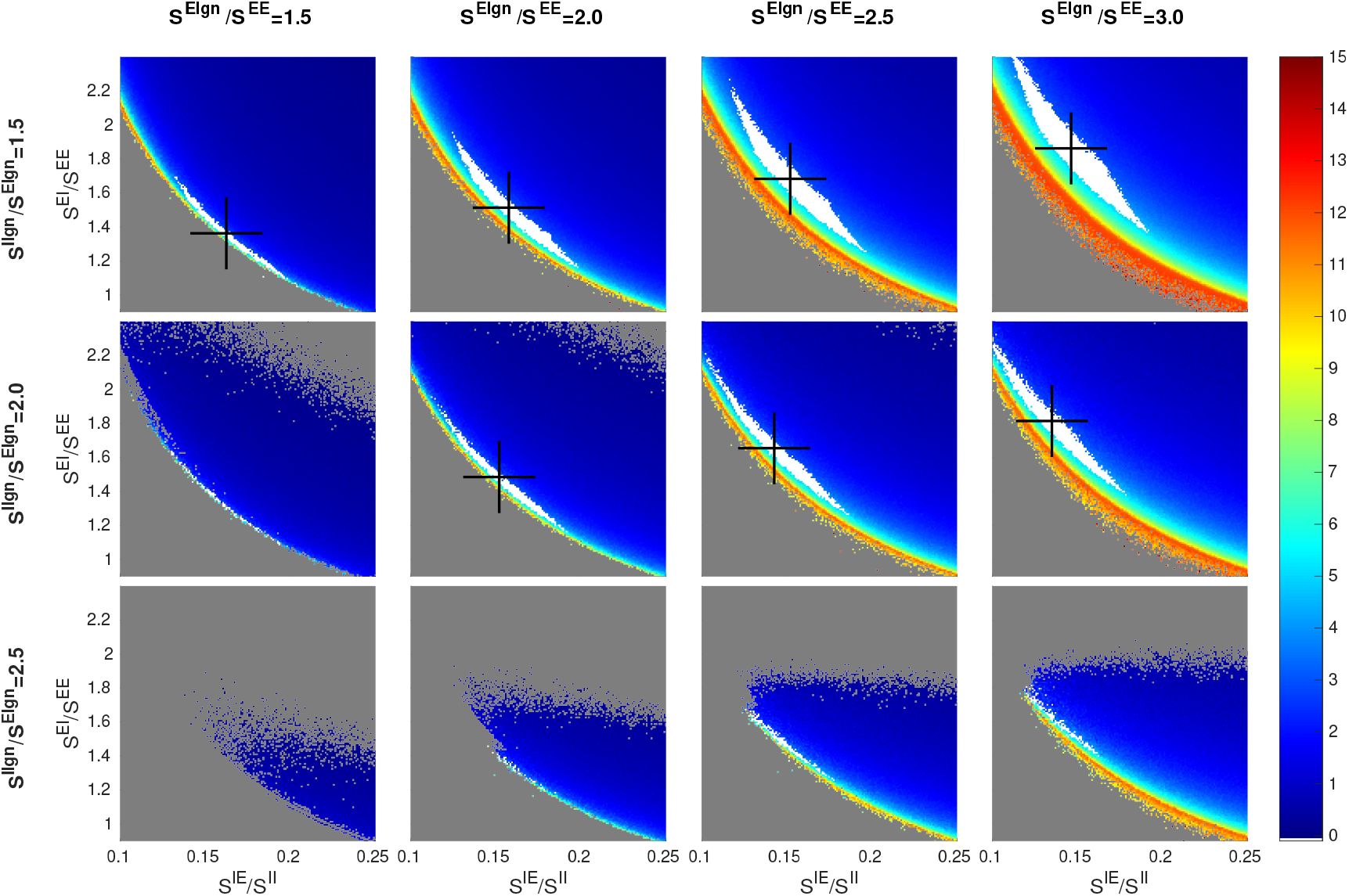
Dependence of firing rate and good area on LGN synaptic strengths. A 3×4 matrix of panels is shown: each row corresponds to a fixed value of *S*^*I*lgn^/*S*^*E*lgn^ and each column a fixed value of *S*^*E*lgn^/*S^EE^*. Each panel shows a heat map for *f_E_* on an inhibition plane (color bar on the right); *x* and *y*-axes are as in Fig. 1. Good areas are in white, and their centers of mass marked by black crosses. The picture for *S*^*E*lgn^/*S^EE^* = 2 and *S*^*I*lgn^/*S*^*E*lgn^ = 2 (row 2, column 2) corresponds to the *f_E_* rate map in Fig. 1. Other free parameters are *S^EE^* = 0.024, *S^II^* = 0.12, and *F*^*I*L6^ = 3 × *F*^*E*L6^.

We consider first the changes as we go from left to right in each row of the matrix in Fig. 2. With *S*^*I*lgn^/*S*^*E*lgn^ staying fixed, increasing *S*^*E*lgn^/*S^EE^* not only increases LGN’s input to E, but also increases LGN to I by the same proportion. It is evident that the rate maps in the subpanels are all qualitatively similar, but with gradual changes in the location and the shape of the good area. Specifically, as LGN input increases, (i) the center of mass of the good area (black cross) shifts upward and to the left following the hyperbola, and (ii) the white region becomes wider.

To understand these trends, it is important to keep in mind that the good area is characterized by having firing rates within a fairly narrow range. As LGN input to E increases, the amount of suppression must be increased commensurably to maintain E-firing rate. Within an inhibition plane, this means an increase in *S^EI^*, explaining the upward move of black crosses as we go from left to right. Likewise, the amount of E-to-I input must be decreased by a suitable amount to maintain I-firing at the same level, explaining the leftward move of the black crosses and completing the argument for (i). As for (ii), recall that the SI_*E*_ measures the degree of suppression of E-cells *from within L4* (and L6, the synaptic weights of which are indexed to those in L4). Increased LGN input causes E-cells to be less suppressed than their SI_*E*_ index would indicate. This has the effect of spreading the *f_E_* contours farther apart, stretching out the picture in a northeasterly direction perpendicular to the hyperbolas and widening the good area.

Going down the columns of the matrix, we observe a compression of the contours along the same northeasterly axis and a leftward shift of the black crosses. Recall that the only source of excitation of I-cells counted in the SI_*E*_-index is from L4 (hence also L6). When LGN to E is fixed and LGN to I is increased, the additional external drive to I produces a larger amount of suppression than the SI_*E*_-index would indicate, hence the compression. It also reduces the amount of *S^IE^* needed to produce the same I-firing rate, hence the leftward shift of the good area.

The changes in LGN input to E and to I shown cover nearly all of the biological range. We did not show the row corresponding to *S*^*I*lgn^/*S*^*E*lgn^ = 3 because the same trend continues and there are no viable regions. Notice that even though we have only shown a 3 × 4 matrix of panels, the trends are clear and one can easily interpolate between panels.

###### LGN-to-I *versus* L6-to-I

Next, we examine the relation between two sources of external drive: LGN and L6. In principle, this involves a total of 8 quantities: total number of spikes and coupling weights, from LGN and from L6, received by E and I-neurons in L4. As discussed in Sect. 1.2, enough is known about several of these quantities for us to treat them as “fixed”, leaving as free parameters the following three: *S*^*E*lgn^, *S*^*I*lgn^ and *F*^*I*L6^. Above, we focused on *S*^*I*lgn^/*S*^*E*lgn^, the impact of LGN on I relative to E. We now compare *S*^*I*lgn^/*S*^*E*lgn^ to *F*^*I*L6^/*F*^*E*L6^, the corresponding quantity with L6 in the place of LGN.

As there is little experimental guidance with regard to the range of *F*^*I*L6^, we will explore a relatively wide region of *F*^*I*L6^: Guided by the fact that

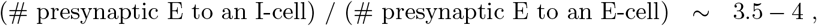

in L4 and the hypothesis that similar ratios hold for inter-laminar connections, we assume *F*^*I*L6^/*F*^*E*L6^ ∈ (1.5,6). We have broadened the interval because it is somewhat controversial whether the effect of L6 is net-Excitatory or net-Inhibitory: the modeling work [53] on monkey found that it had to be at least slightly net-Excitatory, while [54] reported that it was net-Inhibitory in mouse.

Fig. 3 shows, not surprisingly, that increasing LGN and L6 inputs to I have very similar effects: As with *S*^*I*lgn^/*S*^*E*lgn^, larger *F*^*I*L6^/*F*^*E*L6^ narrows the strip corresponding to the good area and shifts it leftwards, that is, going from left to right in the matrix of panels has a similar effect as going from top to bottom. Interpolating, one sees, e.g., that the picture at (*S*^*I*lgn^/*S*^*E*lgn^,*F*^*I*L6^/*F*^*E*L6^) = (1.5,4) is remarkably similar to that at (2, 1.5). In background, changing the relative strengths of LGN to I vs to E has a larger effect than the corresponding changes in L6, because LGN input is a larger component of the Excitatory input than L6. This relation may not hold under drive, however, where L6 response is known to increase significantly.

**Fig. 3:**
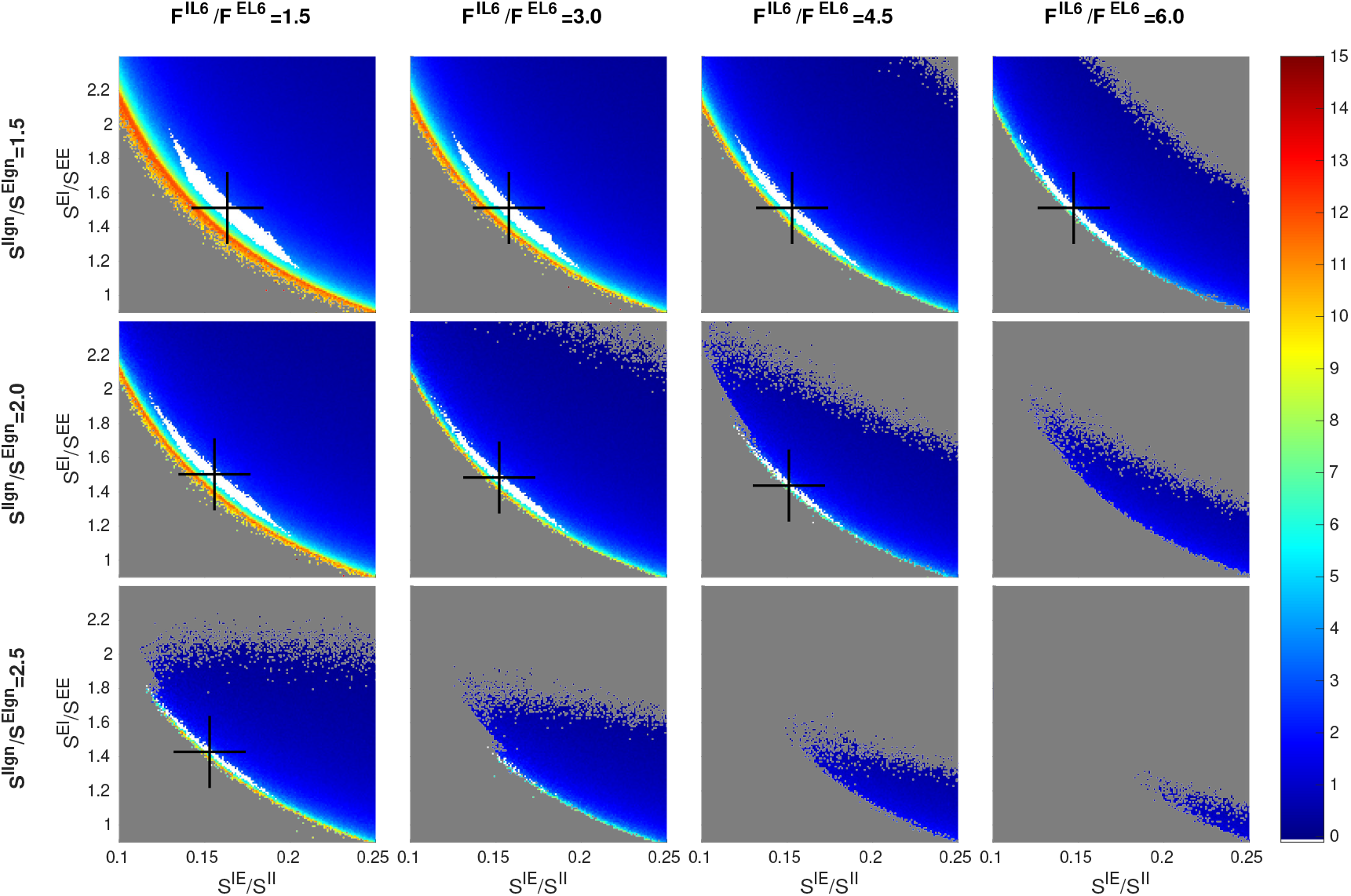
Dependence of firing rate and good area on LGN *versus* L6. A 3× 4 matrix of panels is shown: each row corresponds to a fixed value of *S*^*I*lgn^/*S*^*E*lgn^ and each column a fixed value of *F*^*I*L6^/*F*^*E*L6^. Smaller panels depict *f_E_* maps on inhibition planes and are as in Fig 2. The panel for *F*^*E*L6^/*F*^*I*L6^ = 3 and *S*^*I*lgn^/*S*^*E*lgn^ = 2 (row 2, column 2) corresponds to the *f_E_* rate map in Fig. 1. Other free parameters are *S^EE^* = 0.024, *S^II^* = 0.12, and *S*^*E*lgn^ = 2 × *S^EE^*.

#### 2.3 Scaling with *S^EE^* and *S^II^*

We have found *S^EE^* and *S^II^* to be the most “basic” of the parameters, and it has been productive indexing other parameters to them. In Fig. 4, we vary these two parameters in the matrix rows and columns, and examine changes in the inhibition planes.

**Fig. 4:**
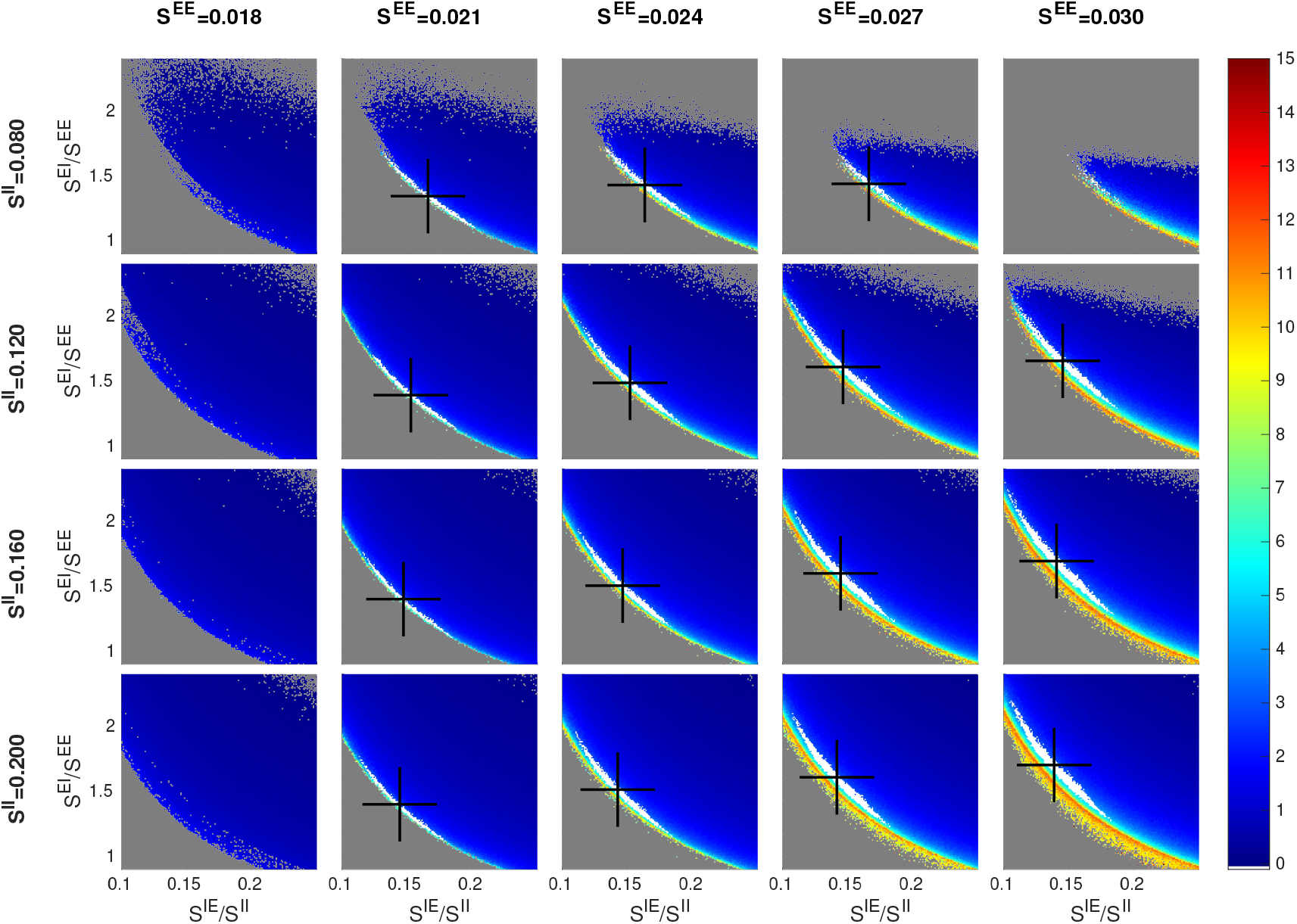
Dependence of firing rate and good area on *S^EE^* and *S^II^*. Smaller panels are as in Figures 2 and 3, with good areas (where visible) in white and black crosses denoting their centers of mass. The picture for *S^EE^* = 0.024 and *S^II^* = 0.120 (row 2, column 3) corresponds to the *f_E_* rate map in Fig. 1. Other free parameters are *S*^*E*lgn^ = 2 × *S^EE^*, *S*^*I*lgn^ = 2 × *S*^*E*lgn^, and *F*^*I*L6^ = 3 × *F*^*E*L6^.

We assume *S^EE^* ∈ (0.015,0.03). This follows from the conventional wisdom [41, 50] that when an E-cell is stimulated *in vitro*, it takes 10-50 consecutive spikes in relatively quick succession to produce a spike. Numerical simulations of a biologically realistic V1 model suggested *S^EE^* values lie well within the range above [55]. As for *S^II^*, there is virtually no direct information other than some experimental evidence to the effect that EPSPs for I-cells are roughly comparable in size to those for E-cells; see [56] and also **SI**. We arrived at the range we use as follows: With *S^II^* ∈ (0.08, 0.2) and *S^IE^*/*S^II^* ∈ (0.1,0.25), we are effectively searching through a range of *S^IE^* ∈ (0.008, 0.05). As this interval extends quite a bit beyond the biological range for *S^EE^*, we hope to have cast a wide enough net given the roughly comparable EPSPs for E and I-cells.

Fig. 4 shows a matrix of panels with *S^EE^* and *S^II^* in these ranges and the three ratios *S*^*E*lgn^/*S^EE^* = 2.5, *S*^*I*lgn^/*S*^*E*lgn^ = 2 and *F*^*I*L6^/*F*^*E*L6^ = 3. As before, each of the smaller panels shows an inhibition plane. Good areas with characteristics similar to those seen in earlier figures varying from panel to panel are clearly visible.

A closer examination reveals that (i) going along each row of the matrix (from left to right), the center of mass of the good area (black cross) shifts upward as *S^EE^* increases, and (ii) going down each column, the black cross shifts slightly to the left as *S^II^* increases, two phenomena we now explain. Again, it is important to remember that firing rates are roughly constant on the good areas.

To understand (i), consider the currents that flow into an E-cell, decomposing according to source as follows: Let [4E], [6E], [LGN] and [amb] denote the total current into an E-cell from E-cells in L4 and L6, from LGN and ambient, and let [I] denote the magnitude of the I-current. As *f_E_* is determined by the difference (or gap) between the Excitatory and Inhibitory currents, we have

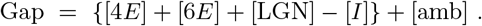

It is an empirical fact that the quantity in curly brackets is strictly positive (recall that ambient alone does not produce spikes). An increase of *x*% in *S^EE^* will cause not only [4E] to increase but also [6E] and [LGN], both of which are indexed to *S^EE^*, to increase by the same percentage. If [*I*] also increases by *x*%, then the quantity inside curly brackets will increase by *x*% resulting in a larger current gap. To maintain E-firing rate hence current gap size, [I] must increase by more than *x*%. Since I-firing rate is unchanged, this can only come about through an increase in the ratio *S^EI^*/*S^EE^*, hence the upward movement of the black crosses.

To understand (ii), we consider currents into an I-cell, and let [⋯] have the same meanings as before (except that they flow into an I-cell). Writing

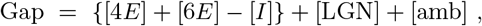

we observe empirically that the quantity inside curly brackets is slightly positive. Now we increase *S^II^* by *x*% and ask how *S^IE^* should vary to maintain the current gap. Since [LGN] and [amb] are unchanged, we argue as above that *S^IE^* must increase by < *x*% (note that 6E is also indexed to *S^IE^*). This means *S^IE^/S^II^* has to decrease, proving (ii).

We have suggested that the inhibition plane picture we have seen many times is *canonical*, or *universal*, in the sense that through any point in the designated 7D parameter cube, if one takes a 2D slice as proposed, pictures qualitatively similar to those in Figures 2–4 will appear. To confirm this hypothesis, we have computed a number of slices taken at different values of the free parameters (see **SI**), at

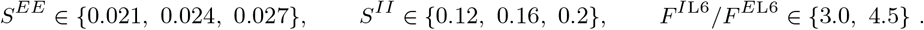

For all 18 = 3 × 3 × 2 combinations of these parameters, we reproduce the 3 × 4 panel matrix in Fig. 2, i.e., the 4D slices of (*S*^*E*lgn^/*S^EE^*,*S*^*I*lgn^/*S^II^*) × (*S^EI^*/*S^EE^*,*S^IE^*/*S*^*II*^), the first pair corresponding to rows and columns of the matrix, and the second to the xy-axes of each inhibition plane plot. All 18 plots confirm the trends observed above. Interpolating between them, we see that the contours and geometric shapes on inhibition planes are indeed universal, and taken together they offer a systematic, interpretable view of the 7D parameter space.

#### 2.4 Comparing network simulations and the MF+v algorithm

Figs. 1-4 were generated using the MF+v algorithm introduced in Sect. 1.3 and discussed in detail in **Methods**. Indeed, the same analysis would not be feasible using direct network simulations. But how accurately does the MF+v algorithm reproduce network firing rates? To answer that question, we randomly selected 128 sets of parameters in or near the good areas in Figs. 1-4, and compared the values of *f_E_* and *f_I_* computed from MF+v to results of direct network simulations. The results are presented in Fig. 5. They show that in almost all cases, MF+v overestimated network firing rates by a little, with < 20% error for ~ 80% of the parameters tested. In view of the natural variability of neuronal parameters, both within a single individual under different conditions and across a population, we view of this level of accuracy as sufficient for all practical purposes. Most of the larger errors are associated with network E-firing rates that are lower than empirically observed (at about 2 spikes/sec).

**Fig. 5:**
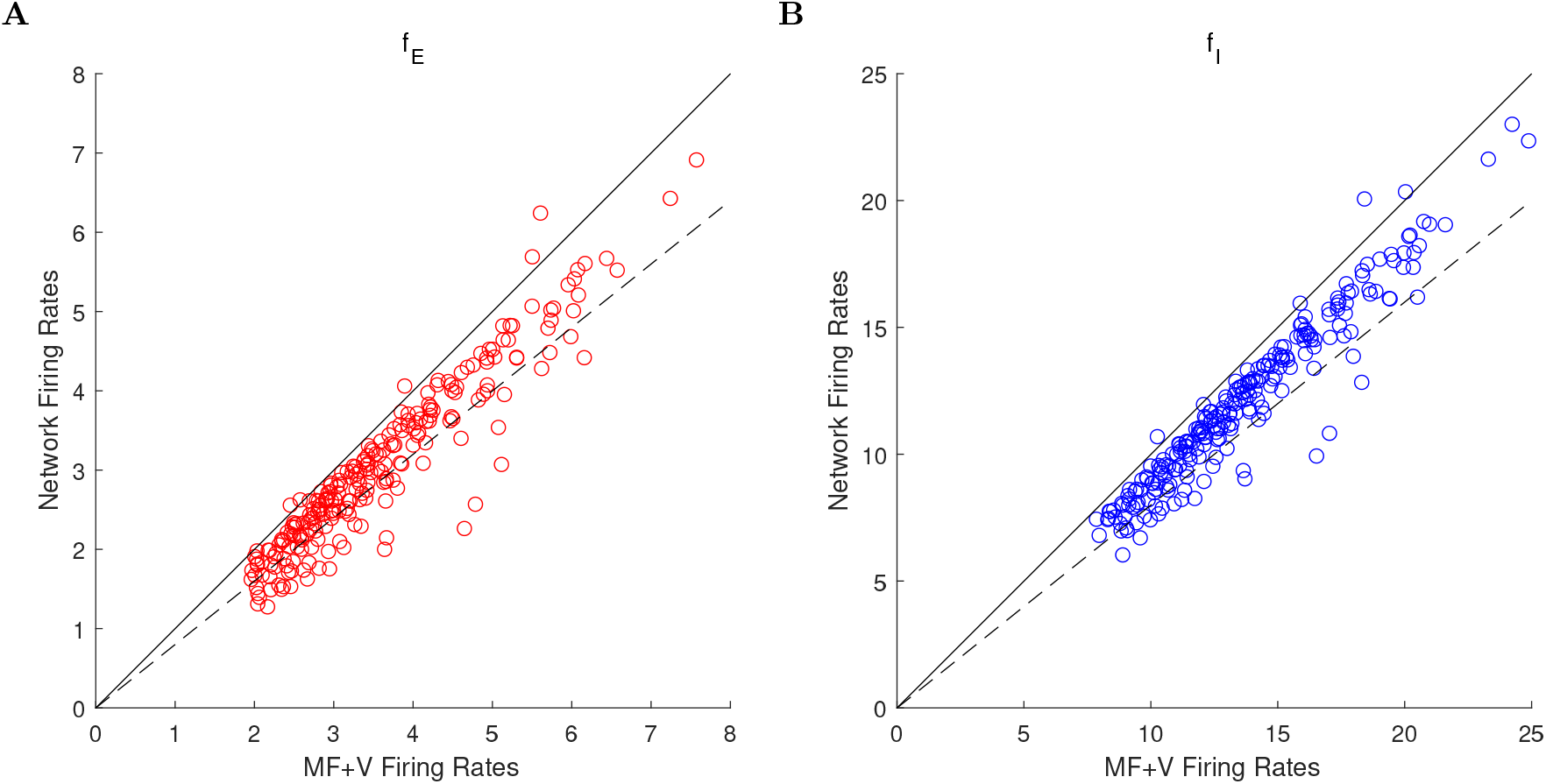
Comparison of firing rates computed using the MF+v algorithm to those from direct network simulations. The scatter plots show results for 128 sets of parameters randomly chosen in or near the good areas in Figs. 1–4. **A.** Comparison of *f_E_*. **B.** Comparison of *f_I_*. Solid lines: *y* = *x*. Dashed lines: *y* = 0.8*x*. A majority of data points fall in the range of 20% accuracy.

### 3 Other views of the viable region

We have shown in Sect. 2 that a systematic and efficient way to explore parameter dependence is to slice the viable region using inhibition planes with rescaled coordinate axes, but there are many other ways to view the 6D manifold that approximates the viable region. Here are some examples.

Fig. 6 shows two views of the viable region projected to two different low dimension subspaces. The left panel shows the viable region as a surface parametrized by hyperbolas with varying aspect ratios. This is how it looks in unscaled coordinates, compared to the panels in, e.g., Fig. 4, where we have uniformized the aspect ratios of the hyperbolas by plotting against *S^IE^/S^II^* instead of *S^IE^*. The right panel of Fig. 6 shows a bird’s-eye view of the same plot, with the {*f_E_* = 4}-contours in the (unscaled) inhibition plane, giving another view of the *S^II^* dependence.

**Fig. 6:**
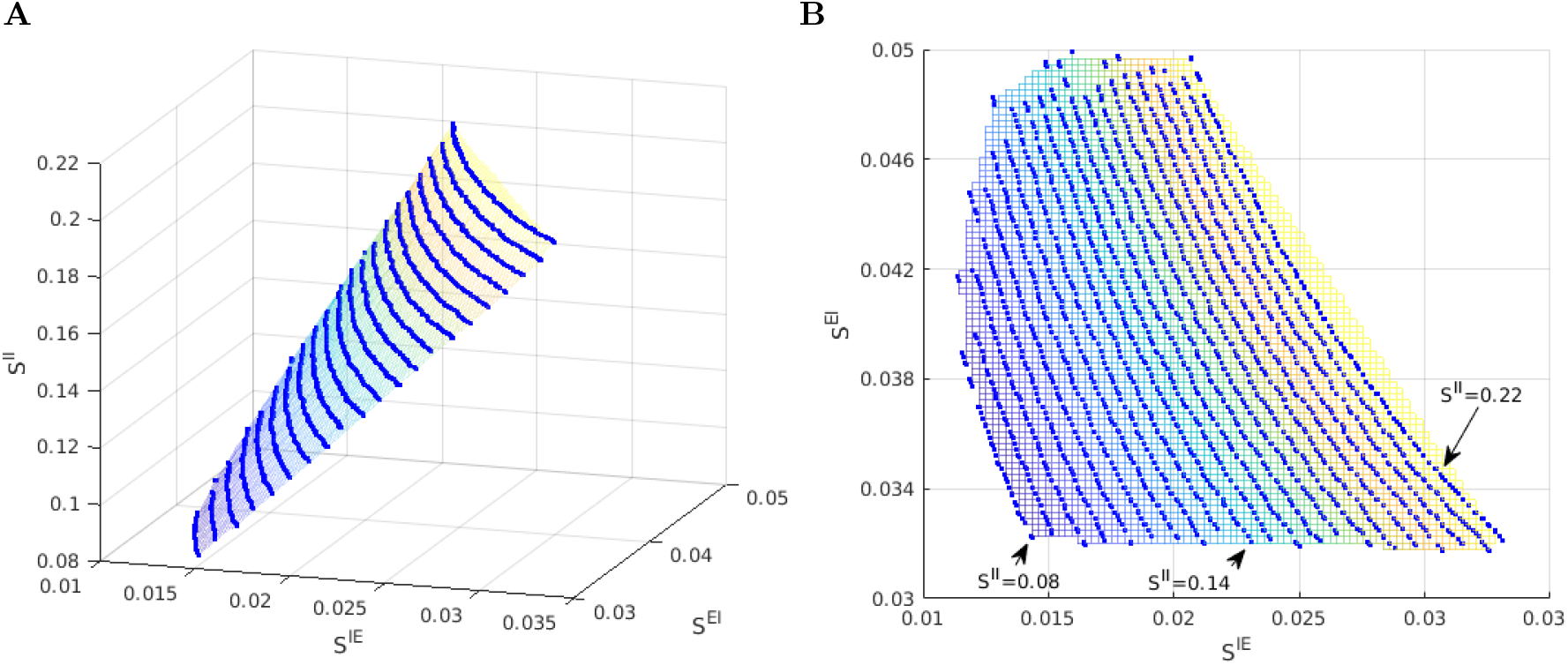
Other views of the viable regions. **A.** A 2D surface approximating the viable region projected to the 3D-space defined by *S^IE^* × *S^EI^* × *S^II^*. Blue lines are projections of contours for *f_E_* = 4 Hz intersected with the good areas, computed on 21 different inhibition planes. Parameters are fixed at *S^EE^* =0.024, *S*^*E*lgn^/*S^EE^* = 2, *S*^*I*lgn^/*S*^*E*lgn^= 2, and *F*^*I*L6^/*F*^*E*L6^ = 2; *S^II^* varies from 0.22 for the top contour to 0.08 for the bottom contour. **B.** A bird’s eye view of the left panel, with colors indicating corresponding locations of the projected E-contours.

Table 1 gives a sense of how the viable region near the reference parameter point looks when we cut through the 7D parameter cube with 1D lines. In Fig. 7, we show several heat maps for firing rates obtained by slicing the viable region with various 2D planes though the same reference point. In the top row, we have chosen pairs of parameters that covary (positively), meaning to stay in the viable region, these pairs of parameters need to be increased or decreased simultaneously by roughly constant proportions. The idea behind these plots is that to maintain constant firing rates, increased coupling strength from the E-population must be compensated by a commensurate increase in coupling strength from the I-population (left and middle panels), and increased drive to E must be compensated by a commensurate increase in drive to I (right panel). In the second row, we have selected pairs of parameters that *covary negatively*, i.e., their *sums* need to be conserved to stay in the viable region. The rationale here is that to maintain constant firing rates, total excitation from cortex and from LGN should be conserved (left and middle panels), as should total drive from L6 and from LGN (right panel).

**Fig. 7:**
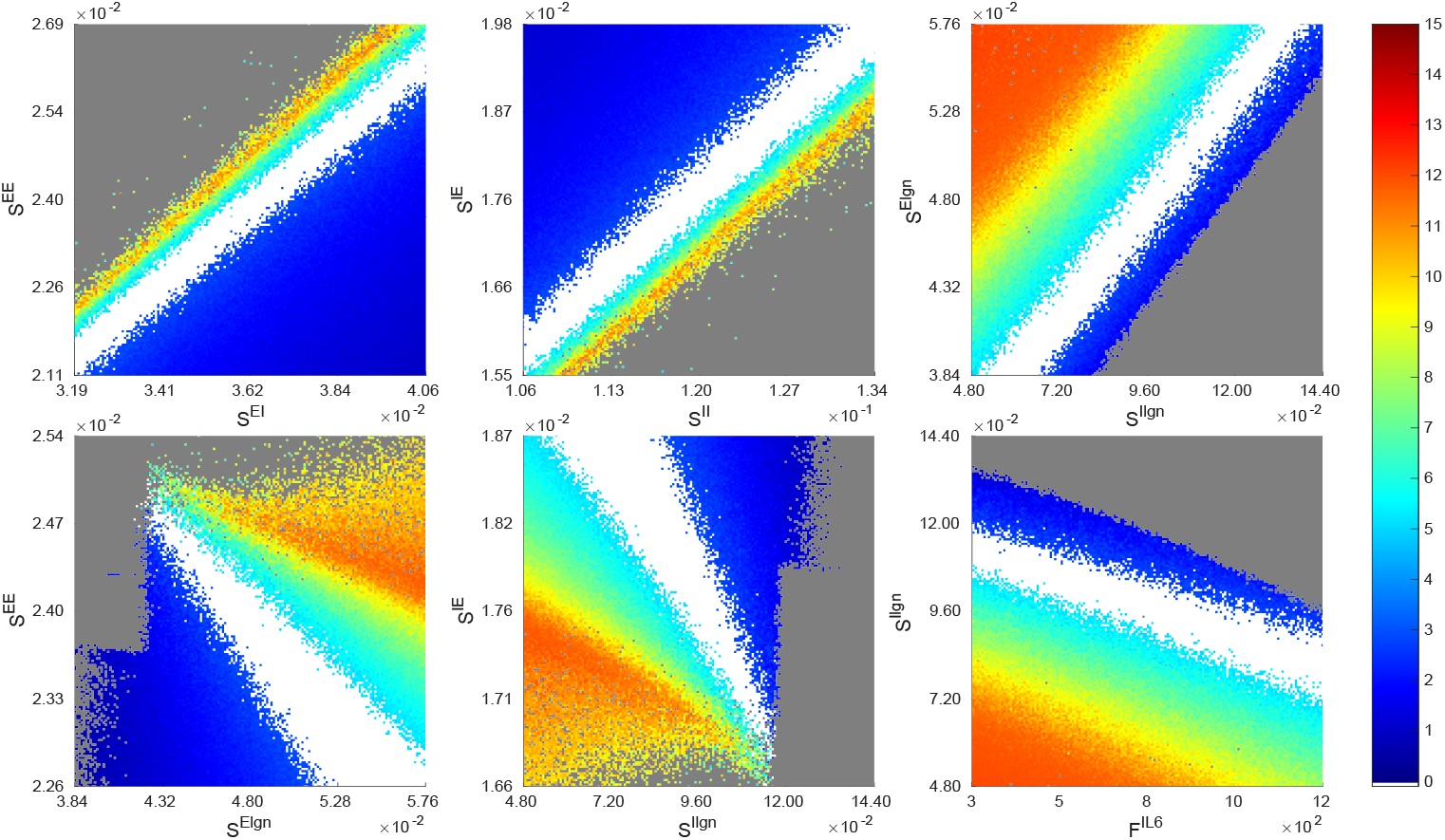
Slicing the 7D parameter space from other directions through the reference point. We compute the rate maps of *f_E_* and indicate viable regions on six other 2D slices, namely, the planes of *S^EE^* × *S^EI^*, *S^IE^* × *S^II^*, *S*^*E*lgn^ × *S*^*I*lgn^, *S^EE^* × *S*^*E*lgn^, *S^IE^* × *S*^*I*lgn^, and *S*^*I*lgn^ × *F*^*I*L6^. Firing rates are as in the color bar, and as usual the good area is in white.

Thus, together with the results in Sect. 2, we have seen three different ways in which pairs of parameters can relate: (i) they can covary, or (ii) their sums can be conserved, or, (iii) as in the case of inhibition planes, it is the *product* of the two parameters that needs to be conserved. Like (iii), which we have shown to hold ubiquitously and not just through this one parameter point, the relations in (i) and (ii) are also valid quite generally.

## Discussion

We began with some very rough *a priori* bounds for the 7 free parameters identified in Sect. 1.2, basing our choices on physiological data when available and casting a wide net when they are not. We also identified a biologically plausible region (referred to as “the viable region”) defined to be the set of parameters that lead to spontaneous E- and I-firing rates compatible with experimental values, and sought to understand the geometry of this region of parameter space. Our most basic finding is that the viable region as defined is a slightly thickened 6D submanifold – the amount of thickening varies from location to location, and is so thin in places that for all practical purposes the submanifold vanishes. This is consistent with Table 1, which shows that varying certain parameters by as little as 1% can take us out of the viable region. One can think of directions that show greater sensitivity in Table 1 as more “perpendicular” to the slightly thickened 6D surface, while those that are more robust make a smaller angle with its tangent plane. The codimension-1 property is largely a consequence of the E-I balance and has a number of biological implications, the most important of which is that the parameters giving rise to biologically plausible regimes are robust — provided one *compensates* appropriately when varying parameters. Such compensation can come about from a variety of sources *in vivo*, e.g., synaptic depression of I-neurons [57, 58]; increased thresholds for potential generation of E-spikes due to *K_v_* currents [59]; and a host of other homeostatic mechanisms [60]. To a first approximation, one can view these mechanisms as regulating synaptic weights, and our findings may be pertinent to anyone wishing to study homeostatic mechanisms governing neuronal activity.

Our analysis offers a great deal more information on the structure of the viable region beyond its being a thickened 6D manifold. We have found it profitable to slice the 7D parameter cube with *inhibition planes*, 2D planes containing the parameter axes *S^EI^* and *S^IE^*. Each inhibition plane intersects the viable region in a narrow strip surrounding a segment of a hyperbola (noted as “good area” above). Moreover, in rescaled variables *S^EI^*/*S^EE^* and *S^IE^*/*S^II^*, these hyperbolas are not only remarkably alike in appearance but their exact coordinate locations and aspect ratios vary little as we move from inhibition plane to inhibition plane, suggesting *approximate scaling relations* for firing rates as functions of parameters.

Summarizing, we found that most points in the viable region have *S^EE^* ∈ [0.02, 0.03] and *S^II^* ∈ [0.1, 0.2]; the lower limits of these ranges were found in our parameter analysis and the upper limits were *a priori* bounds. Our parameter exploration also shows that *S^EI^*/*S^EE^* ∈ (1, 2), and *S^IE^*/*S^II^* ∈ (0.1, 0.15), *S*^*E*lgn^/*S^EE^* ∈ (1.5, 3), *S*^*I*lgn^/*S*^*E*lgn^ ∈ (1.5, 2), and *F*^*I*L6^/*F*^*I*L6^ ∈ (3,4.5). We have further observed a strong correlation between degeneration of good areas and external inputs to I being too large in relation to that of E. For example, when *S*^*E*lgn^/*S^EE^* is too low, or when *S*^*I*lgn^/*S*^*E*lgn^ or *F*^*I*L6^/*F*^*E*L6^ is too high, the hyperbolic strips in inhibition planes narrow, possibly vanishing altogether. (See also Sect. S3.) We have offered explanations in terms of a *suppression index* for E-cells.

### Relation to previous work on MF

Since the pioneering work of Wilson, Cowan, Amari, and others [22–37, 61], MF ideas have been used to justify the use of firing rate-based models to model networks of spiking neurons. The basic idea underlying MF is to start with a relation between averaged quantities, e.g., an equation similar or analogous to Eq. (4), and supplement it with an “activation” or “gain” function relating incoming spike rates and the firing rate of the postsynaptic neuron, thus arriving at a closed governing equation for firing rates. MF and related ideas have yielded valuable mathematical insights into a wide range of phenomena and mechanisms, including pattern formation in slices [25, 30], synaptic plasticity and memory formation [62–66], stability of attractor networks [67–73], and many other features of network dynamics involved in neuronal computation [31, 32, 34, 36, 61, 74–100]. However, as far as we are aware, MF has not been used to systematically map out cortical parameter landscape.

Another distinction between our approach and most previous MF models has to do with intended use. In most MF models, the form of the gain function is *assumed*, usually given by a simple analytical expression; see, e.g., [39]. In settings where the goal is a general theoretical understanding and the relevant dynamical features are insensitive to the details of the gain function, MF theory enables mathematical analysis and can be quite informative. Our goals are different: our MF models are computationally efficient surrogates for realistic biological network models, models that are typically highly complex, incorporating the anatomy and physiology of the biological system in question. For such purposes, it is essential that our MF equations capture quantitative details of the corresponding network model with sufficient accuracy. In particular, we are not free to design gain functions; they are dictated by the connectivity statistics, types of afferents and overall structure of the network model. We have termed our approach “data-informed MF” to stress these differences with the usual MF theories.

We have tried to minimize the imposition of additional hypotheses beyond the basic MF assumption of a closed model in terms of rates. As summarized in Results and discussed in depth in Methods and SI, we sought to build an MF equation assuming only that the dynamics of individual neurons are governed by leaky integrate-and-fire (LIF) equations with inputs from lateral and external sources, and when information on mean voltage was needed to close the MF equation, we secured that from synthetic data using a pair of LIF neurons driven by the same inputs as network neurons. The resulting algorithm, which we have called the “MF+v” algorithm, is to our knowledge novel and is faithful to the idea of data-informed MF modeling.

As we have shown, our simple and flexible approach produces accurate firing rate estimates, capturing cortical parameter landscape at a fraction of the cost of realistic network simulations. It is also apparent that its scope goes beyond background activity, and can be readily generalized to other settings, e.g., to study evoked responses.

### Contribution to a model of primate visual cortex

Our starting point was [41], which contains a mechanistic model of the input layer to the monkey V1 cortex. This model was an ideal proving ground for our data-informed MF ideas: it is a large-scale network model of interacting neurons built to conform to anatomy. For this network, the authors of [41] located a small patch of parameters with which they reproduced many visual responses both spontaneous and evoked. Their aim was to show that such parameters existed; parameters away from this patch were not considered — and this is where they left off and where we began: Our MF+v algorithm, coupled with techniques for conceptualizing parameter space, made it possible to fully examine a large 7D parameter cube. In this paper, we identified *all* the parameters in this cube for which spontaneous firing rates lie within certain acceptable ranges. The region we found includes the parameters in [41] and is many times larger; it is a slightly thickened 6D manifold that is nontrivial in size. Which subset of this 6D manifold will produce acceptable behavior when the model is stimulated remains to be determined, but since all viable parameters – viable in the sense of both background and evoked responses – must lie in this set, knowledge of its coordinates should provide a head-start in future modeling work.

### Taking stock and moving forward

In a study such as the one conducted here, had we not used basic biological insight and other simplifications (such as inhibition planes, rescaled parameters, viable regions, and good areas) to focus the exploration of the 7D parameter space, the number of parameter points to be explored would have been *N*^7^, where *N* is the number of grid points per parameter, and the observations in Table 1 suggest that 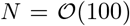 may be the order of magnitude needed. Obtaining this many data points from numerical simulation of the entire network would have been out of the question. Even after pruning out large subsets of the 7D parameter cube and leveraging the insights and scaling relations as we have done, producing the figures in this paper involved computing firing rates for ~ 10^7^ distinct parameters. That would still have required significant effort and resources to implement and execute using direct network simulations. In contrast, using the proposed MF+v algorithm, each example shown in this paper can be implemented with moderate programming effort and computed in a matter of hours on a modern computing cluster.

We have focused on background or spontaneous activity because its spatially and temporally homogeneous dynamics provide a natural testing ground for the MF+v algorithm. Having tested the capabilities of MF+v, our next challenge is to proceed to evoked activity, where visual stimulation typically produces firing rates with inhomogeneous spatial patterns across the cortical sheet. The methods developed in this paper continue to be relevant in such studies: evoked activity is often locally constant in space (as well as in time), so our methods apply to local populations, the dynamics of which form building blocks of cortical responses to stimuli with different spatiotemporal structure.

Finally, we emphasize that while MF+v provides a tremendous reduction to the cost of estimating firing rates given biological parameters, the computational cost of a parameter grid search remains exponential in the number of parameters (“curse of dimensionality”). Nevertheless, we expect our MF+v-based strategy, in combination with more efficient representations of data in high dimensional spaces (e.g., sparse grids [101]) and the leveraging of biological insight, can scale to systems with many more degrees of complexity.

## Methods

As explained in **Introduction** and **Results**, we seek parsimonious phenomenological models that are (i) simple and efficient; (ii) flexible enough to accommodate key biological features and constraints; and (iii) able to faithfully capture mean firing rates and voltages of network models across a wide range of parameters. We use for illustration a model of the monkey primary visual cortex, treating as “ground truth” the network model described in Sect. 1.1 of **Results**. Here we elaborate on the MF+v scheme outlined in Sect. 1.3, applying it to study firing rates in Layer 4C*α* (L4), an input layer to V1 in the network model.

### M1 Mean-field rate-voltage relation

We begin by stating precisely and deriving the MF equations (4) alluded to in Sect. 1.3 of Results. Consider an LIF model (Eq. (1)) for neuron *i* in L4 of the network model. We set *V*_est_ = 0, *V*_th_ = 1, and let *t*_1_, *t*_2_,…*t_n_* be the spiking times of neuron i on the time interval [0,*T*] for some large *T*. Integrating Eq. (1) in time, we obtain

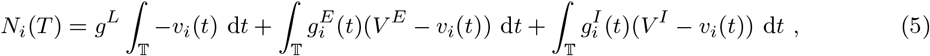

where 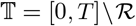, i.e., the time interval [0,*T*] minus the union 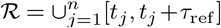 of all refractory periods. Let *f_i_* = line*T*_→∞_ *N_i_*(*T*)/*T* denote the mean firing rate of the *i*th neuron. We then have

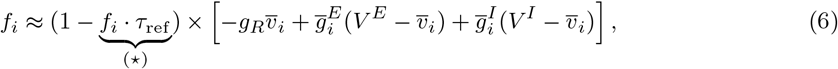

where 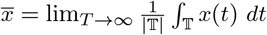 and 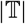 is the total length of 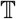. The term (⋆) is the fraction of time the *i*th neuron is in refractory, and 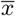 is the conditional expectation of the quantity *x*(*t*) given the cell is *not* refractory at time *t*. We have neglected correlations between conductances and voltages, as is typically done in mean-field (MF) theories. See, e.g., [29].

The long-time averages 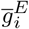 and 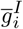 reflect the numbers and sizes of E/I-kicks received by neuron *i*. In our network model (see **SI** for details), the only source of inhibition comes from I-cells in L4, while excitatory inputs come from LGN, layer 6 (L6), ambient inputs (amb), and recurrent excitation from E-cells in L4. Mean conductances can thus be decomposed into:

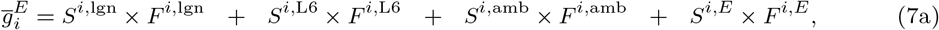

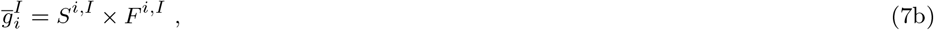

where for *P* ∈ {lgn, L6, amb, E, I}, *S^i,P^* is the synaptic coupling weight from cells in *P* to neuron *i*, and *F^i,P^* is the total number of spikes per second neuron *i* receives from source *P*, i.e., from all of its presynaptic cells in *P* combined. Here and in the rest of **Methods**, “E” and “I” refer to L4, the primary focus of the present study, so that *F^i,E^*, for example, is the total number of spikes neuron i receives from E-cells from within L4.

As discussed in the main text, we are interested in background or spontaneous activity. During spontaneous activity, we may assume under the MF limit that all E-cells in L4 receive statistically identical inputs, i.e., for each *P*, (*S^i,P^, F^i,P^*) is identical for all E-cells *i* in L4. We denote their common values by (*S^EP^, F^EP^*), and call the common firing rate of all E-cells *f_E_*. Corresponding quantities for I-cells are denoted (*S^IP^, F^IP^*) and *f_I_*. Combining Eqs. 6 and 7, we obtain the MF equations for E/I-cells:

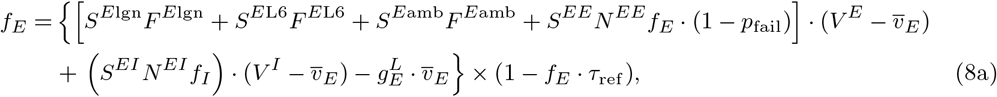

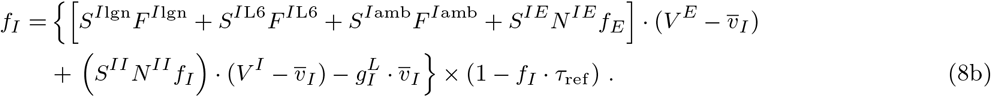

In Eq. (8), 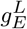 and 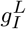xsxs are leakage conductances; *N^Q′ Q^* represents the average number of type-*Q* neurons in L4 presynaptic to a type-*Q*′ neuron in L4. These four numbers follow estimations of neuron density and connection probability of Layer 4C*α* of the monkey primary visual cortex. Refractory periods are *τ*_ref_, and E-to-E synapses are assumed to have a synaptic failure rate *p*_fail_, also fixed. Details are discussed in **SI**.

We seek to solve Eq. (8) for (*f_E_*, *f_I_*) given network connections, synaptic coupling weights and external inputs. That is, we assume all the quantities that appear in Eq. (8) are fixed, except for *f_E_*, *f_I_*, 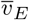 and 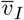. The latter two, the mean voltages 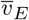 and 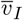, cannot be prescribed freely as they describe the sub-threshold activity of L4 neurons once the other parameters are specified. Thus what we have is a system that is not closed: there are four unknowns, and only two equations.

A second observation is that once 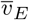 and 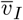 are determined, Eq. (8) has a very simple structure. To highlight this near-linear structure, we rewrite Eq. (8) in matrix form as

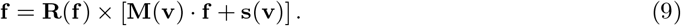

Here **f** = (*f_E_*, *f_I_*), 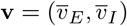, **M**(**v**) is a (voltage-dependent) linear operator acting on L4 E/I firing rates, **s** includes inputs from external sources and leakage currents, and **R** accounts for refractory periods. (See Sect. S3.)) Neglecting refractory periods, Eq. (9) is linear in **f** assuming **M**(**v**) and **s**(**v**) are known. At typical cortical firing rates in background, the refractory factor **R** contributes a small nonlinear correction.

To understand the dependence on **v**, we show in Fig. 8A the level curves of *f_E_* and *f_I_* as functions of 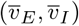 from the mapping defined by Eq. (8). As expected, (*f_E_*, *f_I_*) vary with 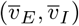, the nearly straight contours reflecting the near-linear structure of Eq. (9). One sees both *f_E_* and *f_I_* increase steadily (in a nonlinear fashion) as we decrease 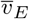 and increase 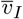, the dependence on 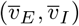 being quite sensitive in the lower right part of the panels. The sensitive dependence of *f_E_* and *f_I_* on 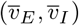 rules out arbitrary choices on the latter in a MF theory that aims to reproduce network dynamics. How to obtain reasonable information on 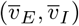 is the main issue we need to overcome.

**Fig. 8:**
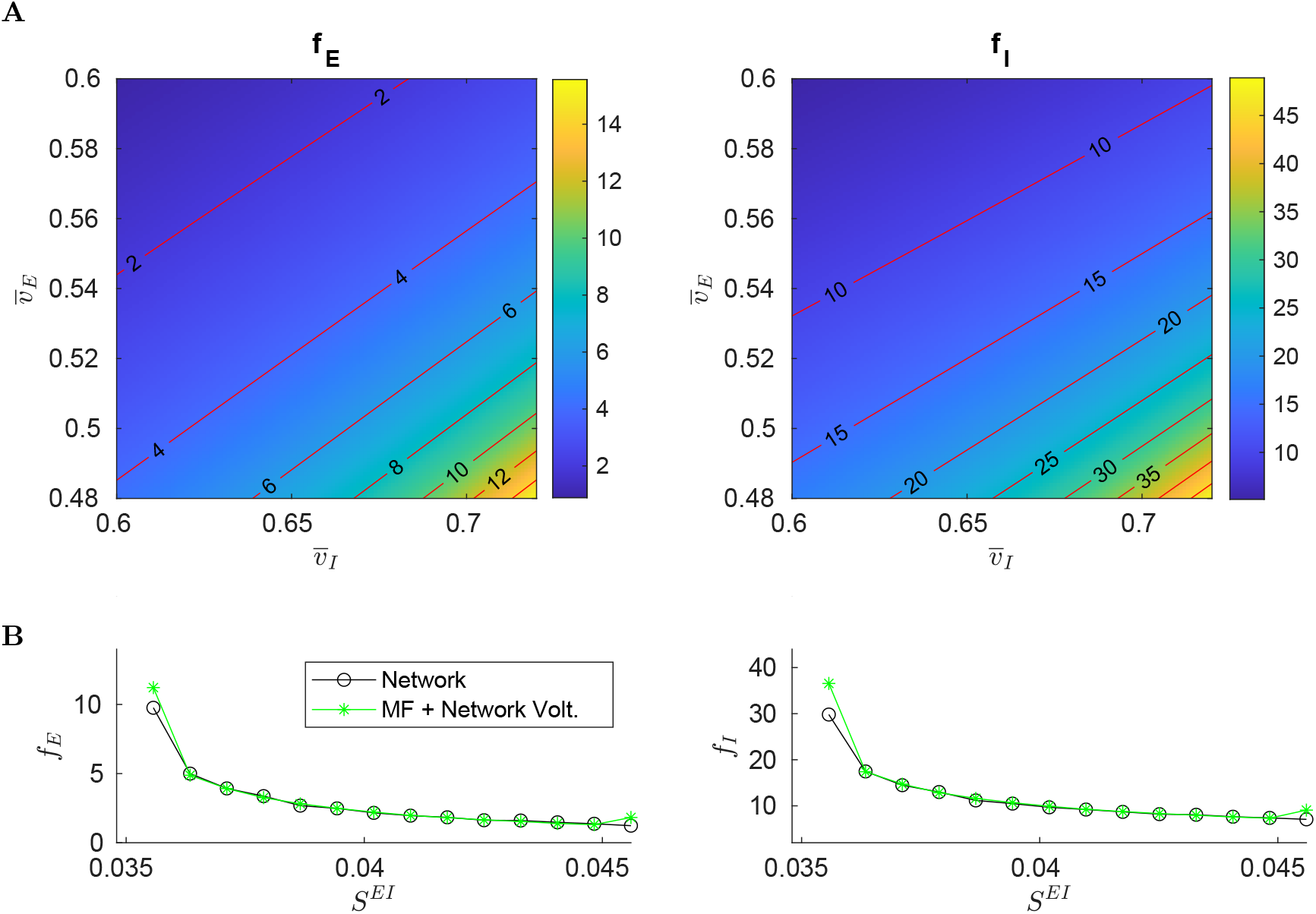
MF approximations. **A.** Contours of E/I firing rates as functions of 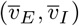. **B.** Comparison of network firing rates (black) and (*f_E_*, *f_I_*) computed from Eq. (8) using network-computed mean voltages (green).

We propose in this paper to augment Eq. (8) with values of 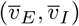 informed by (synthetic) data. To gauge the viability of this idea, we first perform the most basic of all tests: We collect firing rates and mean voltages (averaged over time and over neurons) computed directly from network simulations, and compare the firing rates to *f_E_* and *f_I_* computed from Eq. (8) with 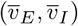 set to network-computed mean voltages. The results for a range of synaptic coupling constants are shown in Fig. 8B, and the agreement is excellent except when firing rates are very low or very high.

### M2 The MF+v algorithm

Simulating the entire network to obtain 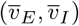 defeats the purpose of MF approaches, but the results in Fig. 8B suggest that we might try using single LIF neurons to represent typical network neurons, and use them to estimate mean voltages.

The idea is as follows: Consider a pair of LIF neurons, one E and one I, and fix a set of parameters and external drives. In order for this pair to produce 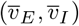) similar to the mean voltages in network simulations, we must provide these cells with surrogate inputs that mimic what they would receive if they were operating within a network. However, the bulk of the input into L4 cells are from other L4 cells. This means that in addition to surrogate LGN, L6, and ambient inputs, we need to provide our LIF neurons *surrogate L4 inputs* (both E and I) commensurate with those received by network cells. Arrival time statistics will have to be presumed (here we use Poisson), but firing rates should be those of L4 cells – the very quantities we are seeking from our MF model. Thus there is the following consistency condition that must be fulfilled: For suitable parameters and external inputs, we look for values 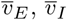, *f_E_*, and *f_I_* such that

- given 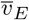 and 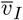, Eq. (8) returns *f_E_* and *f_I_* as firing rates; and
- when L4 firing rates of *f_E_* and *f_I_* are presented to the LIF pair along with the stipulated parameters and external inputs, direct simulations of this LIF pair return values of 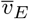 and 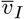.

If we are able to locate values of 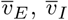, *f_E_*, and *f_I_* that satisfy the consistency relations above, it will follow that the LIF pair, acting as surrogate for the E and I-populations in the network, provide mean voltage data that enable us to determine mean network firing rates in a self-consistent fashion.

A natural approach to finding self-consistent firing rates is to alternate between estimating mean voltages using LIF neurons receiving surrogate network inputs — including L4 firing rates from the previous iteration — and using the MF formula (Eq. (4)) to estimate L4 firing rates using voltage values from the previous step. A schematic representation of this iterative method is shown in Fig. 9A. In more detail, let **v**_*p*_ and **f**_*p*_ be the mean voltage and firing rate estimates obtained from the pth iteration of the cycle above. In the next iteration, we first simulate the LIF cells for a prespecified duration *t^LIF^*, with L4 firing rate set to **f**_*p*_, to obtain an estimate **v**_*p*+1_ = LIF(**f**_*p*_, *t^LIF^*) of the mean voltage. We then update the rate estimate by solving

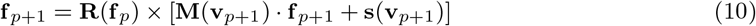

for **f**_*p*+1_, leading to

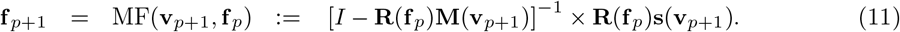

**Fig. 9:**
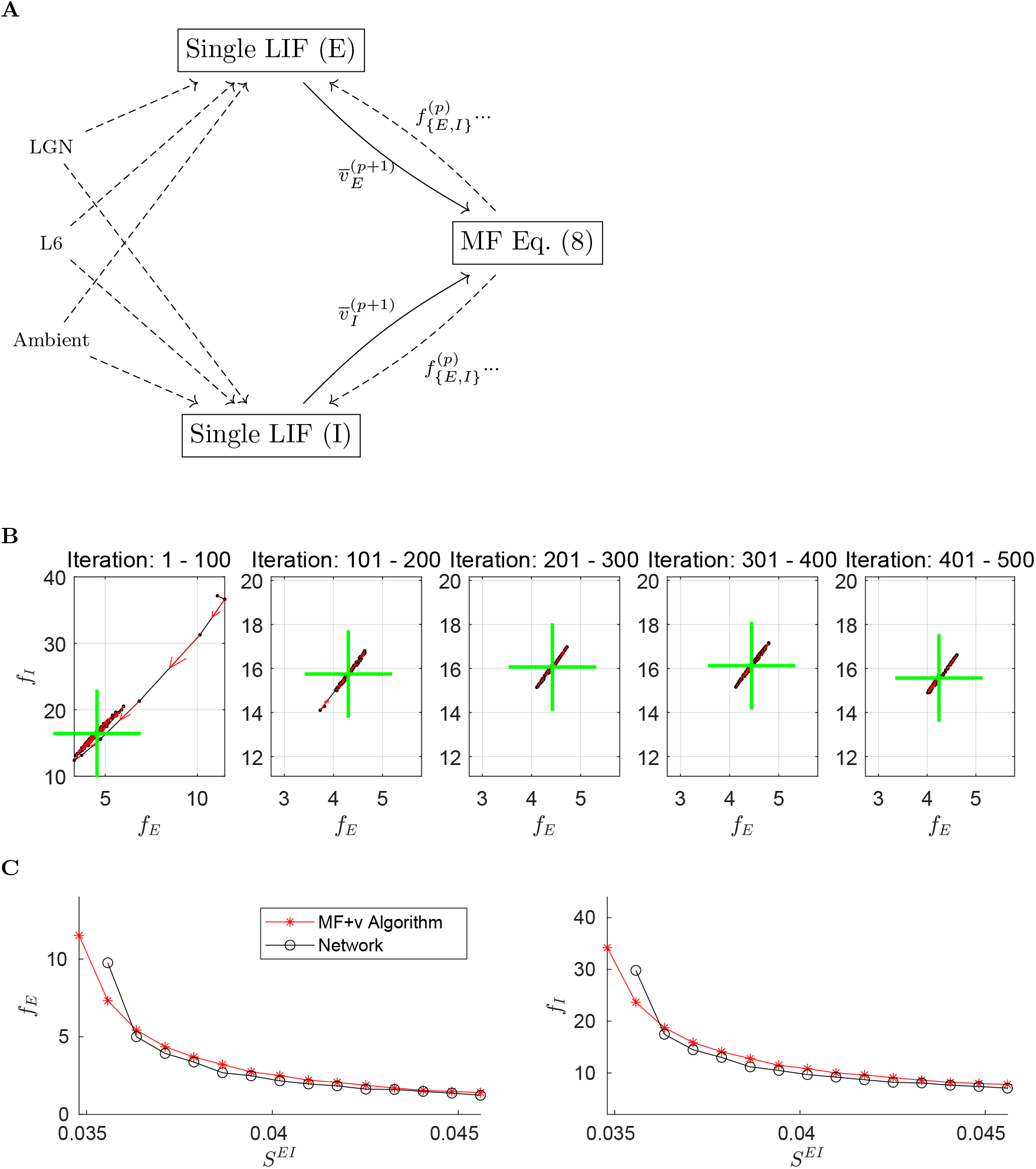
The MF+v algorithm. **A.** Schematic of the algorithm. We begin by choosing initial values 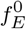 and 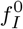. These values are used to drive a pair of LIF neurons for 20 seconds (biological time). The resulting membrane voltages are then fed into the MF equation (8), which gives us firing rates for the next iteration. This is repeated until certain convergence criteria are met (see text). In the above, all dashed lines are modeled by Poisson processes. **B.** 500 training iterations of the MF+v. The means of every 100 iterations are indicated by green crosses, stability properties of which are quite evident. **C.** Comparison of network (black) and MF+v computed (red) firing rates.

When the iteration converges, i.e., when **f**_*p*_ = **f**_*p*+1_ = **f** and **v**_*p*_ = **v**_*p*+1_ = **v**, we have a solution (**f**, **v**) of Eq. (9).

Fig. 9B shows 500 iterations of this scheme in **f**_*p*_-space. Shown are iterations and running means (green crosses). Observe that after transients, the **f**_*p*_ settle down to what appears to be a narrow band of finite length, and wanders back and forth along this band without appearing to converge. We have experimented with doubling the integration time *t^LIF^* and other measures to increase the accuracy of the voltage estimates **f**_*p*_, but they have had no appreciable impact on the amplitudes of these oscillations. A likely explanation is that the contours of the E/I firing rates (Fig. 8A) are close to but not exactly parallel: Had they been exactly parallel, a long line of **v** would produce the same **f**, implying the MF equations (8) do not have unique solutions. The fact that they appear to be nearly parallel then suggests a large number of near-solutions, explaining why our attempt at fixed point iteration cannot be expected to converge in a reasonable amount of time.

However, the oscillations shown in Fig. 9B are very stable and well defined, suggesting a pragmatic way forward: After the iterations settle to this narrow band, we can run a large number of iterations and average the **f**_*p*_ to produce a single estimate. Specifically, we first carry out a number of “training” iterations, and when the firing rate estimates settle to a steady state by a heuristic criterion, we compute a long-time average and output the result. Combining this with the MF formula (4) yields the MF+v algorithm. See **SI** for more details.

Fig. 9C compares MF+v predictions with network simulations. As one would expect, averaging significantly reduces variance. The results show strong agreement between MF+v predictions and their target values given by direct network simulations.

Finally, we remark that a natural alternative to our MF+v algorithm might be to forgo the MF equation altogether, and construct a self-consistent model using single LIF neurons. We found that such an LIF-only method is much less stable than MF+v. (See **SI**.) This is in part because firing rates require more data to estimate accurately: For each second of simulated activity, each LIF neuron fires only 3-4 spikes, whereas we can collect a great deal more data on voltages over the same time interval.

### M3 Issues related to implementation of MF+v

The hybrid approach of MF+v, which combines the use of the MF formula (4) with voltage estimates from direct simulations of single neurons, has enabled us to seamlessly incorporate the variety of kick sizes and rates from L6, LGN, and ambient inputs while benefiting from the efficiency and simplicity of Eq. (4). Nevertheless there are some technical issues one should be aware of.

#### Failure to generate a meaningful result

We have found that MF+v iterations can fail to give meaningful answers in the following two situations. First, in some parameter regimes, the linear operator **M**(**v**) can become singular, which can result in unreasonably low or even negative values of **f**_*p*_ in the MF+v algorithm (see **SI**). Second, when firing rates are too low (e.g., *f_E_* < 0.1 Hz), the low rate of L4 kicks to the LIF neurons in the MF+v model can result in large fluctuations of **v**_*p*_, which can destabilize the computed **f**_*p*_ unless the integration time *t^LIF^* is sufficiently large.

In **Results**, whenever MF+v fails, we exclude the parameter set and label it with a gray pixel in the canonical picture; see, e.g., Fig. 1.

#### Computational cost

The majority of computation in MF+v is spent on collecting the mean voltages **v**_*p*_ in each iteration, which can be time-consuming depending on the time-scale and accuracy of LIF-neuron simulation. By repeating for *M* iterations, the computational cost of MF+v is *O*(*Mt*^LIF^), where a general choice of *t^LIF^* ∈ (5, 40)s. If we simulate each neuron for *t^LIF^* = 20s per iteration for up to *M* = 100 iterations, we typically obtain a firing rate estimate within ~ 20% accuracy of the network firing rate in ~ 1.5s. In contrast, the cost of simulating a large network with *N* neurons typically grows at least as fast as *N*, and may even grow nonlinearly in *N*. With the parameters in this paper, a typical network simulation using *N* ≈ 4 × 10^4^ cells may require up to ~ 60 seconds, which is substantial when used to map a 7-dimensional parameter grid. The MF+v algorithm thus represents an ~ 40-fold speedup over the corresponding large network simulation in contemporary computing environments.^2^

It is possible to further reduce the computational cost of MF+v. First, instead of the simple iteration scheme we used in MF+v (i.e., Picard iteration), one can use a stochastic variant of a higher-order method (see, e.g., [102]). Second, one need not make an independent computation of the mean voltages **v**_*p*_ for each iteration. Instead, we can precompute the mean voltages for a coarse grid in (*f_E_*, *f_I_*) space, then, by interpolation and smoothing, construct a table of values of mean voltages as functions of L4 rates. This approximation can then be used as a surrogate for the Monte Carlo simulation presently used in MF+v. In certain regimes, it may also be possible to compute **v**_*p*_ in a more analytical manner, e.g., via the Fokker-Planck equation for LIF neurons.

## Acknowledgements

ZCX was supported by the Swartz Foundation. KL was supported in part by NSF grant DMS-182128. LSY was supported in part by NSF grants SMA-1734854 and DMS-1901009.

## Author Contributions

Conceptualization, Methodology, Analysis, Writing: ZCX, LSY, KL. Data Curation and Software: ZCX.

## Supplementary Information

### S1 Details of network model: network architecture and governing equations

In this section and the next, we describe the V1 network model we view as “ground truth”. Except for minor simplifications, we follow [41]; interested readers are directed to [41] and references therein for more details.

#### “External” input to layer 4Cα neurons

We model a small piece of layer 4C*α* of the Macaque primary visual cortex (V1), focusing on the background regime, i.e., spontaneous network activity. Layer 4C*α* neurons deliver excitatory and inhibitory signals to each other. In addition, each Layer 4C*α* neuron receives excitatory input from three categories of external (meaning external to 4C*α*) sources, which we label *LGN, Layer 6*, and *ambient.* “LGN” refers to input from the lateral geniculate nucleus, which carries visual signals from the retina; “Layer 6” represents recurrent excitation from Layer 6 of V1; “ambient” is an amalgamation of weak cortical neuromodulatory signals. Since our primary goal is to investigate background firing rates (when external input rates are low), we model signals from all three sources as statistically independent, approximately Poisson (Bernoulli) processes, i.e., to generate a point process of rate *R* with timestep Δ*t*, we place 0 or 1 spike in each time bin with probability *R*Δ*t*.

#### Architecture of layer 4Cα network in V1

Consider a part of a 2D cortical sheet representing 3 × 3 hypercolumns, each occupying 0.5 × 0.5 mm^2^ in layer 4C*α*. This region contains 26,244 E-cells and 8,649 I-cells, which we assume are uniformly distributed, resulting in 3,877 cells (2,916 E, 961 I) per hypercolumn. In our model, E-cells are assumed to be spiny stellate cells and I-cells PV basket cells. The strength of connectivity between cells depends on the cell types (E or I, to be discussed later in *Equations* and Sect. S2), while the probability of connection between any two cells in a local circuit is determined by a truncated Gaussian function of cell-to-cell distances. (In layer 4C*α* there are no long-range connections.) For E-to-Q and I-to-Q connections, the standard deviations of the Gaussian functions are 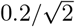 mm and 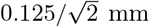, respectively, reflecting the different reach of E and I cells. The peak connection probability for E→E is 0.15, while the numbers for E→I, I→E, and I→I are set as 0.6, due to the much denser I-cell connections. We truncate all the Gaussian functions at *X^o^* = 0.36 mm. Specifically, for a pair of neurons (*i,j*) which are *x* mm away from each other, the projection probability from *j* to *i* is

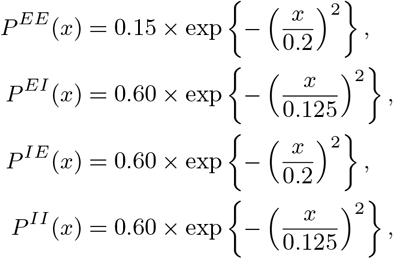

and

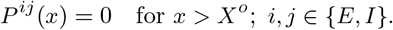

The connection probabilities and cell densities result in that, on average, each E-cell has ‘---210 presynaptic E-cells and ~110 I presynaptic cells, while each I -cell has ~840 presynaptic E-cells and ~110 presynaptic I-cells. We leave the neurons close to the boundary as is and choose not to compensate for the missed presynaptic neurons.

##### Equations

We use conductance-based leaky-integrate-and-fire (LIF) models for neuronal dynamics within the Layer 4C*α* network. For a neuron *i* with cell type *Q* ∈ {*E,I*}, its membrane potential *v_i_* advances by

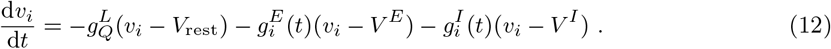

We nondimensionalize voltages, setting resting potential to *V*_rest_ = 0 and spiking threshold *V*_th_ = 1. Every time *v_i_* reaches *V*_th_, a spiking event occurs at neuron *i* and the signal is sent to all its postsynaptic neurons. Afterwards, *v_i_* enters a refractory period for *τ*_ref_ = 2 ms right away, then reset to *V*_rest_.

With the selections of the reversal potentials *V^E^* = 14/3 and *V^I^* = —2/3 [49], the membrane potential *v_i_* is driven by three current terms in Eq. (12):

i. Towards *V*_rest_ = 0 due to the leaky current 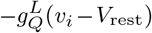, where 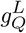 stands for membrane leakage conductance of cell type *Q*.
ii. Towards *V^I^* = −2/3 due to the inhibitory current 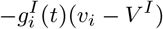, where the inhibitory conductance 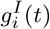 is determined by the spiking series generated by inhibitory cells presynaptic to neuron *i*, i.e.,

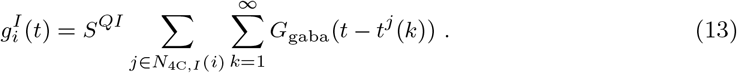 Here, *S^QI^* stands for the connectivity from an I-cell to a type-*Q* cell. Cell *j* belongs to *N*_4*C,I*_(*i*), the collection of all presynaptic I-cells to neuron *i*, generating a spiking time series {*t^j^*}. In addition, *G*_gaba_(*t*) is a Green’s function modeling the temporal increment of inhibitory conductances induced by each I-spike through GABA receptors (details provided in Sect. S2).
iii. Towards *V^E^* = 14/3 due to the excitation current 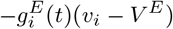. The excitatory conductance of neuron *i*, 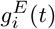, consists of four components:

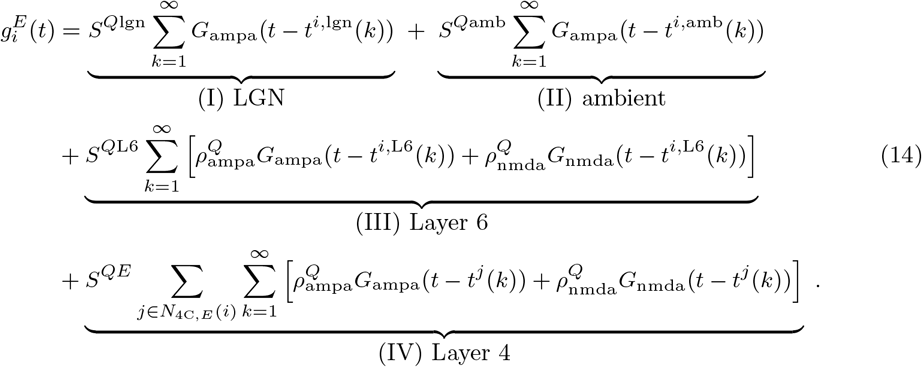 Terms I-IV represent synaptic conductances induced by LGN, ambient, Layer 6 input, and Layer 4 recurrent excitation, respectively. For each neuron *i*, the spiking series in terms I-III are modeled by Poisson processes as described above. For E-spikes from Layer 4 and 6, two different types of excitatory synapses (AMPA and NMDA) induce different temporal increment of 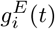 (term III and IV), while only AMPA synapse is involved for LGN and ambient input (term I and II). Here, 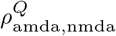 stand for the fractions of synaptic input received by AMPA and NMDA receptors in a type-*Q* neuron; (*S*^*Q*lgn^, *S*^*Q*amb^, and *S*^*Q*L6^, *S^QE^*) denote the respective synaptic coupling weights of these sources towards type-*Q* cells.

For the E-to-E input in Layer 4, two additional biological details are incorporated in the model. First, we consider a possibility for synaptic failure *p*_fail_ = 20%, i.e., whether the *k*-th spike from neuron *j* successfully induces a change in 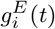 depends on an independent coin-toss with *p* = 0.8. Second, if a spike is “successful”, a random delay is added to *t^j^*(*k*) to model the fact that E-neurons project to the dendrites of other E-cells, instead of the soma. In all, when neuron i is an E-neuron, the term IV in Eq. (14) is replaced by

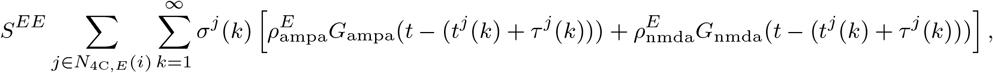

where *σ^j^*(*k*) stands for the coin-toss, and the *τ^j^*(*k*) are independent, identically distributed random delay times uniformly distributed on [0, 1] ms.

### S2 Parameters

We now list the specific parameter values used. We remark that while EPSC and IPSCs have been measured in the laboratory and so can be assumed to be known, the coupling weights — which involve how one neuron affects another — cannot be measured directly. This is the main reason we mostly regard the coupling weights as free parameters to be investigated in this paper, in spite of experimental evidence that may provide certain ranges for them.

#### Neuronal parameters

The following parameters are used for L4 neurons, and are fixed throughout the paper.

i. Reversal potentials: *V^I^* = —2/3, *V^E^* = 14/3
ii. Leakage conductances: 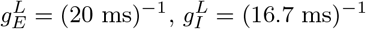
iii. Postsynaptic conductances:

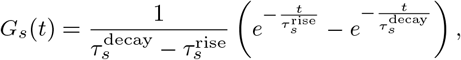

where 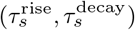 stand for the time scales of activation/inactivation of synapse type *s* = ampa, nmda, gaba; the time constants used here are

- 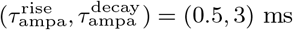
- 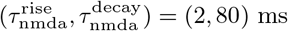
- 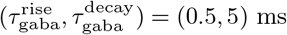
iv. Fraction of AMPA and NMDA receptors activated by E-spikes: 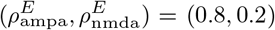, and 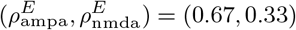
v. Synaptic failure: *σ^j^*(*k*) = 1 with *p* = 0.8, and *σ^j^*(*k*) = 0 with *p* = 0.2
vi. Synaptic delays: the *τ^j^*(*k*) are uniformly distributed between [0,1] ms
vii. Refractory period: *τ*_ref_ = 2 ms

For biophysical constants, see, e.g., [49]. We follow [41, 103] for all other parameters.

#### Network parameters

These include all synaptic coupling weights and rates of external input, making up the parameter space in which we compute the landscape of E- and I-firing rates. We specify below our choices of them. Those parameters that are specified with ranges form the 7D space of *free parameters* we investigate in this paper. These *a priori* constraints of free parameters are discussed later in this section. Generally, we first choose the values of *S^EE^* and *S^II^* (independently of other parameters), then index some of the other parameters to *S^EE^* and *S^II^*.

i. Synaptic coupling weights chosen independently: *S^EE^* ∈ (0.018, 0.030), *S^II^* ∈ (0.08, 0.20)
ii. Synaptic coupling weights depending on *S^EE^*:

- *S*^*E*lgn^ ∈ (1.5, 3) × *S^EE^*, *S*^*I*lgn^ ∈ (1.5, 3) × *S*^*E*lgn^
- 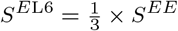
- *S^EI^* ∈ (0.9, 2.4) × *S^EE^*
iii. Synaptic coupling weights depending on *S^II^*:

- *S^IE^* ∈ (0.1, 0.25) × *S^II^*
- *S*^*I*L6^ = 3 × *S^IE^*
iv. LGN input rates: *F*^*E*lgn^ = *F*^*I*lgn^ = 80 Hz
v. Layer 6 input rates: *F*^*E*L6^ = 250 Hz, *F*^*I*L6^ ∈ (1.5, 6) × *F*^*E*L6^
vi. Ambient input:

- *F*^*E*amb^ = *F*^*I*amb^ = 500 Hz
- *S*^*E*amb^ = *S*^*I*amb^ 0.01

#### Prior biological constraints and scaling conventions

We justify our choices of the ranges of *free parameters* above. Following (often indirect) suggestions from experimental observations and heuristic reasoning, one can arrive at some bounds on them. The ranges we impose are broader than those suggested by available data; the greater the uncertainty, the wider the net we cast. Specifically:

- *S^EE^* ∈ (0.015, 0.03): This follows from the conventional wisdom that when an E-cell is stimulated *in vitro*, it takes 10-50 consecutive spikes in relatively quick succession to produce a spike. Numerical simulation suggests the assumed order of magnitude for *S^EE^* is reasonable [55].
- *S^EI^* ∈ (0.9, 2.4) × *S^EE^* and *S^IE^* ∈ (0.1, 0.25) × *S^II^*: In the absence of experimental guidance, we located these ranges numerically as follows: We examined firing rate maps for wider ranges than these, and found that, for the most part, the geometry on inhibition planes forces the good areas to lie within these ranges.
- *S^II^* ∈ (0.08, 0.20): There is no direct empirical information on this parameter; however, there is evidence that EPSPs for I-cells are similar in size to those for E-cells [56]. We choose the range for *S^II^* by following the logistic that *S^II^* ∈ (0.08, 0.20) and *S^IE^* ∈ (0.1, 0.25) × *S^II^* means *S^IE^* ∈ (0.008, 0.05), which contains and is significantly larger than the range of *S^EE^* above.
- *S*^*E*lgn^ ∈ (1.5, 3.0) × *S^EE^*: Results from [50] suggest that the sizes of EPSPs from LGN are ~ 2× those from L4. We therefore assume a range around 2.
- *S*^*I*lgn^ ∈ (1.5,3.0) × *S*^*E*lgn^: we assume *S*^*I*lgn^ > *S*^*E*lgn^ because it has been reported that LGN produces larger EPSCs in I-cells [52].
- *F*^*I*L6^ ∈ (1.5, 6) × *F*^*E*L6^: Within L4, an I-cell has 3.5-4 times as many presynaptic E-neurons as an E-cell. If we hypothesize a similar ratio between L6 and L4, it would follow that *F*^*I*L6^ ∈ (3.5,4) × *F*^*E*L6^.xsxsxs We relax the interval to (1.5, 6) because of uncertainty surrounding L6: whether the effect of L6 on L4 is net-excitatory or net-inhibitory is an issue that is currently unresolved for the real cortex. A wider range also serves to absorb potential errors in the assumption that 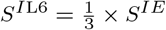.

### S3 Closer look at MF+v: solvability of MF equation and implementation details

#### Biologically meaningful solutions of MF equation

In some situations, the MF equation (4) yields negative firing rates when given valid mean voltages; we have indicated such parameters by gray in all parameter plots. Here, we discuss some of the reasons underlying these failures.

First, recall that in Methods, we had asserted that the MF equation Eq. (8) can be written in matrix form as

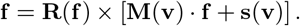

(This was Eq. (9) in Methods.) This can be seen by defining

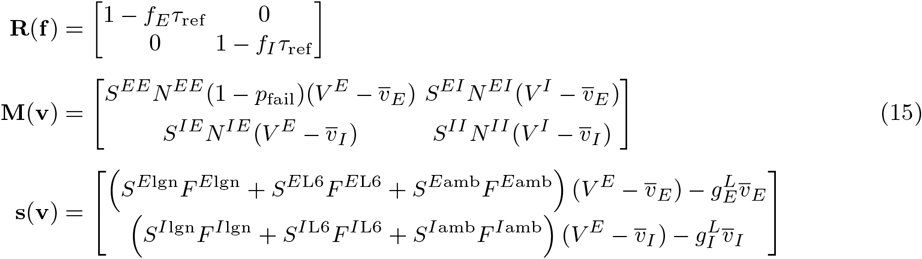

and verifying directly. Our interest is in finding nonnegative solutions **f** of this equation given mean voltages **v**. The solvability of Eq. (9) depends on the properties of the matrix **M**(**v**), which in turn depends on L4 connectivity and the mean voltages **v**. Note that the entries of **s**(**v**) represent the mean currents into E and I cells, respectively, and must be positive for cells to fire with positive rates.

Among the scenarios in which these equations fail to give meaningful firing rates, by far the simplest is when the equations are (nearly) singular. Ignoring the refractory factor **R**(**f**) (whose effect is perturbative), Eq. (9) is equivalent to

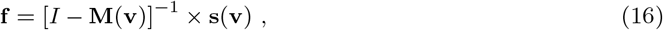

where

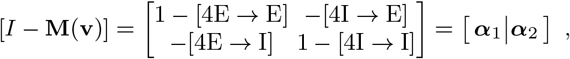

provided *I* – **M**(**v**) is nonsingular. In the above, [P → Q] indicate the corresponding entry of matrix *M*(**v**), i.e., the net contribution to an E/I-cell from one E/I-kick. When det(*I* – **M**(**v**)) = 0, the linearized equation above may not have a solution, suggesting MF+v iteration is likely to fail when I — **M**(**v**) is (nearly) singular.

One can in fact take a more geometric view of the solvability of Eq. (9), one that makes questions surrounding solvability more transparent. Let ***α***_1_ and ***α***_2_ be the columns of *I* – **M**(**v**). Eq. (9) yields nonnegative firing rates precisely when there exist *f*_1_, *f*_2_ ≥ 0 such that *f*_1_***α***_1_ + *f*_2_***α***_2_ = **s(v)**. This can be visualized by defining *S* = {*f*_1_***α***_1_ + *f*_2_***α***_2_ | *f*_1_, *f*_2_ ≥ 0} and noting that the intersection of *S* with the first quadrant {*s*_1_, *s*_2_ ≥ 0} is precisely the set of all **s** such that **f** = (*I* – **M**(**v**))^-1^**s** results in nonnegative firing rates. Fig. S1A shows an example. Here, the two (normalized) column vectors ***α***_1_ and ***α***_2_ are linearly independent, and the set S (the region bounded by span(***α***_1_) and span(***α***_2_) and contains the black line) has a large intersection with the first quadrant, so that *most* nonnegative values of **s** lead to nonnegative rates. Note that these parameters lie well above the good area; *cf.* Figs. 1 and S1.

**Fig. S1:**
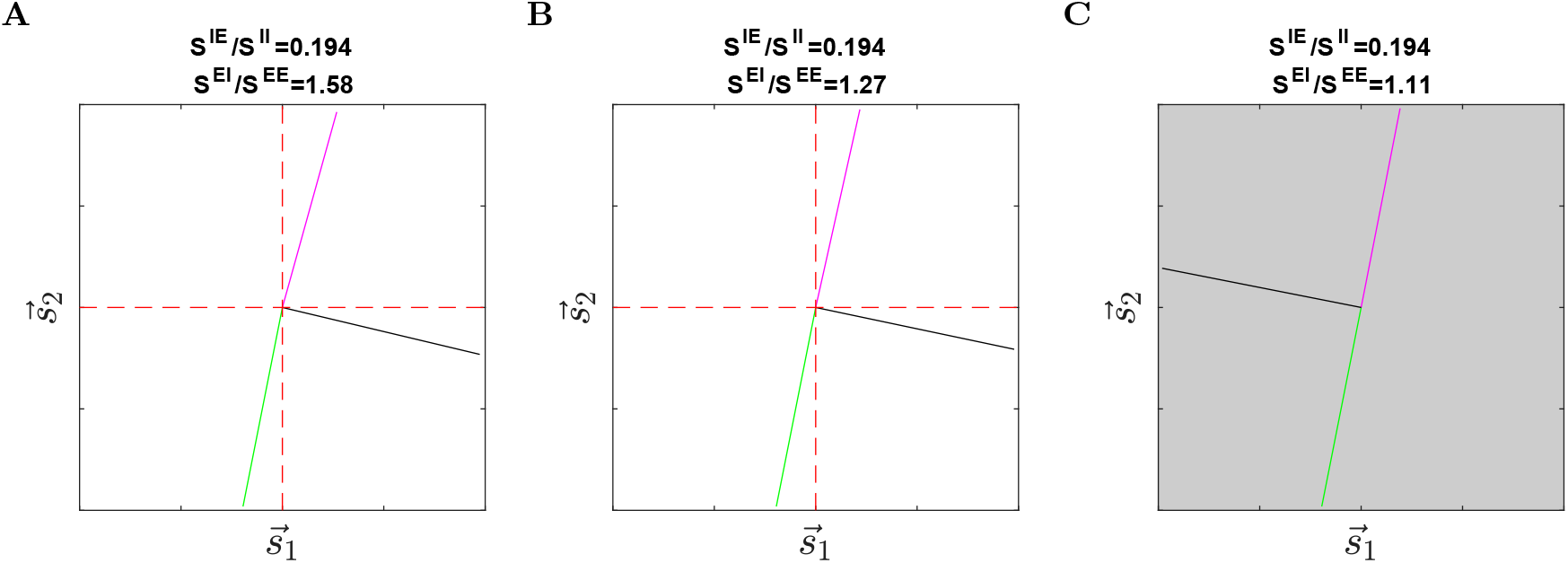
A geometric view of the solvability of MF equations. Each panel contains two normalized column vectors of I — **M**(**v**), i.e., ***α***_1_ (green vector in the third quadrant) and ***α***_2_ (purple vector in the first quadrant). The black line segment marks the region *S* = {*f*_1_***α***_1_ + *f*_2_***α***_2_ | *f*_1_, *f*_2_ ≥ 0}. **A.** Eq. (9) is nonsingular and well-behaved. **B.** Eq. (9) is near-singular and its column vectors are nearly collinear. **C.** det(*I* – **M**(**v**)) changes sign and MF+v gives negative firing rates.

To see what else might happen, we now move along a line in the inhibition plane defined by *S^IE^/S^II^* = 0.194, starting from the value *S^EI^/S^EE^* = 1.58 used in Fig. S1A and moving down. Fig. S1B shows what happens for *S^EI^/S^EE^* = 1.27, which lies within the good area: ***α***_1_ and ***α***_2_ become more nearly collinear, though there is still a sizable intersection between S and the first quadrant, so that most values of **s** lead to positive firing rates. However, as *S^EI^* decreases even further, det(*I* – **M**(**v**)) changes sign, and the set S abruptly flips to the other side of the dividing line, leading Eq. (9) to produce negative firing rates for many values of **s**.

In Fig. S2, we extend this picture to the inhibition plane. For each choice of *S^EI^*/*S^EE^* and *S^IE^*/*S^II^*, we compute **M**(**v**) by assuming **v** = (*v_E_, v_I_*) = (0.55,0.65). Each panel shows the two (normalized) column vectors ***α***_1_ and ***α***_2_, along with a black line marking the region *S*(***α***_1_, ***α***_2_). Observe that as *S^EI^*/*S^EE^* and *S^IE^*/*S^II^* decrease, ***α***_1_ and ***α***_2_ first become linearly dependent (so that *I* – **M**(**v**) becomes singular), then changing orientation and resulting in negative firing rate estimates (gray panels).

Two final remarks. First, the boundary between gray and white panels corresponds roughly to where det(*I* – **M**(**v**)) changes sign. This is also where explosive dynamics occur in our *network* simulations due to low suppression index SI_*E*_ or very low E-firing rates (caused by high external input to I-cells; more below). This suggests that parameters where the MF+v algorithm fails to give biologically meaningful estimates are also parameters where the network model itself fails to give biologically meaningful results.

Second, concerning the narrow wedge of **s**-values in the first quadrant that yield negative firing rate estimates, i.e., the set bounded by ***α***_2_ (oblique purple vector in the first quadrant) and the vertical axis in Fig. S1AB. The same pattern also occur in the upper part of Fig. S2: These correspond to when the external inputs to I-cells are too high. Though the corresponding sets of **s**-values are small, they can have a significant impact. For example, for high-*S*^*I*lgn^ and/or high-*F*^*I*L6^ regimes, this can lead to large swaths of gray. See Figs. 2 and 3, and **SI** (Sect. S5).

#### Implementation and design of MF+v

As explained in Methods (see M2), our first attempt at a fixed point iteration did not result in a convergent algorithm. So we iterate until the firing rate estimates stabilize to a narrow, nearly linear band in firing rate space, then average a number of successive estimates to produce an estimate.

Algorithm 1 gives a precise summary and lists all other hyperparameter values used. A practical issue is that we need to check the variance of the voltage and firing estimates to determine when to stop iterating. To make this efficient, instead of carrying out accurate but expensive long-time average for every iteration, we use shorter runs that may be noisy by themselves but can be averaged together to produce accurate estimates. We then use a small number of consecutive iterations to check convergence during an initial, “training” phase, and when certain stopping criteria are satisfied, we compute a more accurate estimate and output the result.

**Fig. S2:**
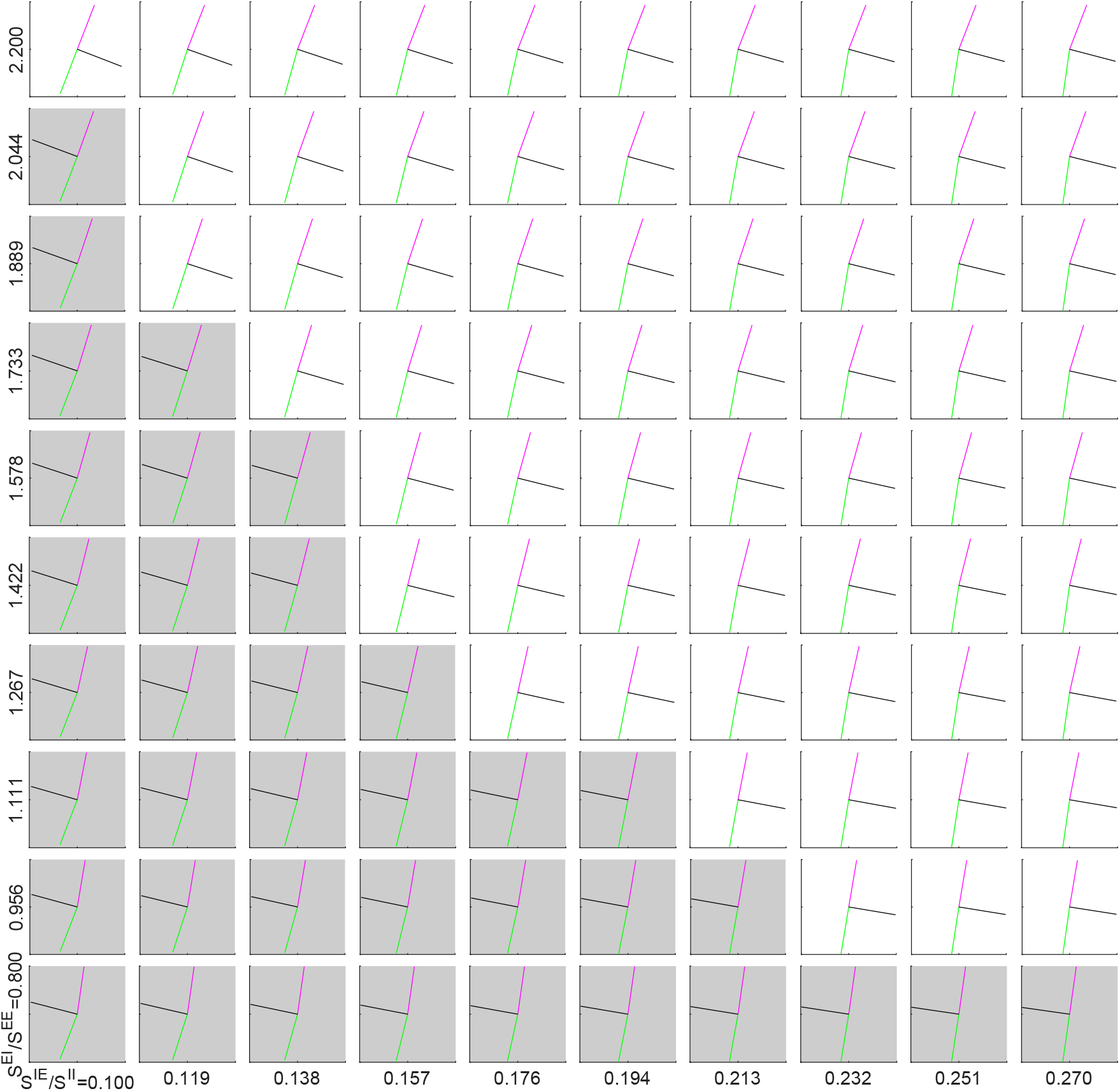
Solvability of Eq. (9) on the inhibition plane. Here, *S^EE^* = 0.024 and *S^II^* = 0.120, and we fix **v** = (*v_E_, v_I_*) = (0.55, 0.65) for all panels. Panels with gray color corresponds to negative firing rates from the MF equations, and the boundary between gray and white panels roughly corresponds to where det(*I* – **M**(**v**)) changes sign.

In this paper, the precise stopping criterion is based on comparing the *coefficient of variation*

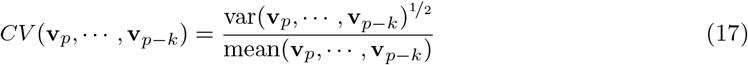

of the last *k* voltage estimates is against a pre-specified tolerance *ε*. When the stopping criterion is satisfied, we use moving averages of the voltages to compute a larger number of iterations, and use these to estimate the firing rate. For the network models studied in this paper, we have found the estimates to be insensitive to the exact choice of the maximum iteration number *M*. We typically set *M* in the range 300–500.

##### Algorithm 1: The MF+v method.

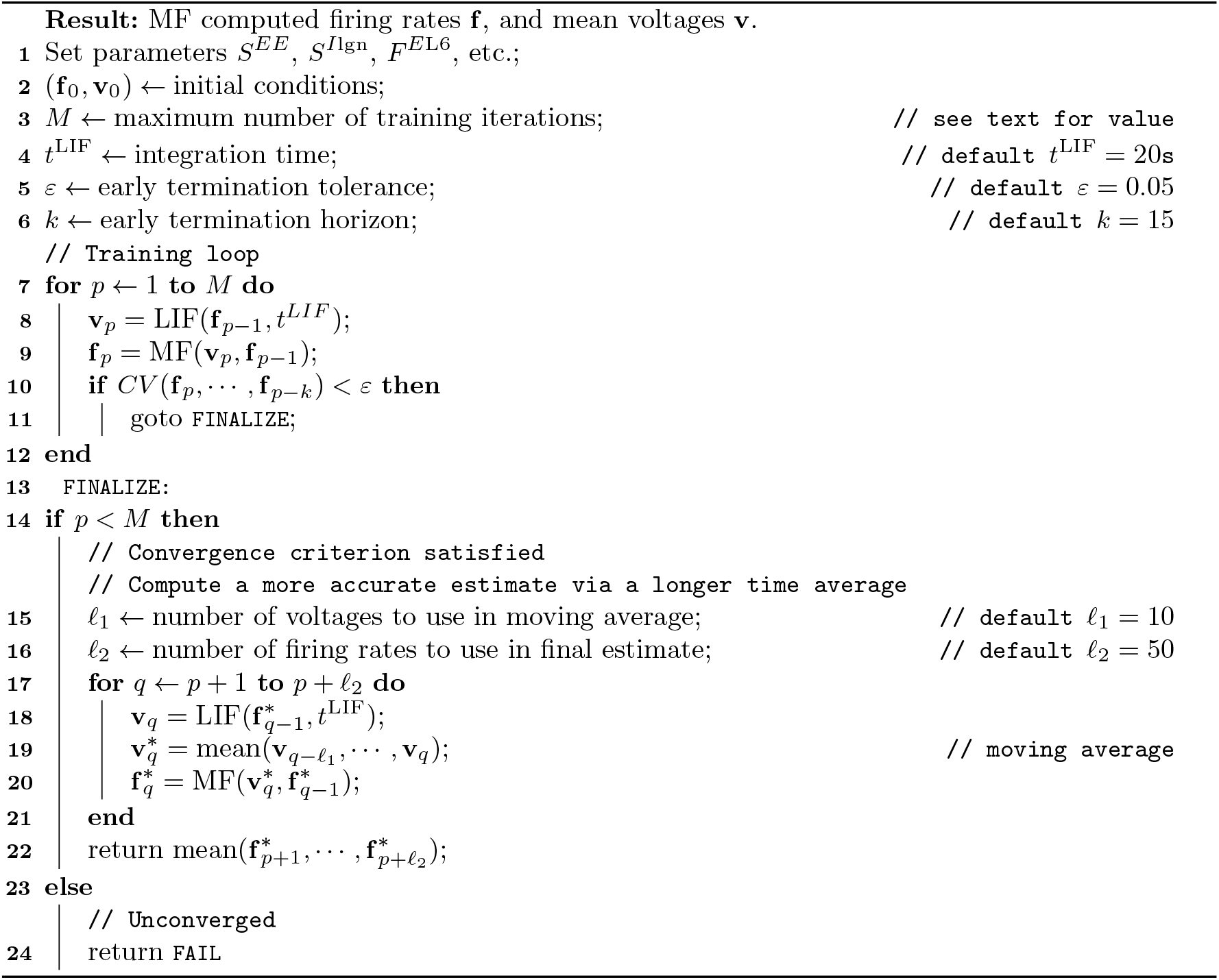

### S4 Miscellaneous information on MF+v

Here we record some additional information that have affected our decision to use MF+v in this paper.

#### Effects of refractory period and different kick sizes

The mean voltages **v** produced by the LIF equations (and hence the MF-computed firing rates **f**) can depend on parameters in a nontrivial way, making it difficult to estimate **v** using analytical methods. Here, we illustrate the parameter dependence of LIF neurons via two examples, using the parameters in Sect. S2. In both examples, a pair of LIF neurons (one E and one I) are each presented with Poissonian spike trains modeling L4 inputs, with input rates 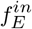 and 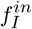, in addition to L6, LGN, and amb inputs. The two neurons are uncoupled and given independent inputs. We denote the resulting output rates 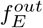 and 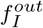.

Our first example concerns the refractory period *τ*_ref_, which can significantly impact neuronal activity because membrane conductances and currents steadily decay during refractory periods; the larger the *τ*_ref_, the more conductance is “missed” by the neuron while refractory. For instance, for an E-cell with a 3 Hz firing rate, its membrane potential stays unchanged for 3 × *τ*_ref_ in each second. This can be a non-negligible fraction of time, and the effect is exacerbated by higher firing rates. Fig. S3A) shows the mean voltages and firing rates as *τ*_ref_ varies from 0 to 4 ms. Though 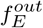 does not change much (5.5-5.8 Hz), 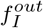 experiences a sharp change (19-25 Hz).

Our second example is motivated by the observation that for neurons in a mean-driven regime (a limit often studied in theoretical analyses of neuron models), their output rate depends on L4 kick sizes *S^QQ′^* (*Q, Q′* ∈ {*E, I*}) only through the product 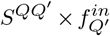. Thus, if we vary *S^QQ′^* while keeping 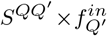 constant, any variation in output rates (**f**^*out*^) would be due to fluctuations in the L4 inputs. To test this, we perform the scaling 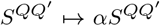 and 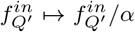 for a range of scaling factors *α*. Other parameters, including 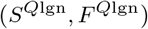 and (*S*^*Q*L6^, *F*^*Q*lgn^) (which were previously indexed to *S^QE^*) are kept constant. Fig. S3B shows the results. Observe that both **v** and **f**^*out*^ experience sharp changes when *α* moves away from 1. In particular, the firing rates are almost 0 when α is small (low fluctuation), and unreasonably high when α is large (high fluctuation). These results suggest that for the background regime studied here, MF+v (as is the network that it models) operates in a fluctuation driven regime.

**Fig. S3:**
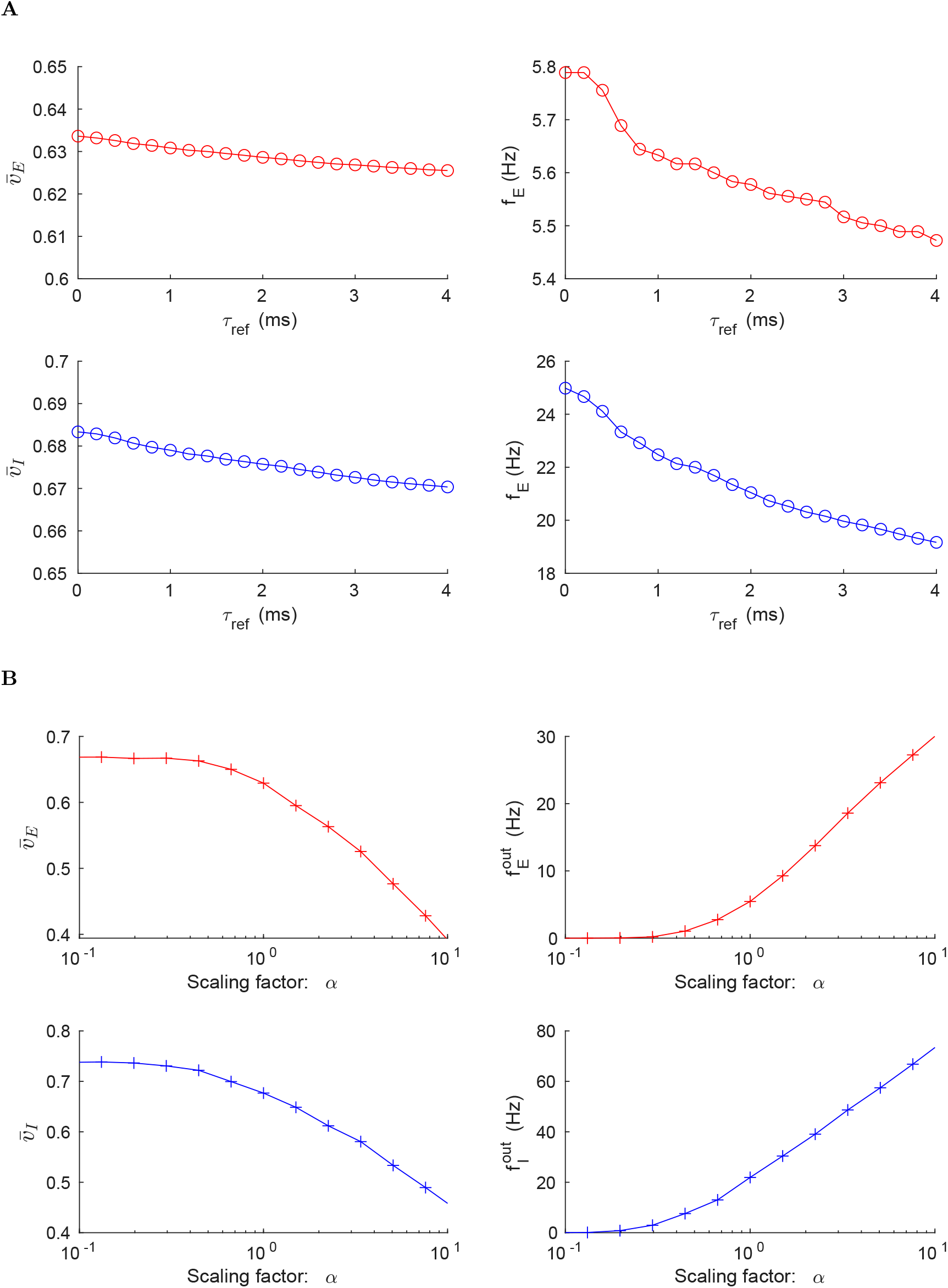
Mean voltages and firing rates computed by MF+v with different **A.** refractory periods (*τ*_ref_) and **B.** L4 excitatory kick sizes (*S^EE^* scaling factor *α*). Here, 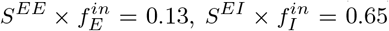, 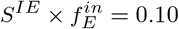, and 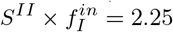.

**Fig. S4:**
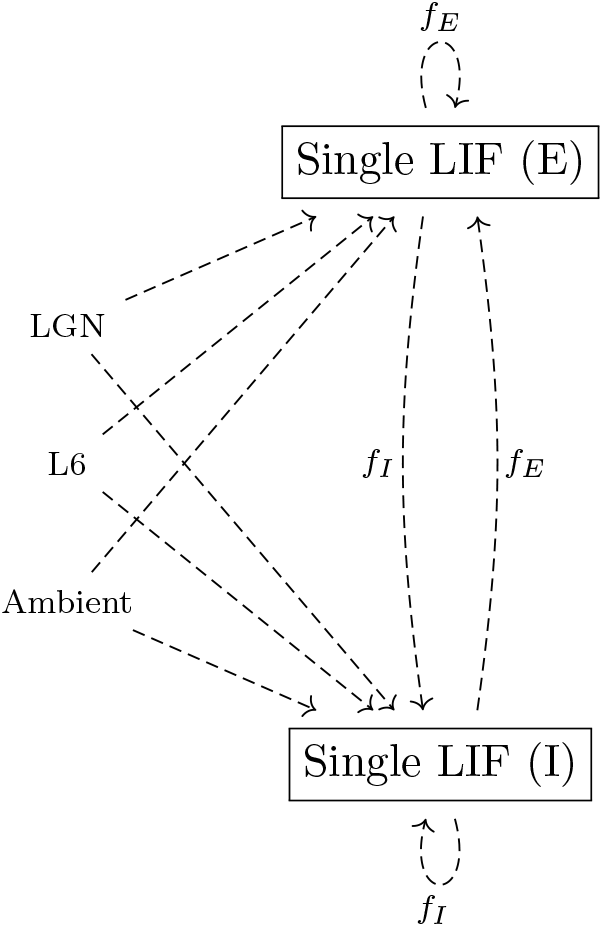
The LIF-only algorithm. We begin by choosing initial values 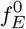 and 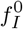. In the first iteration, these values are used to drive a pair of LIF neurons for 20 seconds (biological time). The resulting firing rates (instead of the membrane voltages in MF+v) are then fed into the next iteration as the L4 input. In the above, all dashed lines are modeled by Poisson processes.

#### Comparison to LIF-only

A natural alternative to MF+v is an LIF-only method: We drive a pair of LIF models (one E and one I) with Poissonian spike trains of rate **f** as L4 input, along with Poisson spike trains modeling inputs from LGN, L6, and amb, and look for values of **f** that lead to output rates equal to **f**. That is, we look for a self-consistent MF approximation without reference to Eq. (4). This can be implemented by simply iterating LIF neurons and feed 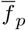 to iteration *p* + 1 directly as the L4 E/I input. A schematic representation is in Fig. S4, with details in Algorithm 2.

We find that all else being equal, the LIF-only algorithm is much less stable than MF+v. To demonstrate this, we select two parameters from Fig. 9C (*S^EI^* ∈ {0.0433, 0.0402}) and compare a simplified version of MF+v and LIF-only. In these runs, to avoid uncertainties associated with early termination, we train both algorithms for *M* = 100 iterations, then compute running averages. For MF+v, this means that in Algorithm 1, we set *M* = 100 and *ε* = 0, and always take the first branch after `FINALIZE`; see Algorithm 2 for LIF-only. Fig. S5 shows the results. In the left panels, we plot the firing rates **f**_*p*_ for iterates *p* ≤ 100 and running averages 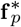 for 100 < *p* ≤ 400. As can be seen, firing rates from MF+v stabilizes quickly to network-computed rates, while LIF-only (right) sometimes exhibits large oscillations.

A potential explanation for the behavior of LIF-only is that (as we noted in Methods Sect. M2) one can obtain many more samples of voltages per unit time than spikes. Since the variance of firing rate estimates is roughly inversely proportional to the number of spikes, the single neuron rate estimates in the LIF-only algorithm are far noisier at typical background firing rates. We have also tested other variants of LIF-only, such as averaging over ensembles of pairs of LIF neurons. However, the LIF-only method remains rather unstable (data not shown). Although LIF-only occasionally gives good predictions of firing rates, it is far less reliable in comparison to MF+v.

##### Algorithm 2: The LIF-only algorithm. We have removed the early termination criterion to simplify comparison with MF+v.

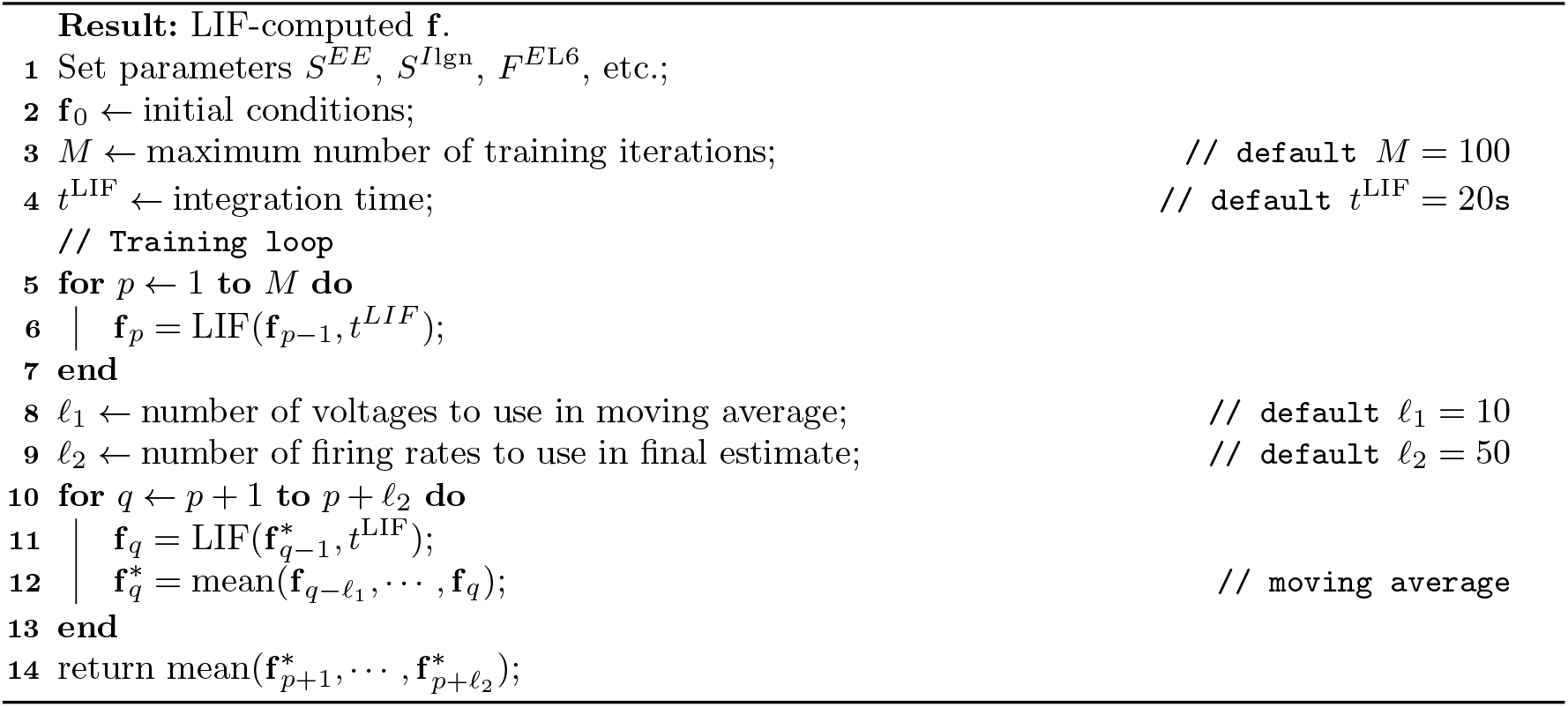

**Fig. S5:**
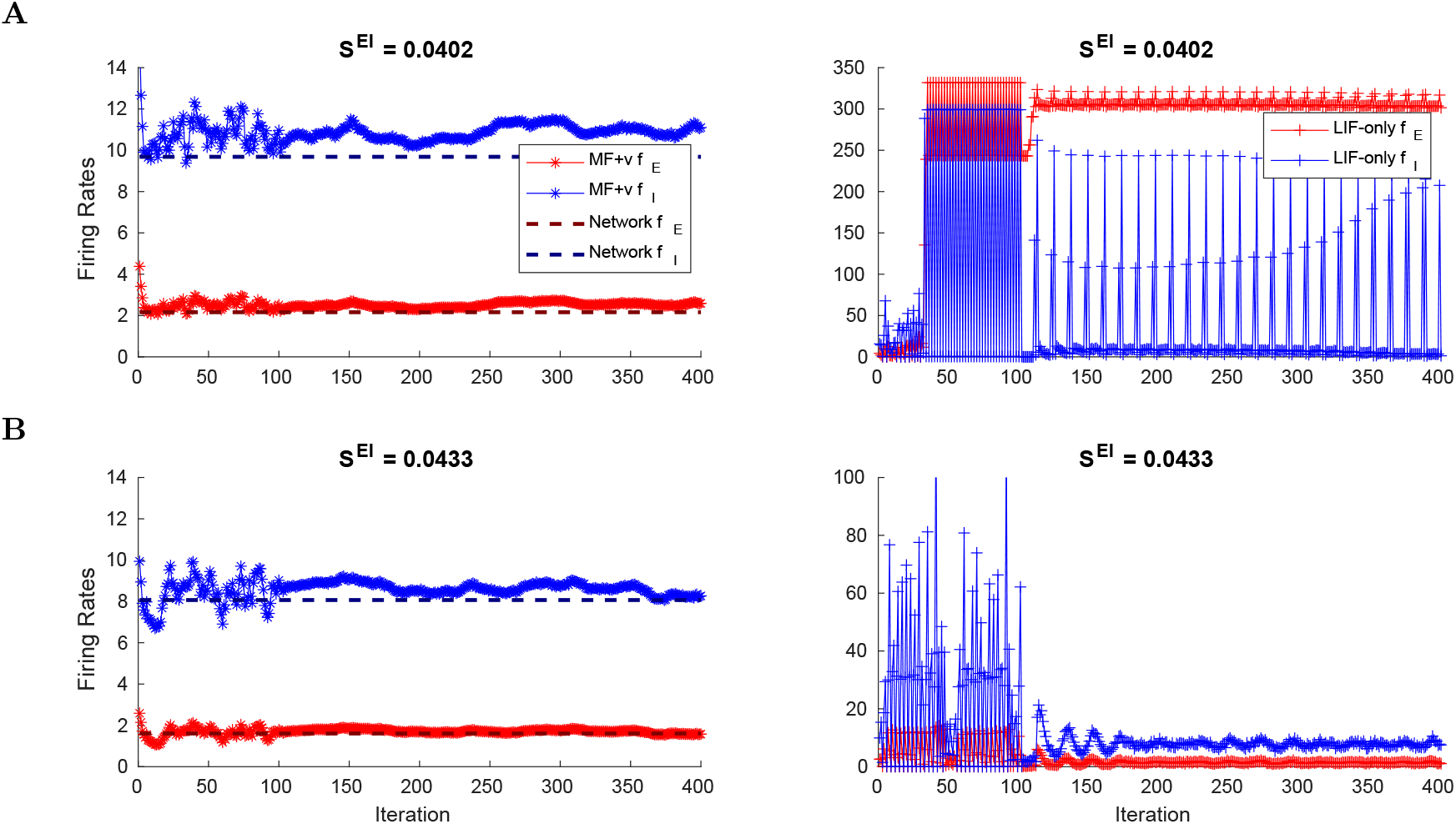
Comparison of MF+v and LIF-only. Both are trained for 100 iterations with identical LIF neuron simulation time *t*^LIF^ = 20s. **A.** E/I firing rate trajectories from **f**_*p*_ (iteration 0-100) and 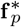 (iteration 101-400), for *S^EI^* = 0.0402. Left: MF+v exhibits stable estimates for both parameter choices. Right: LIF-only exhibit large oscillations. **B.** Same as **A**, but for *S^EI^* = 0.0433.

### S5 Additional firing rate maps

As mentioned in Results, here we show versions of Fig. 2 with different choices of parameters.

**Fig. S6:**
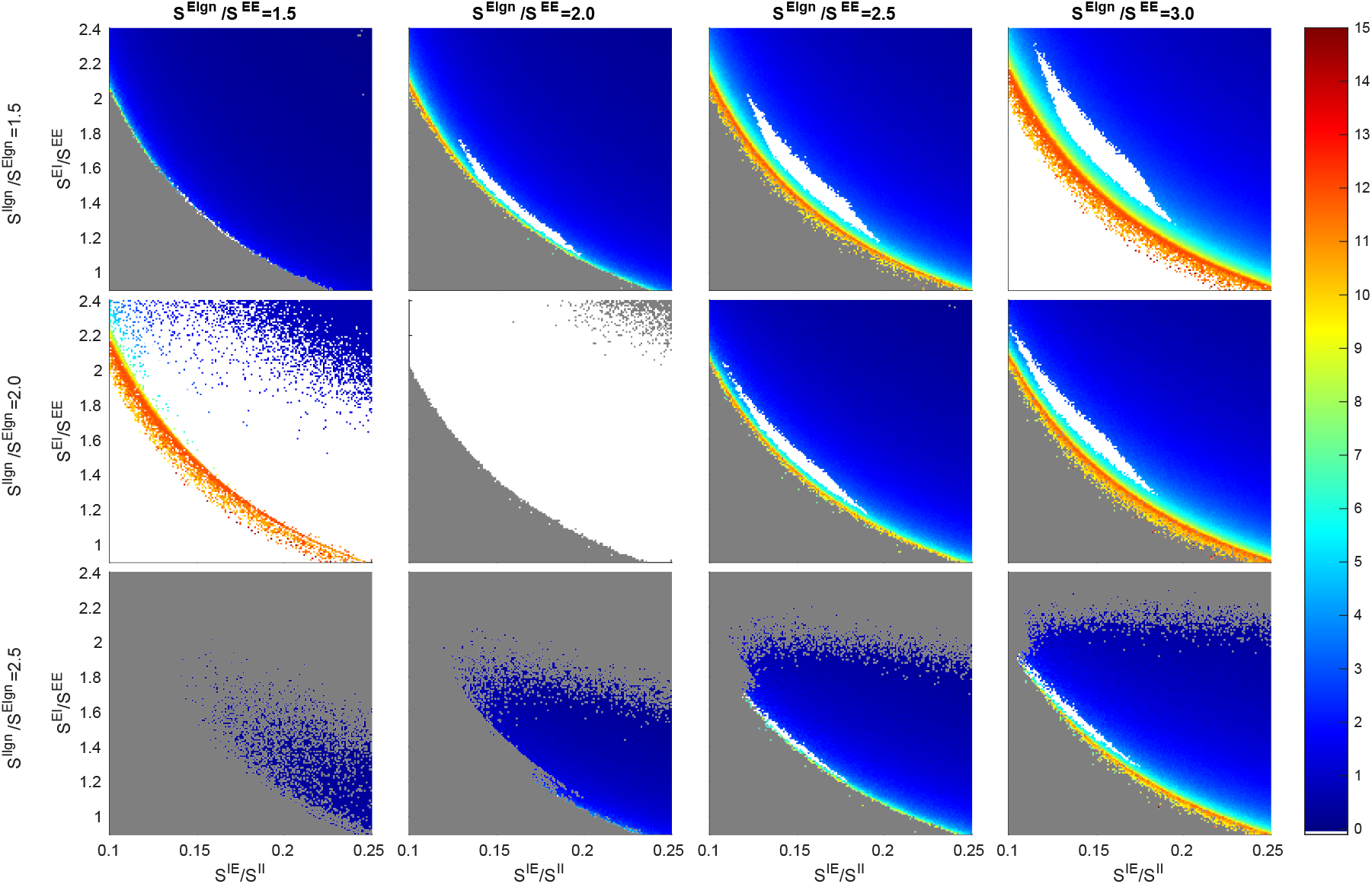
A version of Fig. 2 with *S^EE^* = 0:021, *S^II^* = 0:12, and *F*^*I*L6^/*F*^*E*L6^ = 3.

**Fig. S7:**
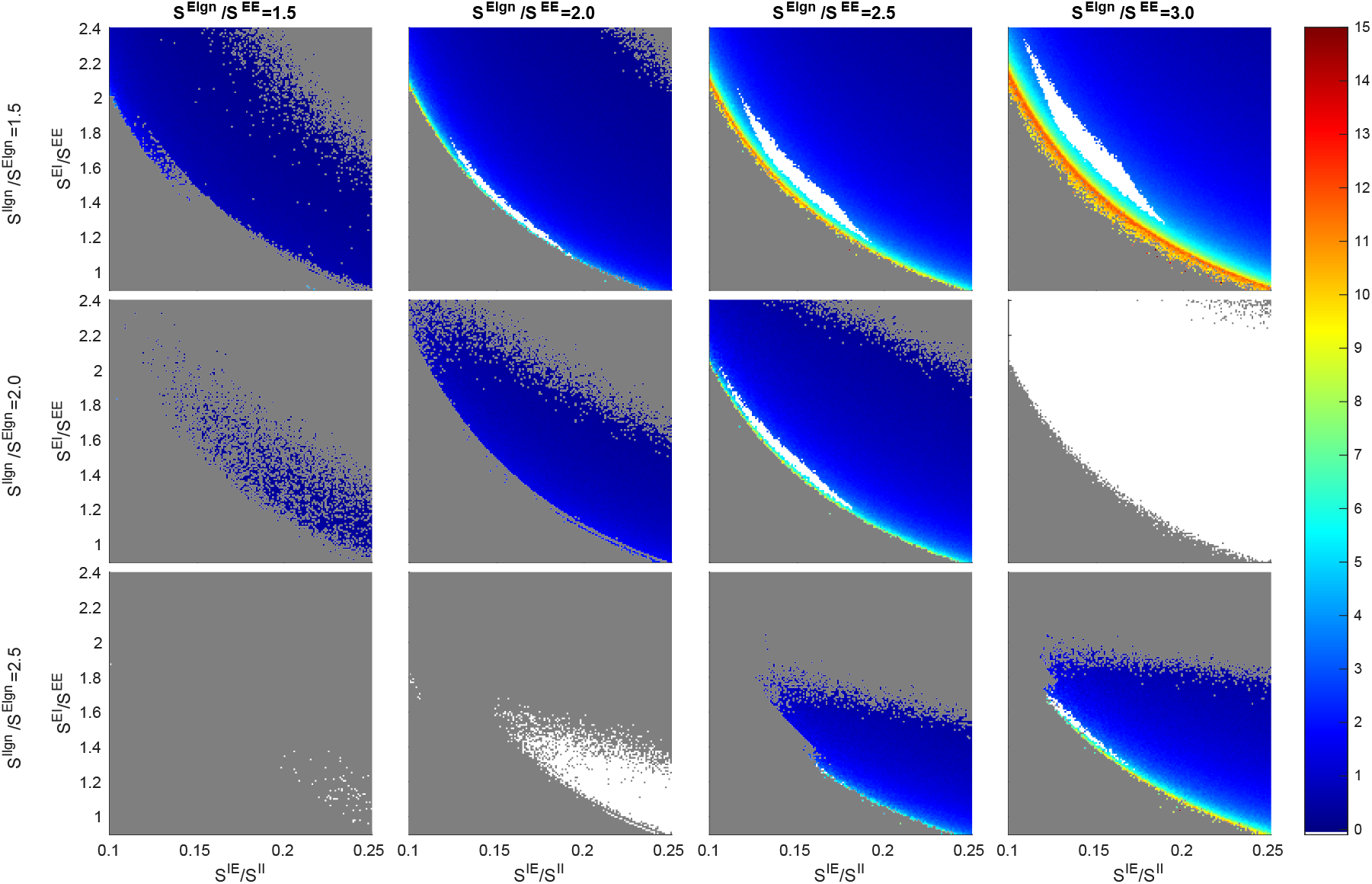
A version of Fig. 2 with *S^EE^* = 0.021, *S^II^* = 0.12, and *F*^*I*L6^/*F*^*E*L6^ = 4.5.

**Fig. S8:**
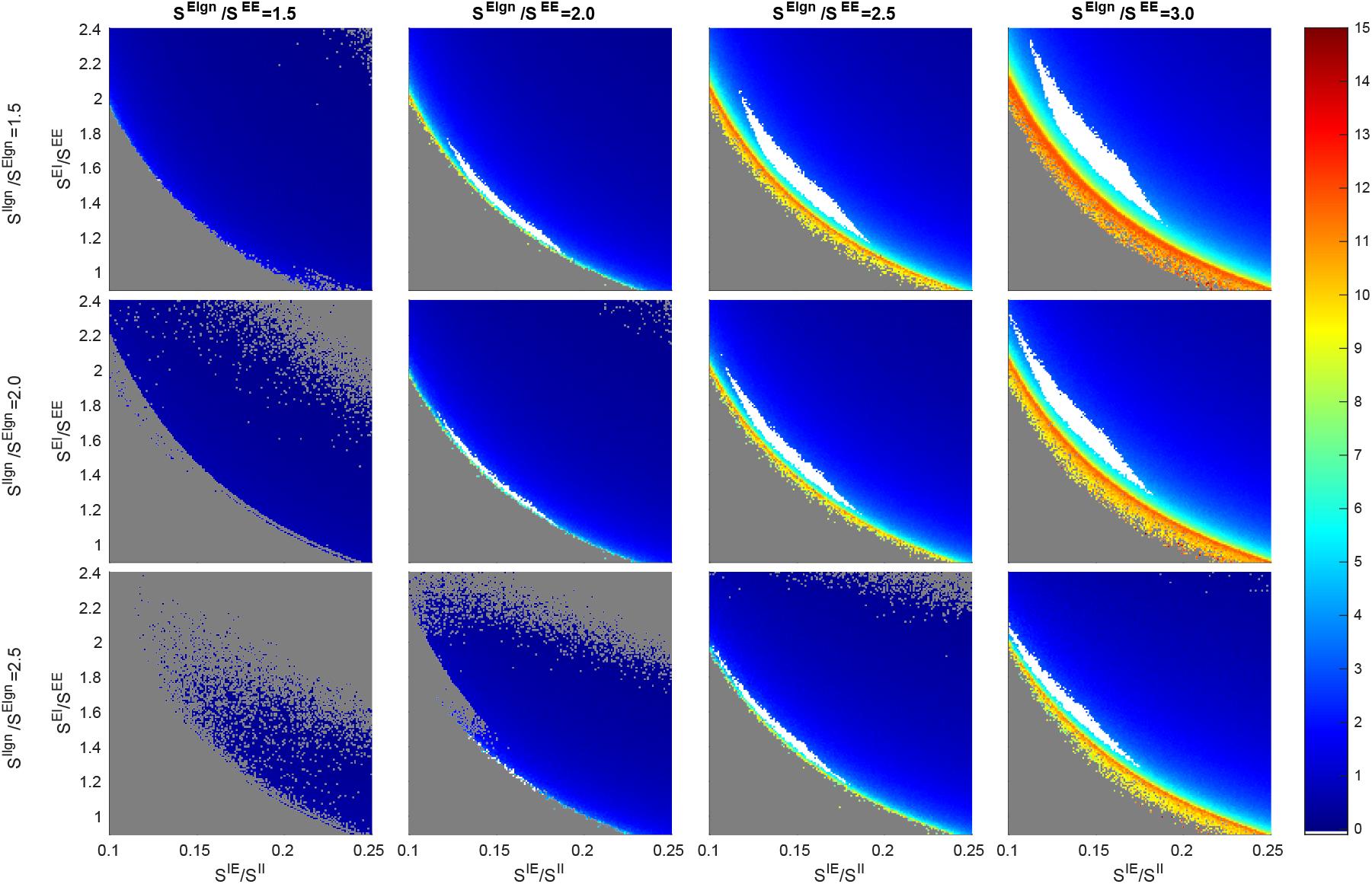
A version of Fig. 2 with *S^EE^* = 0.021, *S^II^* = 0.16, and *F*^*I*L6^/*F*^*E*L6^ = 3.

**Fig. S9:**
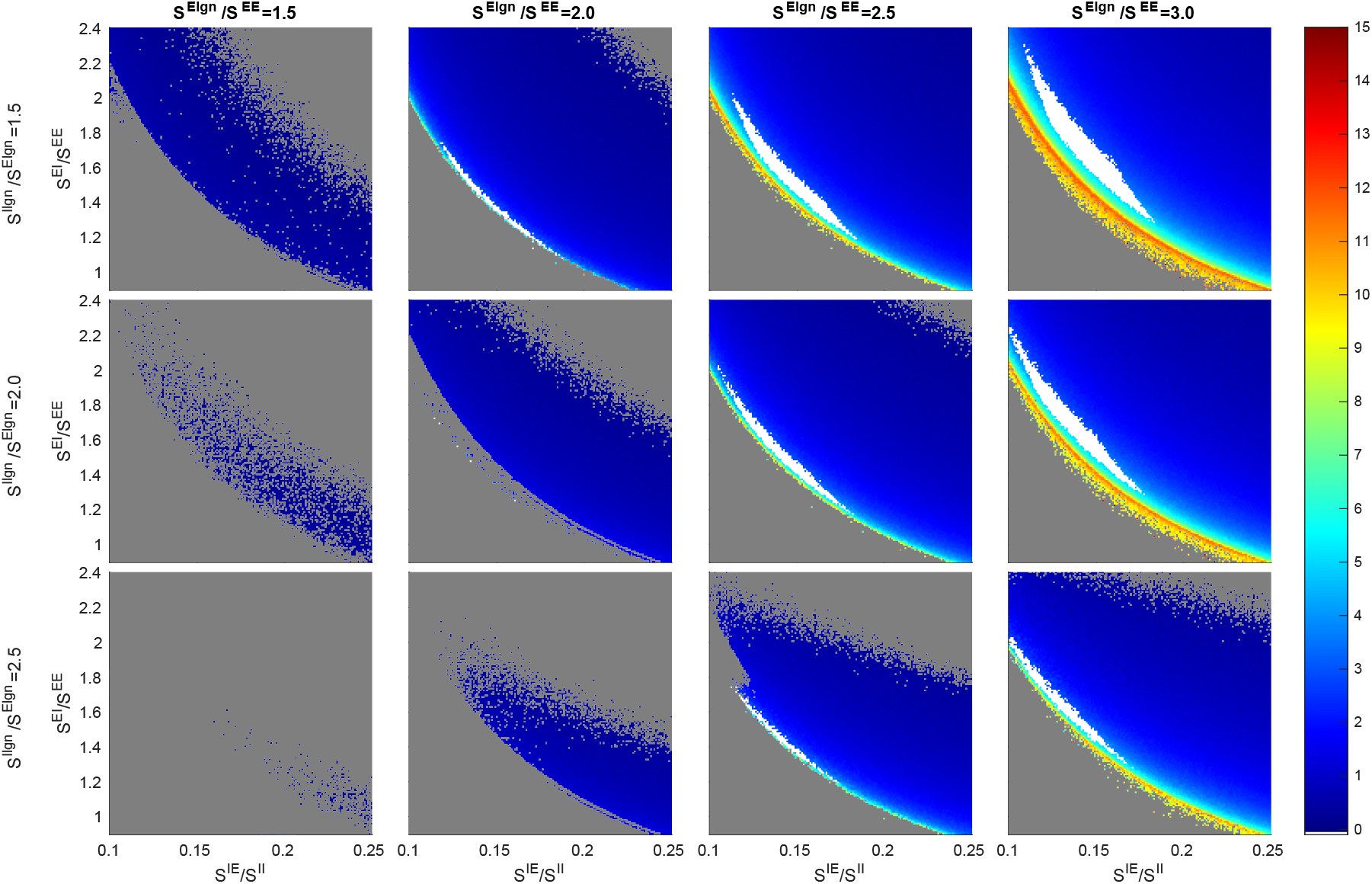
A version of Fig. 2 with *S^EE^* = 0.021, *S^II^* = 0.16, and *F*^*I*L6^/*F*^*E*L6^ = 4.5.

**Fig. S10:**
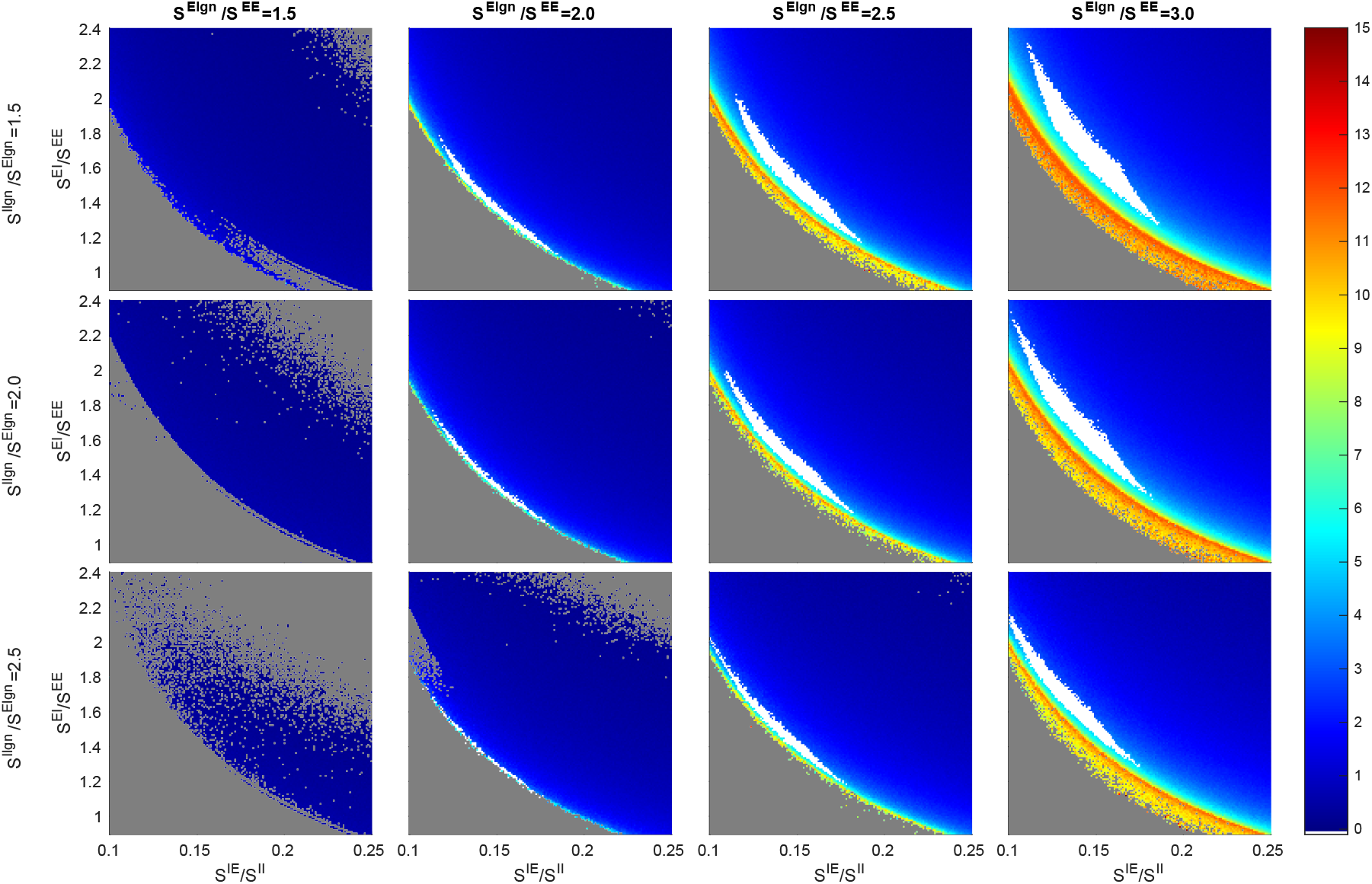
A version of Fig. 2 with *S^EE^* = 0.021, *S^II^* = 0.20, and *F*^*I*L6^/*F*^*E*L6^ = 3.

**Fig. S11:**
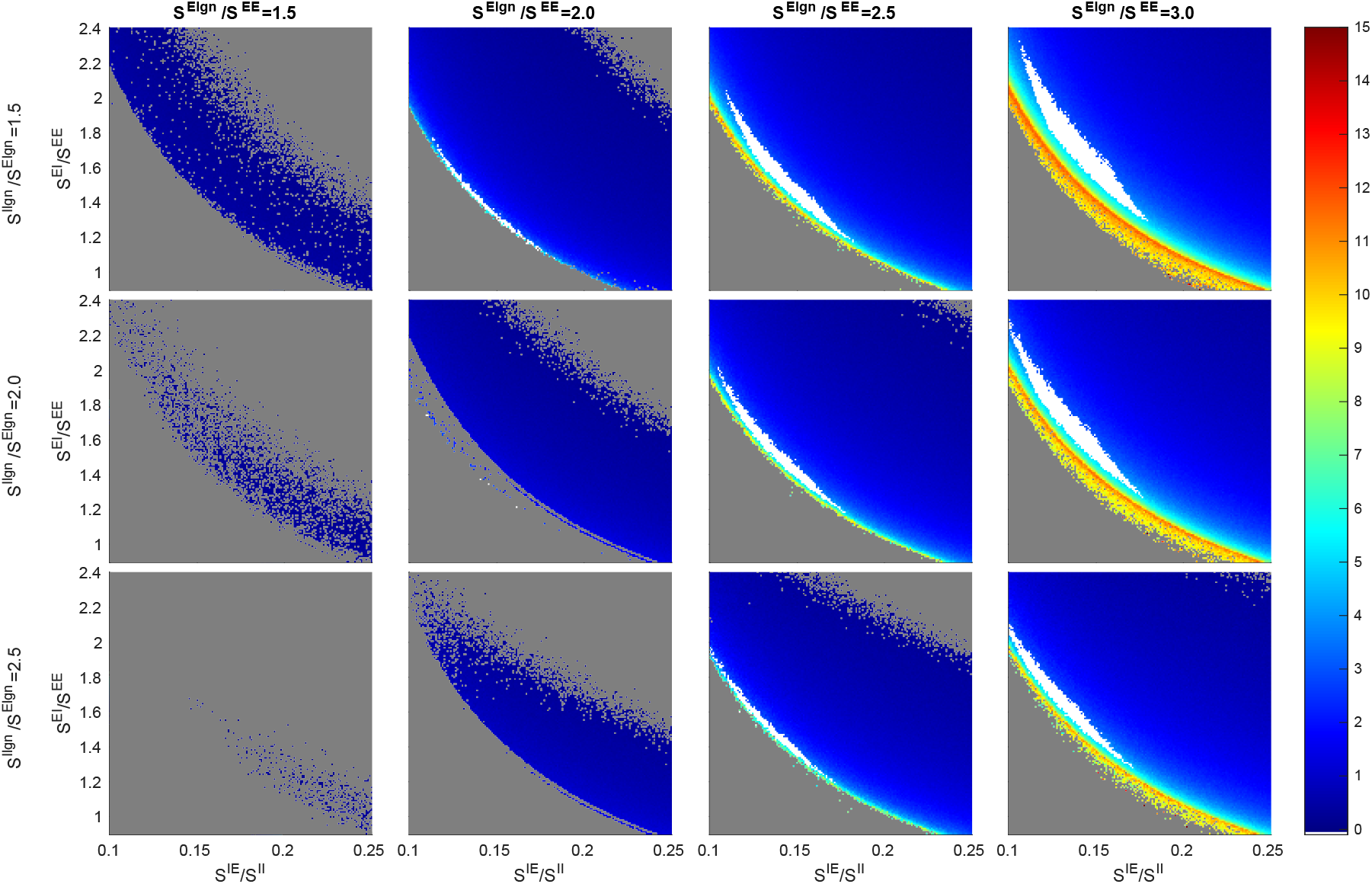
A version of Fig. 2 with *S^EE^* = 0.021, *S^II^* = 0.20, and *F*^*I*L6^/*F*^*E*L6^ = 4.5.

**Fig. S12:**
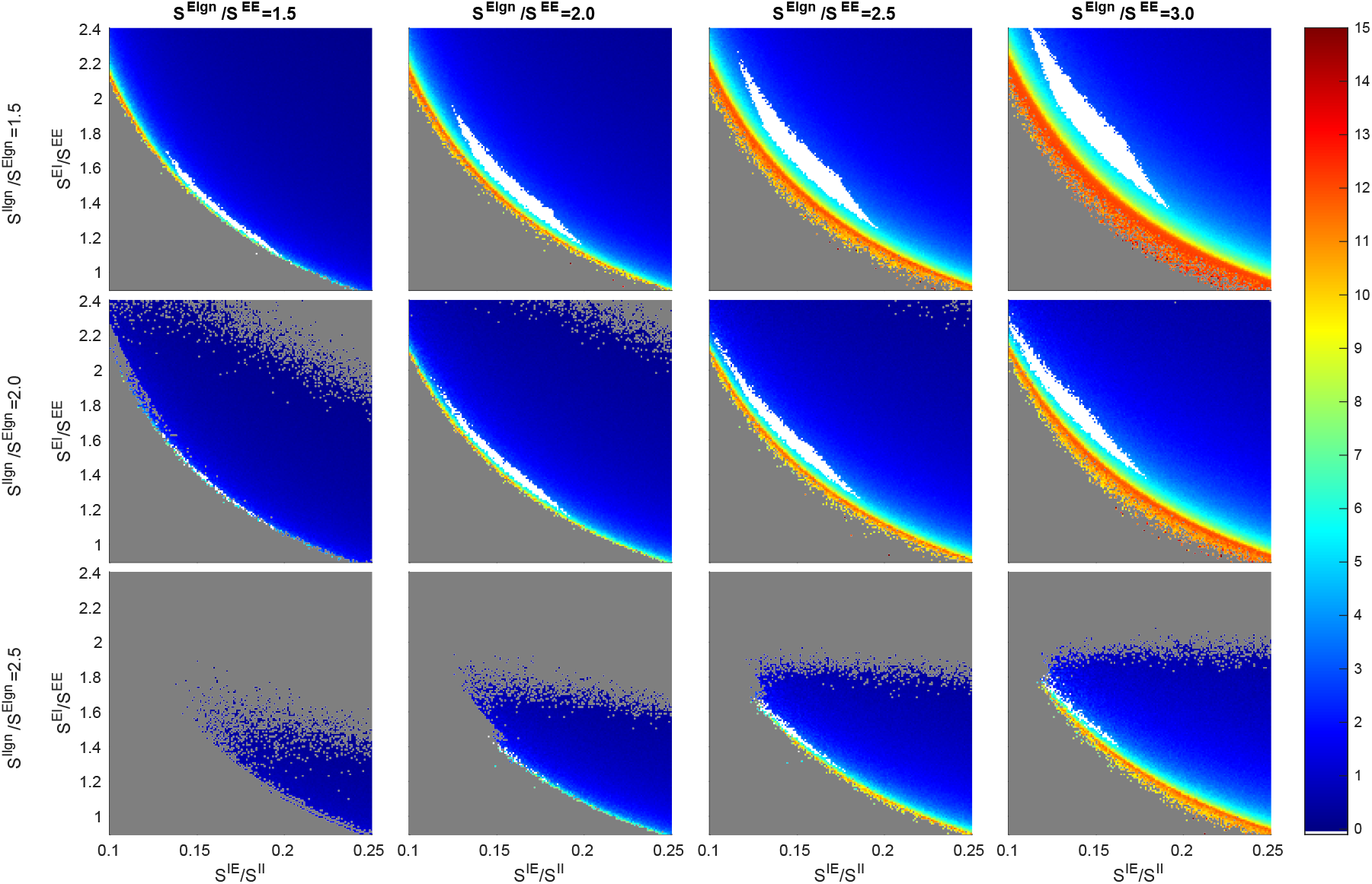
A version of Fig. 2 with *S^EE^* = 0.024, *S^II^* = 0.12, and *F*^*I*L6^/*F*^*E*L6^ = 3.

**Fig. S13:**
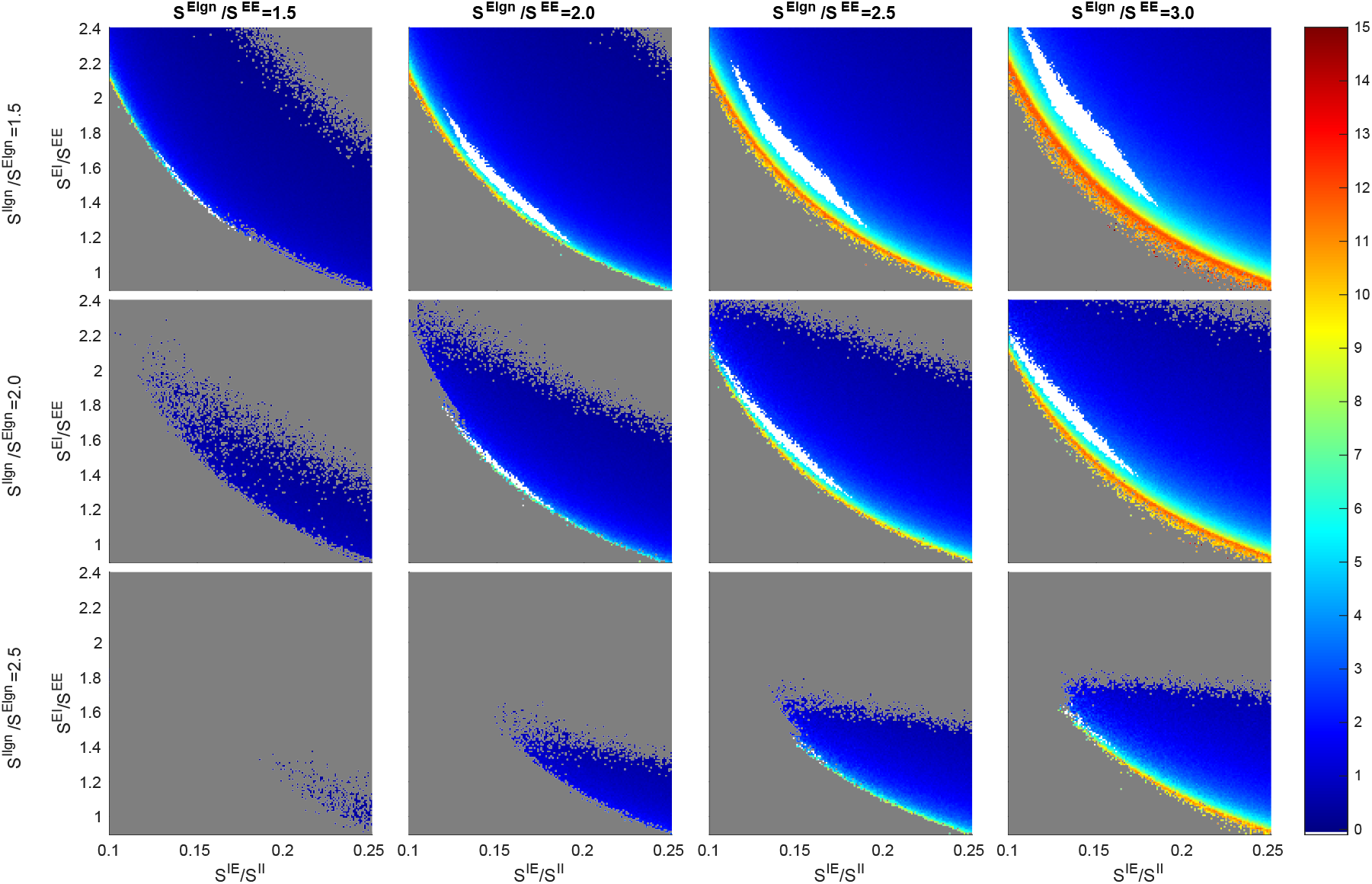
A version of Fig. 2 with *S^EE^* = 0.024, *S^II^* = 0.12, and *F*^*I*L6^/*F*^*E*L6^ = 4.5.

**Fig. S14:**
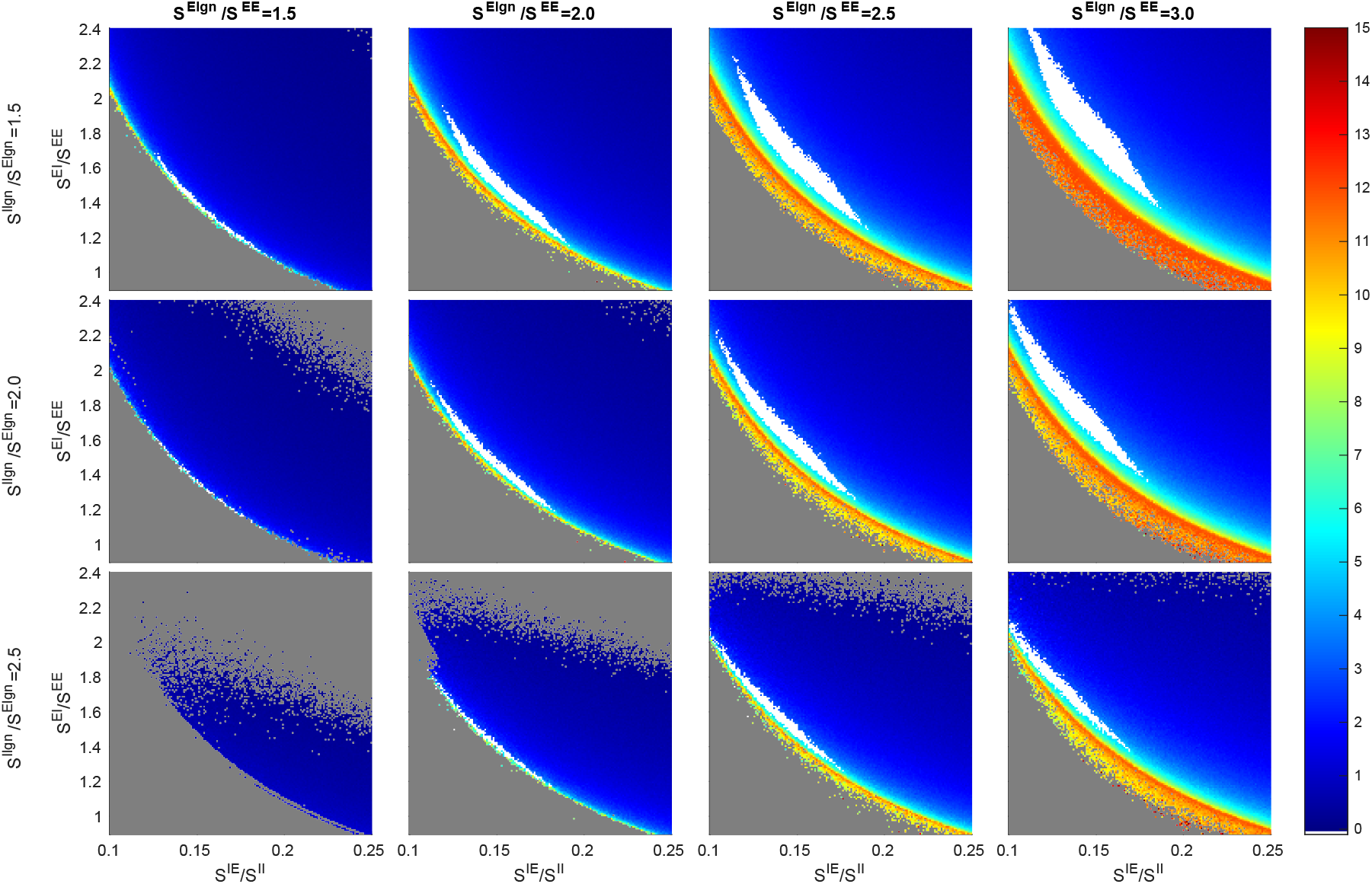
A version of Fig. 2 with *S^EE^* = 0.024, *S^II^* = 0.16, and *F*^*I*L6^/*F*^*E*L6^ = 3.

**Fig. S15:**
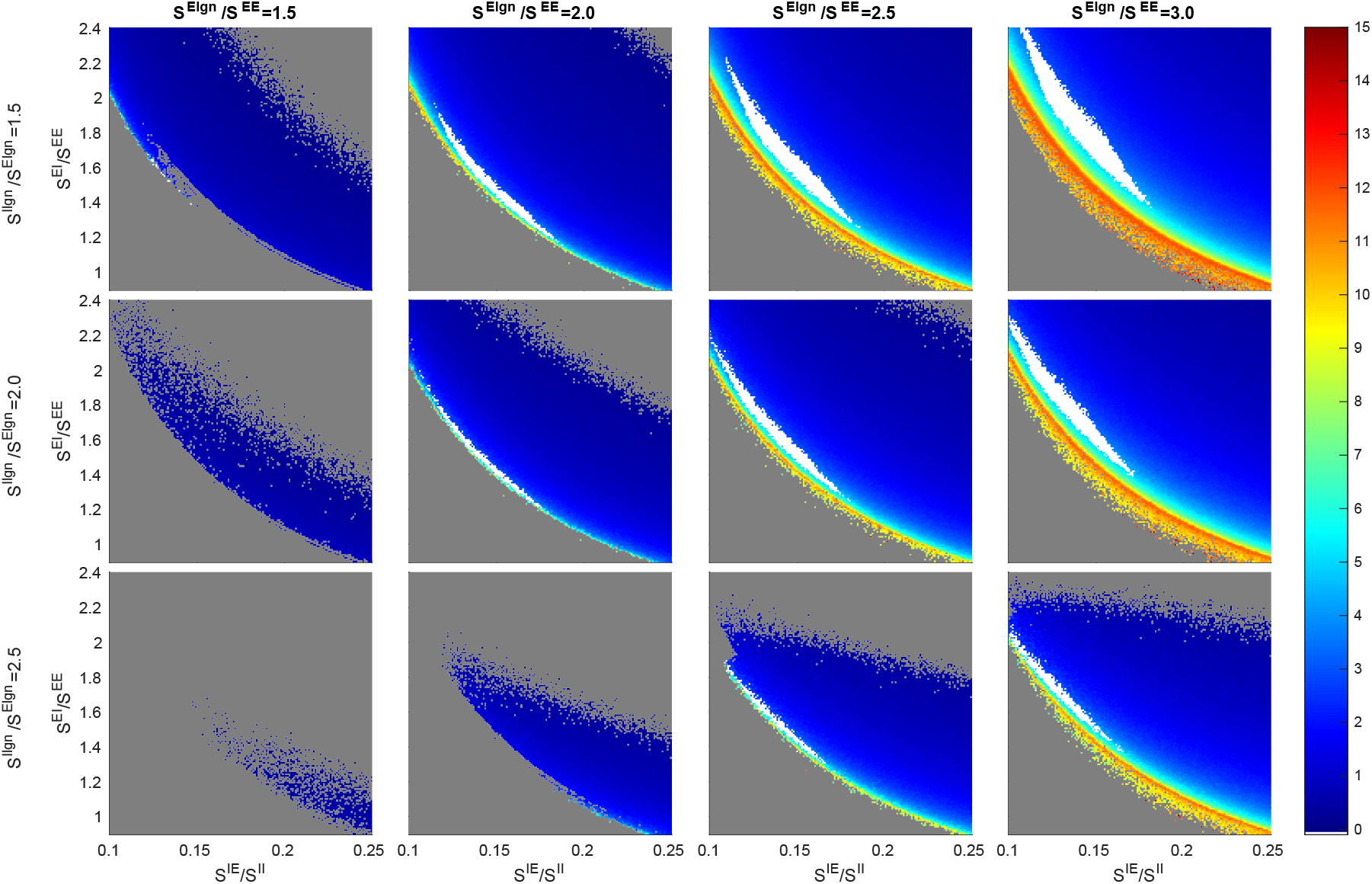
A version of Fig. 2 with *S^EE^* = 0.024, *S^II^* = 0.16, and *F*^*I*L6^/*F*^*E*L6^ = 4.5.

**Fig. S16:**
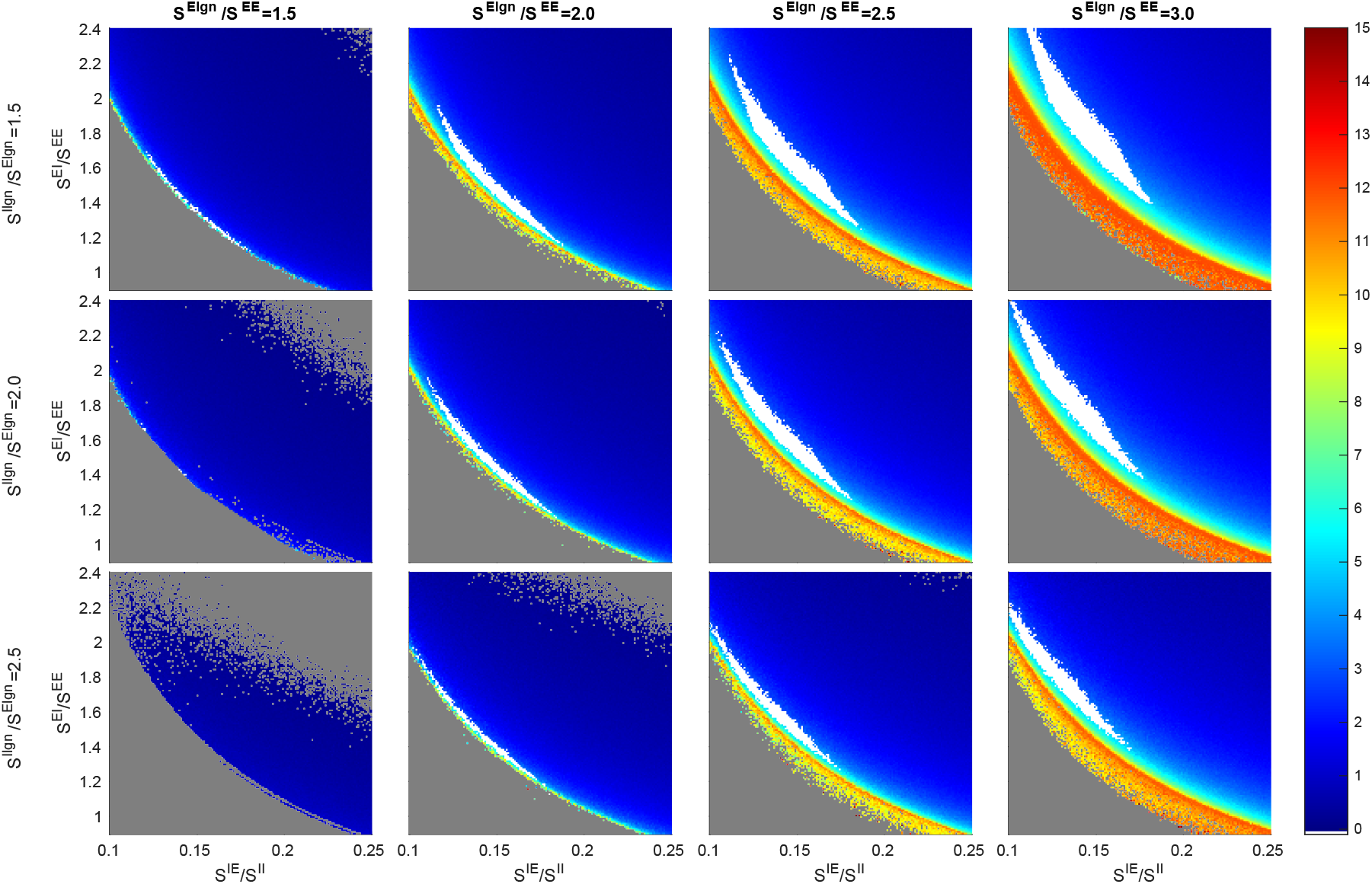
A version of Fig. 2 with *S^EE^* = 0.024, *S^II^* = 0.20, and *F*^*I*L6^/*F*^*E*L6^ = 3.

**Fig. S17:**
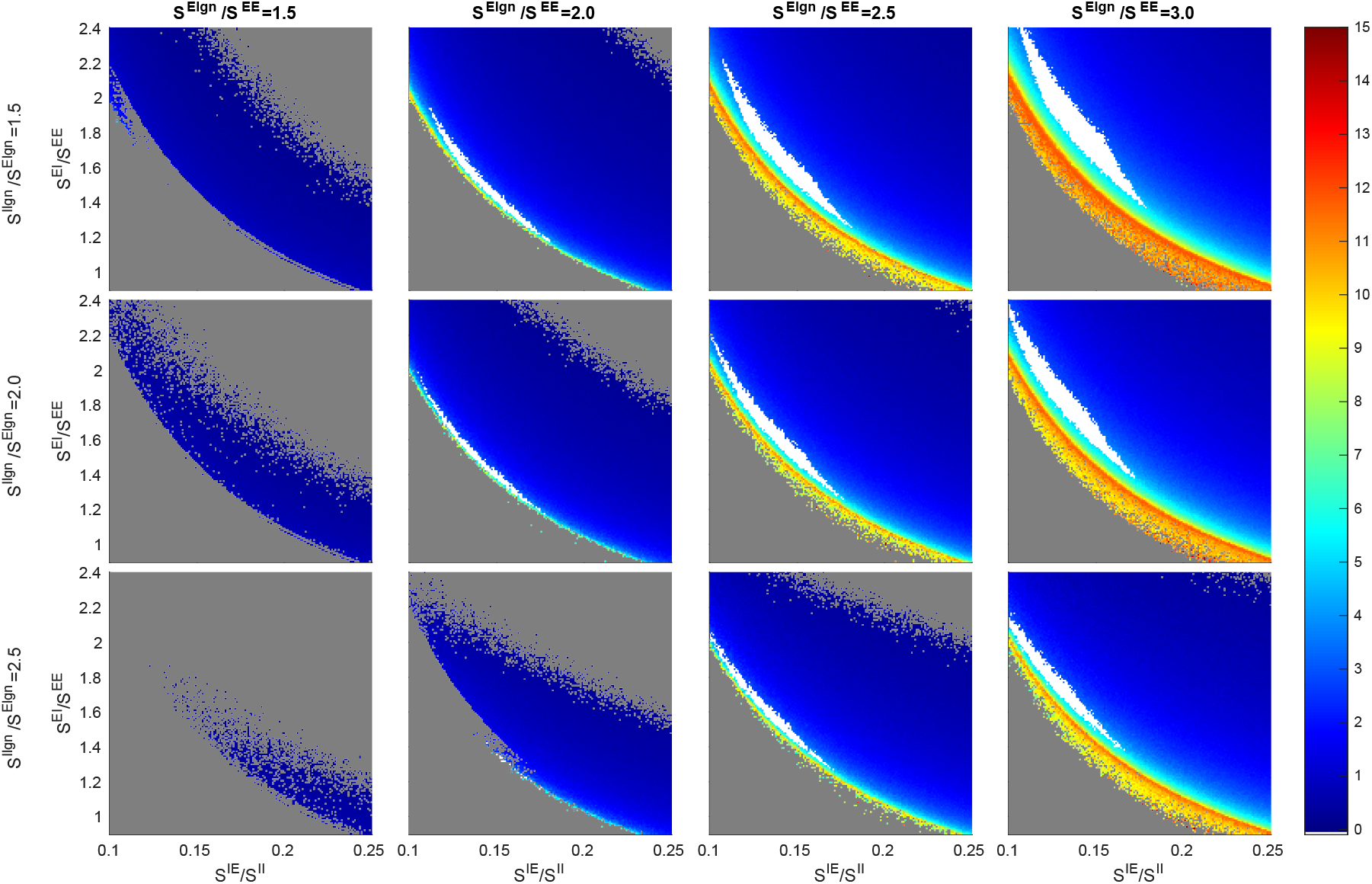
A version of Fig. 2 with *S^EE^* = 0.024, *S^II^* = 0.20, and *F*^*I*L6^/*F*^*E*L6^ = 4.5.

**Fig. S18:**
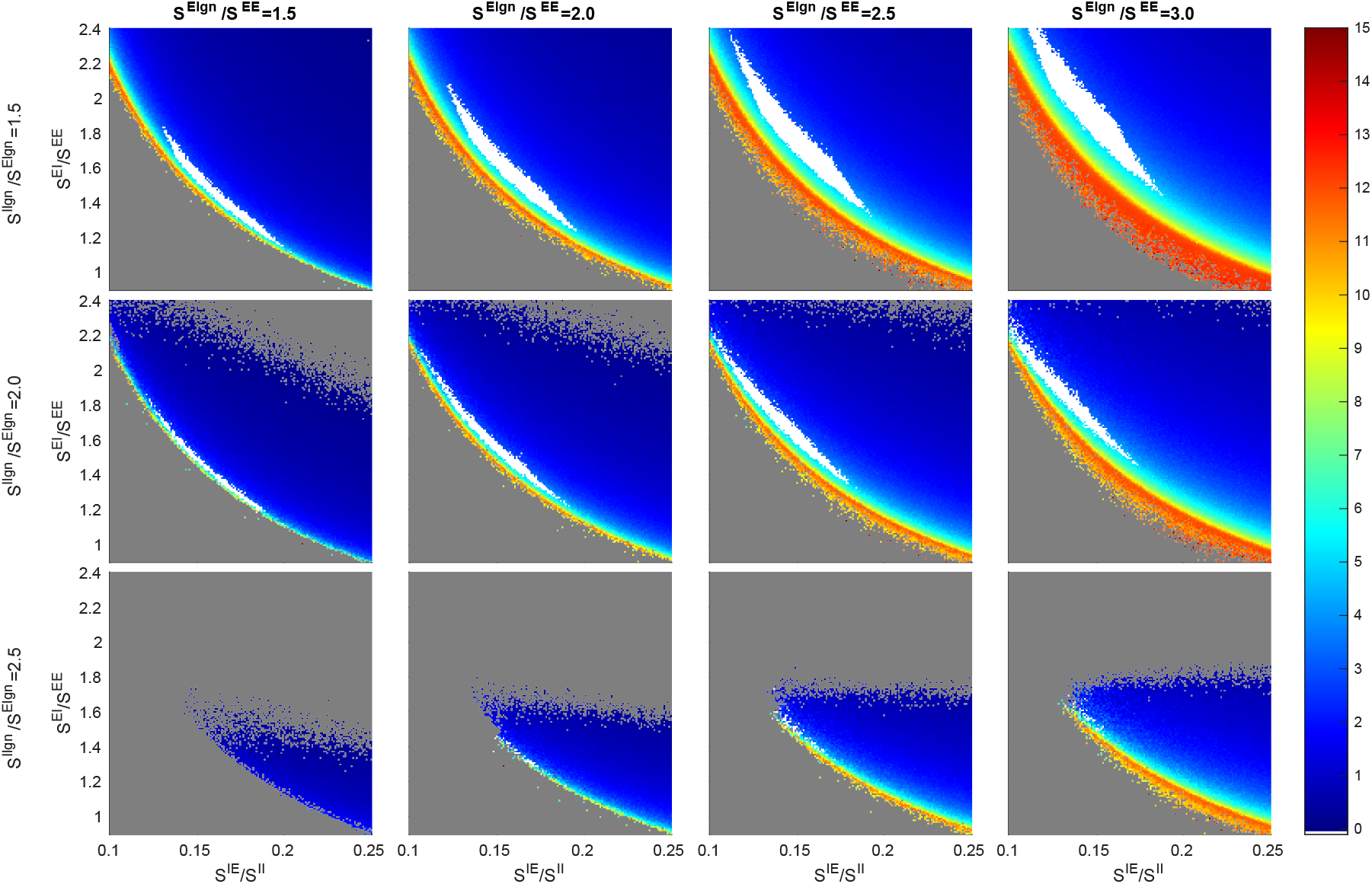
A version of Fig. 2 with *S^EE^* = 0.027, *S^II^* = 0.12, and *F*^*I*L6^/*F*^*E*L6^ = 3.

**Fig. S19:**
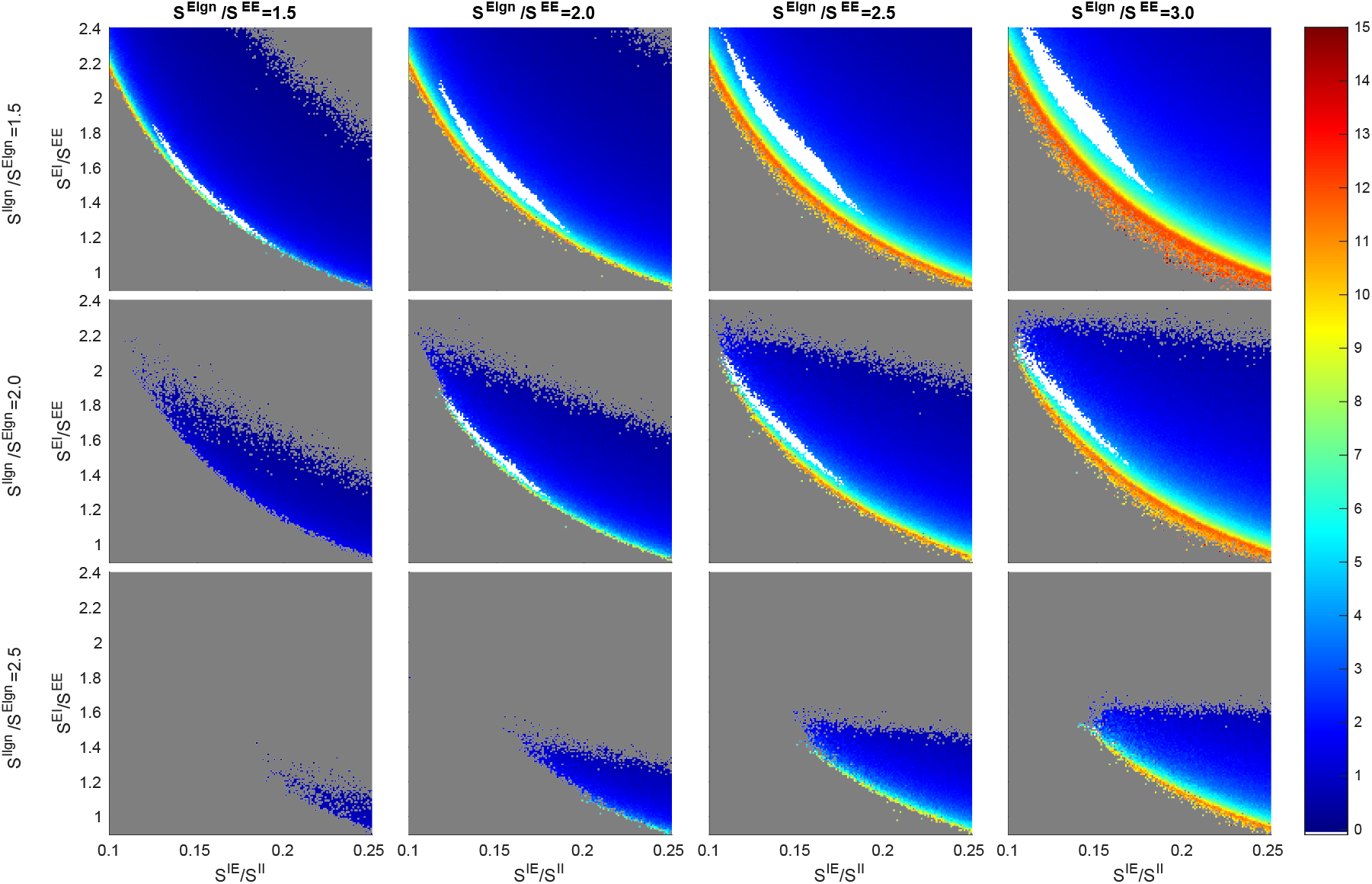
A version of Fig. 2 with *S^EE^* = 0.027, *S^II^* = 0.12, and *F*^*I*L6^/*F*^*E*L6^ = 4.5.

**Fig. S20:**
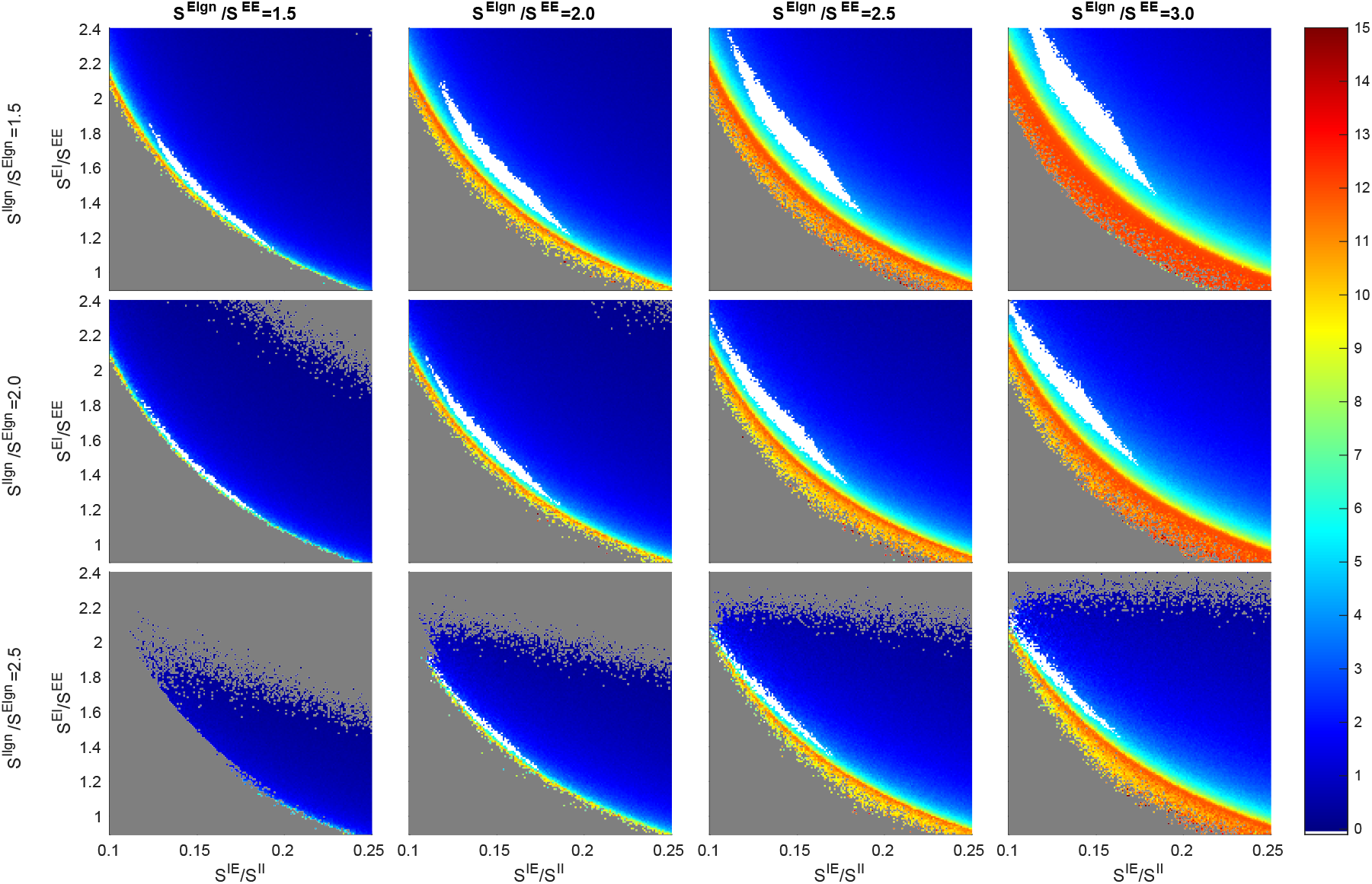
A version of Fig. 2 with *S^EE^* = 0.027, *S^II^* = 0.16, and *F*^*I*L6^/*F*^*E*L6^ = 3.

**Fig. S21:**
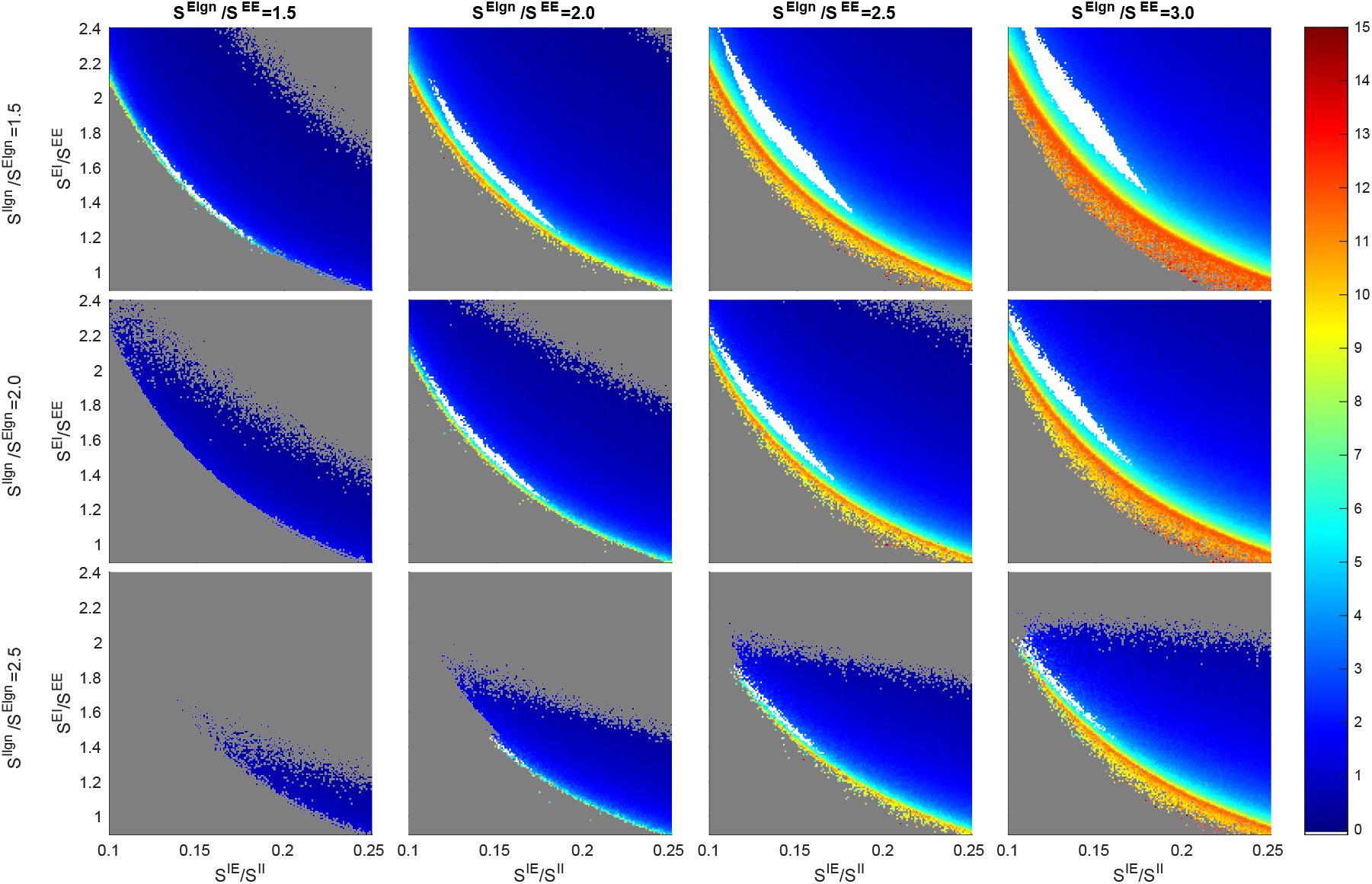
A version of Fig. 2 with *S^EE^* = 0.027, *S^II^* = 0.16, and *F*^*I*L6^/*F*^*E*L6^ = 4.5.

**Fig. S22:**
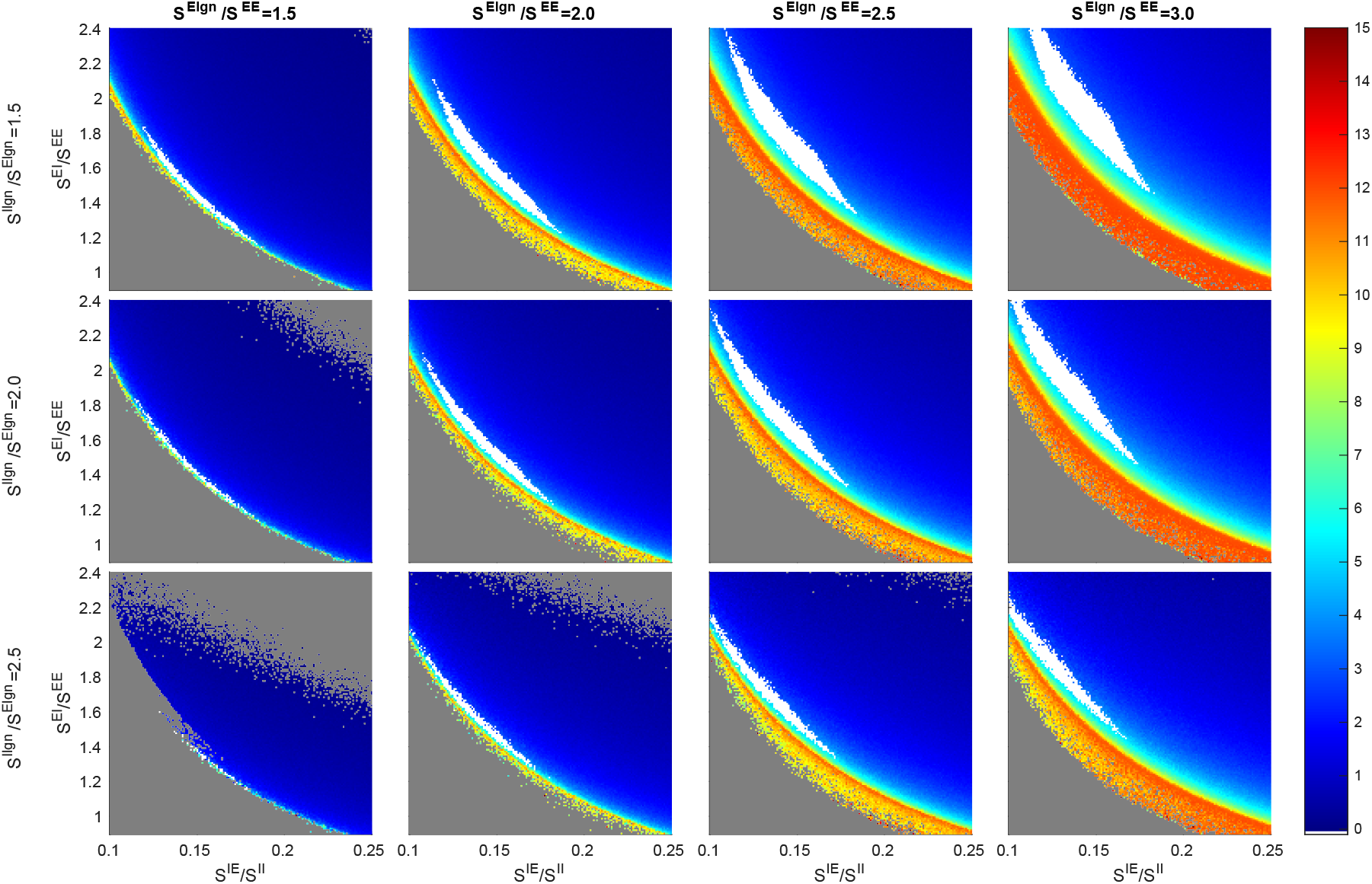
A version of Fig. 2 with *S^EE^ =* 0.027, *S^II^* = 0.20, and *F*^*I*L6^/*F*^*E*L6^ = 3.

**Fig. S23:**
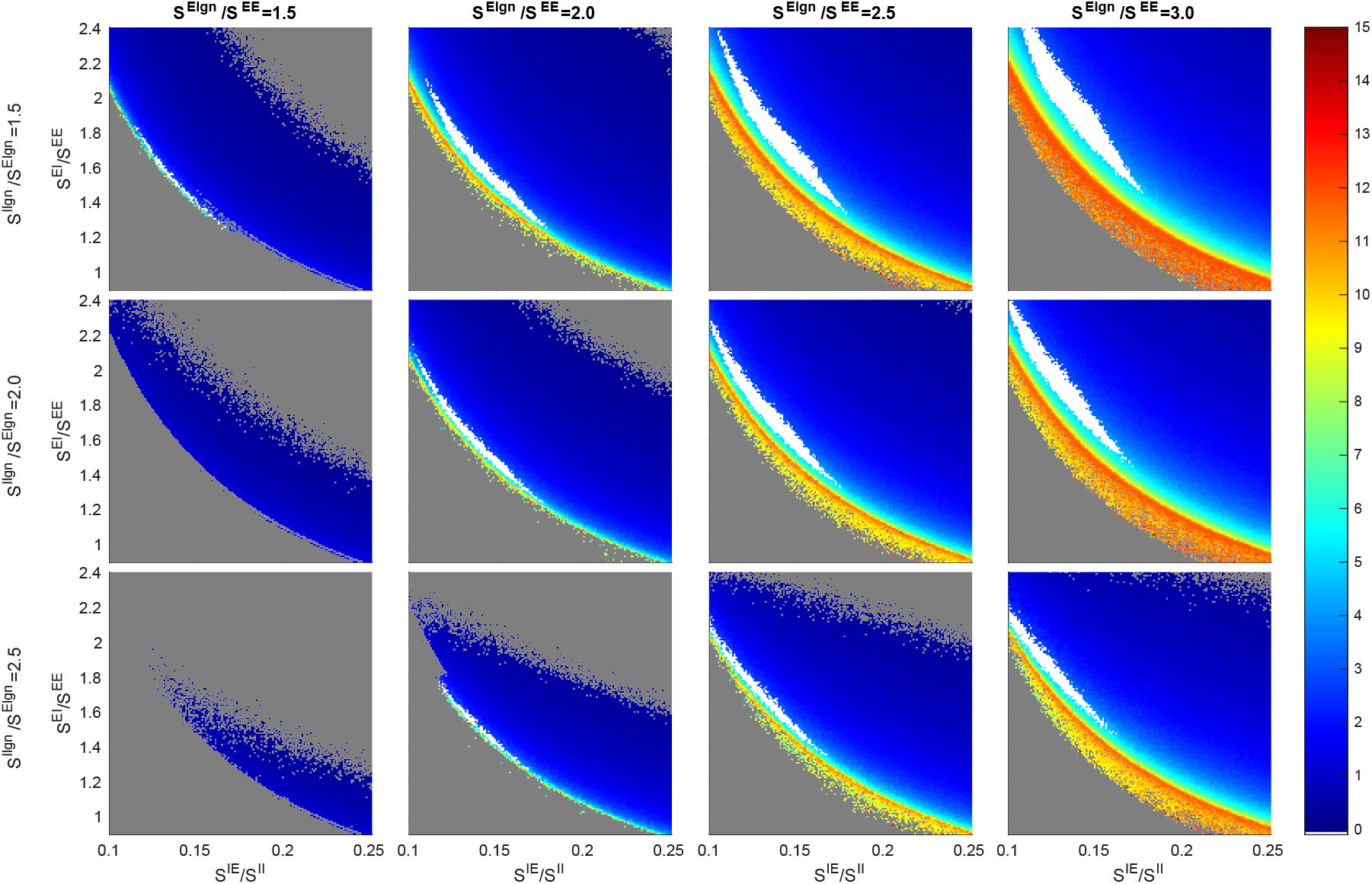
A version of Fig. 2 with *S^EE^* = 0.027, *^II^* = 0.20, and *F*^*I*L6^/*F*^*E*L6^ = 4.5.

## Footnotes

1 In this paper, “cells” and “neurons” both refer to nerve cells in the primary visual cortex.

2 Both network simulations and MF+v computations are implemented using MATLAB^®^ 2020B with Intel Xeon Platinum 8268 24Core 2.9GHz processors.

